# Herbivore prevalence poorly predicts yield in diverse cropping systems

**DOI:** 10.1101/2024.09.06.611601

**Authors:** Luuk Croijmans, Daan Mertens, Dirk F. van Apeldoorn, Yufei Jia, Nelson Ríos Hernández, Erik H. Poelman

## Abstract

Industrialized agriculture needs sustainable alternatives to pesticides to avoid negative impacts on the environment and human health. Crop diversification is known to decrease pest pressure in agricultural crops. Up till now, effects of insect herbivores on crop yield are often assumed equal among cropping systems. Here, we show that cropping system alters the effect that herbivores have on cabbage crop yield, where more herbivores do not necessarily lead to reduced yields. Our most diverse cropping system had simultaneously the highest number of herbivores and highest crop yield. Conversely, in a cultivar mixture we observed negative impacts of herbivores on yield. Our study shows that, in addition to the time of arrival and type of herbivore, cropping system should be considered when assessing how insect herbivores affect crop yield. We emphasize how our perception of herbivorous insects as pests is flawed and limits conservation efforts and sustainable farming practices.

## Introduction

Current high-input industrialized farming practices are highly productive and thereby contribute to the growing food demand. However, they reduce soil health, biodiversity and pollute (natural) ecosystems (Beaumelle et al., 2023; Wagner et al., 2021). Crop diversification can sustainably maintain crop yield and quality by stimulating ecological processes that replace inputs like chemical fertilizers and pesticides (Rakotomalala et al., 2023; Tamburini et al., 2020). The ecological processes that concertedly determine yield, like crop growth and herbivore pressure, exhibit complex interactions that are highly context dependent. Integrated pest management (IPM) is one of the pillars of the EU Farm to Fork strategy to ensure sustainable food production (European Commission, 2020), and crop diversification is suggested as an important IPM tool to prevent pest outbreaks (Barzman et al., 2015). Therefore, greater understanding of how crop diversification affects the ecological interactions between herbivorous insects and crops will ultimately contribute to designing sustainable cropping systems that enhance pest control.

Diversified cropping systems can alter herbivore abundances, crop growth, and consequently crop yield and quality. Appropriately designed, diversified cropping systems facilitate beneficial tri-trophic interactions, improve soil health, and enhance crop yield and quality (Beillouin et al., 2021; Carrillo- Reche et al., 2023). Crop rotations enhance soil quality, diversify below-ground (microbial) communities, and break pest, weed and disease cycles (Kremen & Miles, 2012). Reducing fertilizer use, shifting to organic alternatives, and including nitrogen-fixing crops in the cropping system alter resource availability (Jensen et al., 2010; Viketoft et al., 2021), which increase crop resilience to pests and diseases (Aguilera et al., 2021; Han et al., 2022). Intercropping of structurally dissimilar crops and including semi-natural habitats, like beetle banks and flower strips, enhance natural enemy abundance and performance (Beillouin et al., 2021; Langellotto & Denno, 2004). Intercropping systems reduce apparency or reachability of host plants for herbivores, as neighboring crops serve as a physical or chemical barrier (Mansion-Vaquié et al., 2020). Allowing genotypic variation within crops increases resistance to herbivory (Reiss & Drinkwater, 2018; Wetzel et al., 2018), and selecting for crop genotypes with complementary traits further increases crop resistance to herbivores and diseases (Barot et al., 2017). Thus, cropping system choices in time (e.g. rotations, relay cropping), space (e.g. crop configurations, field sizes), genes (i.e. composition of crop species, cultivars, varieties) (Ditzler et al., 2021) and management (e.g. tillage, pesticide and fertilizer use) affect crop growth, herbivore abundances and crop production in a myriad of ways. However, if and how cropping systems alter how herbivores affect crop yield and quality has rarely been studied, and the effect on yield per herbivore is usually regarded to be similar among cropping systems.

The impact of herbivorous insect pests occurs over the lifetime of crop plants and often involves species-rich communities. Herbivores may affect yield directly by causing crop injury. The effect of herbivores on yield via direct consumption depends on the timing of injury, the capacity of the plant to defend itself and on plant-mediated interactions with other herbivores. Plant growth-defense trade-offs vary with plant ontogeny and determine the direct impact of herbivory on yield (Boege & Marquis, 2005; Züst & Agrawal, 2017). In addition to such direct consumptive interactions, herbivores may indirectly affect crop yield by altering the crop-insect community over the course of the growing season via plant-mediated (multi-trophic) interactions (Mertens et al., 2021; Ohgushi, 2005; Van Zandt & Agrawal, 2004). Herbivore prevalence is affected by plant-mediated legacies of previous herbivore presence, which affects the colonization of crops by new herbivores through e.g. changes in plant apparency or the occupation and depletion of specific niches (Ohgushi, 2005; Van Zandt & Agrawal, 2004). Furthermore, herbivory can lead to enhanced plant defenses throughout the growing season and to the establishment of beneficial tri-trophic interactions by attracting natural enemies of current and future herbivores (Poelman et al., 2023). If injury by early herbivores on non-marketable parts leads to reduced herbivory later or to overcompensation by the plant, then herbivory can even be beneficial for crop yield (Garcia & Eubanks, 2019; Poveda et al., 2010). Which herbivores have the most prominent influence on crop yield and quality is determined by the combination of direct and indirect processes, and these processes are likely affected by the agroecological context that the crops are grown in.

In this multi-year and location study, we examined the direct consumptive effect and the indirect plant- or community-mediated effects of herbivores on plant growth, fresh harvestable weight and damage; and if these effects are altered in different cropping systems. For this purpose, we examined white cabbage (*Brassica oleracea* var. *capitata*) plant growth, herbivore community, and fresh weight and damage of individual cabbage heads in five cropping system treatments: a monoculture and four strip cropping treatments (Fig. 1). The strip treatments differed in the number of crop species and cultivars that were included and whether plant- or animal-based fertilizers were used. Using generalized linear models, we tested whether cropping system affects fresh weight and damage of cabbage heads, and herbivore abundance and richness before and during cabbage head formation (i.e. early- and late season). We defined crop damage as the yield lost by manual removal of injured leaves after harvest to obtain a marketable cabbage head (Juventia et al., 2021). Next, we used piecewise structural equation modelling to test three distinct path diagrams with increasing resolution of herbivore variables. First, we explored how well early and late plant size predict fresh harvestable weight, and if cropping system alters these predictions (Fig. S1a). Thereafter, we examined how cropping systems alter the effect of herbivore abundance and richness on fresh harvestable weight and damage, and whether these effects change depending on whether the herbivores are present before or during cabbage head formation (Fig. S1b). Since different herbivores might interact in time and space (Mertens et al., 2021), and because herbivores differ markedly in the type and severity of injury caused by their feeding (Boege & Marquis, 2005), we also examined how four different herbivore groups affected fresh harvestable weight and damage, both before and during crop formation (Fig. S1c). We also hypothesized that herbivores reduce yield by injuring plants, but that the strength of these effects differs among herbivore groups. Furthermore, we expected that the effect of herbivores on fresh harvestable weight and damage would be reduced in more diverse cropping systems. We discuss how plant ontogeny, herbivore community and cropping system interact to alter crop yield and damage. We emphasize how our perception of herbivorous insects as pests is flawed and limits conservation efforts and sustainable farming practices.

**Fig. 1.**
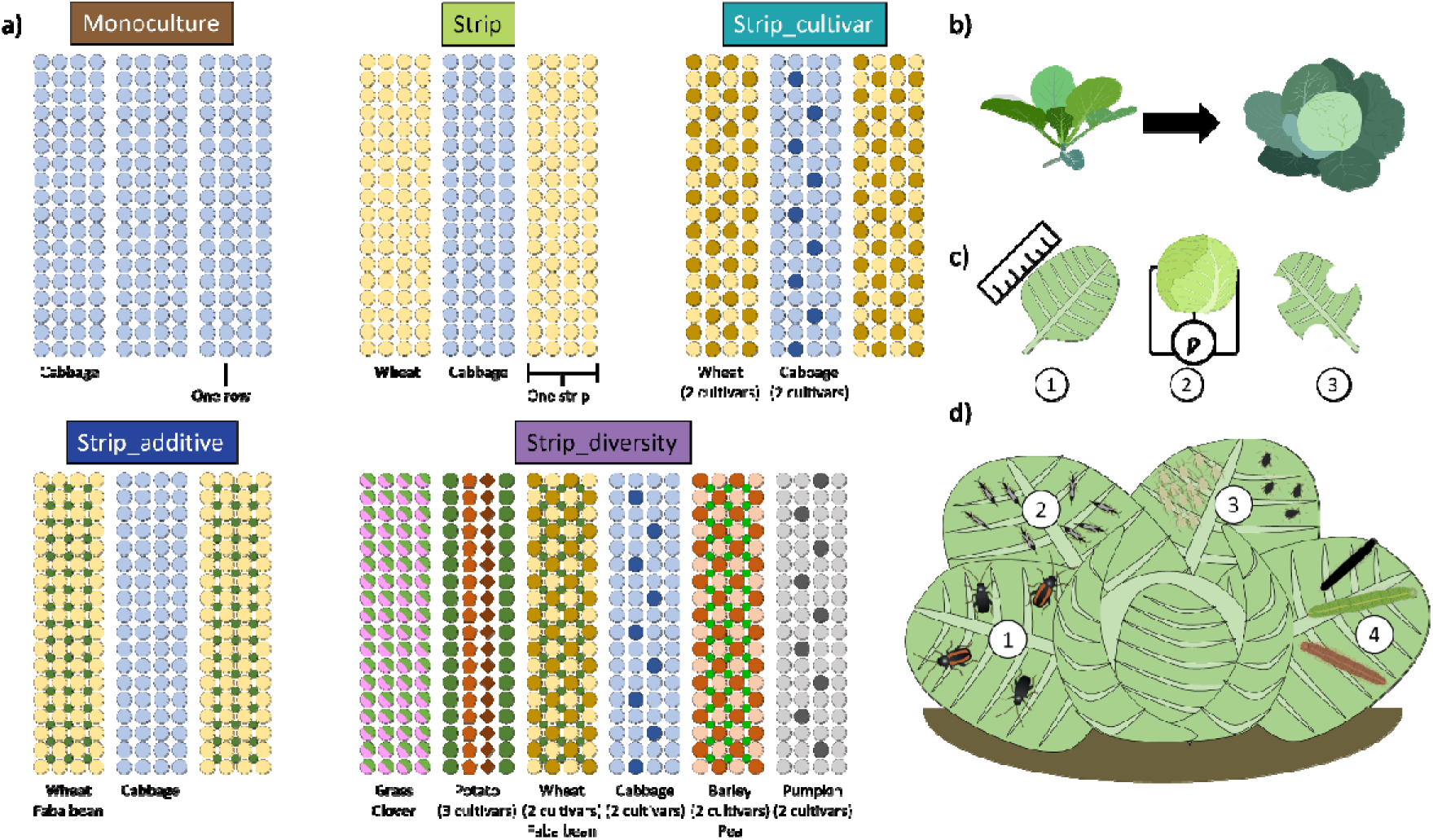
Conceptual representation of experimental set-up and measurements. **(a)** The five cropping system treatments included in this study. Text below strips indicate what was grown in the strips. In the crop configurations with multiple cultivars, these replaced some of the main cultivar plants keeping plant density equal. For the systems with nitrogen-fixing plants (faba bean, pea or clover), these were added to the poaceous crops, increasing plant density. **(b)** We measured individual plants throughout the growing season. **(c)** We measured plant diameter and number of leaves throughout the season (1), total wrapper leaf and cabbage head fresh weights after harvest (2) and damage by chewing insects on the cabbage head (3). **(d)** We counted and identified herbivorous insects and grouped these into four classes: (1) flea beetles, (2) thrips, (3) phloem suckers, and (4) chewing insects. See Table S2 for the species that were classified in each herbivore class.

## Methods

### Study area

The relationship between white cabbage cultivation and insect herbivores was studied over three years in two long-term strip-cropping experiments at two different locations in the Netherlands: Wageningen and Lelystad. Both experimental locations were managed organically without the use of biological or chemical pesticides. The fields in Wageningen were located north of the Wageningen University campus (51°59’24”N, 5°39’36”E, 13 masl), as part of a larger network of experimental fields (Juventia et al., 2021). The soil type was loamy sand. The surrounding landscape was quite diverse with many relatively small experimental fields (<5 ha) and semi-natural habitats like tree lines and ditches. The experimental fields were enclosed by other smaller experimental plots of Wageningen University to the south, a small-scale farm to the west, larger experimental fields to the north, and a busy regional road to the east. The fields at Lelystad were located east of the city of Lelystad at the Wageningen Research Field Crops location (52°32’33”N 5°34’23”E, -4 masl) (Cuperus et al., 2023). The soil type here was sandy clay loam, claimed from the sea in 1957. The surrounding landscape consisted mainly of larger scale experimental fields (5 – 10 ha) and large-scale agriculture (10 – 40 ha), with some small semi-natural habitats throughout the landscape, like small waterways and flower strips.

### Experimental design

At Wageningen, three main crop pairs were used throughout the three years: white cabbage – wheat, potato – grass and pumpkin – barley (see scientific names in Table S1). In 2019, sugar beet was attempted instead of pumpkin, but this crop failed to establish, and these strips were left bare that year. Also, in 2021 a switch was made from wheat to oat. At Lelystad, four main crop pairs were grown: white cabbage – wheat, potato – grass, sugar beet – barley and carrot – onion. The crops were chosen in close collaboration with Dutch farmers to ensure crops were relevant in developing strategies for creating more sustainable crop production systems. The focus of this paper is on white cabbage and associated herbivorous insects.

Five different cropping system treatments were established that differed in spatial and genetic diversity and management, but that followed the same crop sequence over time (Fig. 1). One large-scale monoculture of one crop without genetic variation was established for white cabbage on a reference field at both locations. As the spatial dimensions of the monoculture (at least 45 * 45 meters) were markedly greater than those of the other treatments, we used replication by sub-plots to account for a similar spatial variation. We established four strip treatments: Strip, Strip_cultivar, Strip_additive and Strip_diversity. The simplest strip treatment included white cabbage and wheat or oat in strips of three meters wide (Strip). Here, each strip only included one crop without genetic variation. A single strip of white cabbage consisted of four rows of plants, fitting the optimal planting distance. In Strip_cultivar we introduced intraspecific variation by mixing two cultivars within the strips. In Strip_additive we included legumes within the neighbouring cereal or grass crops and mixtures of endangered arable flora at the edges (see composition in Table S1). Lastly, in Strip_diversity, which was only established at Wageningen, we combined the elements of Strip_cultivar and Strip_additive and included all six main crops in strips next to each other. At Wageningen, the Strip_additive and Strip_diversity systems received organic plant fertilizers instead of animal manure. Crops were rotated and green manures were regularly grown in between two crops in consecutive years. For a more detailed description of seed mixtures, crop cultivars and cropping schemes see Tables S3 – S6.

The treatments were organized in an incomplete block design, as including a monoculture on each field that included all four strip configurations was not possible due to spatial constraints. Thus, at both locations a reference field was used every year that included a monoculture and the simplest strip treatment (Strip). All strip treatments, including Strip, were placed in a complete block design on three (Wageningen) or two (Lelystad) fields. Within a field all treatments were repeated thrice, and only the central four strips of each crop pair per treatment were used for measurements, as the outside two strips could have interfered with neighboring treatments. For a visualization of the field set-up, see Fig. S14.

### Study system

In this agro-ecological study, we examined the interaction between white cabbage yield and quality and the above-ground community of herbivores that naturally occur on white cabbage and were observable by eye. We did not release any of the observed herbivores.

We used white cabbage cultivar Rivera as the main cash crop cultivar, except at Lelystad in 2021, when it was replaced with a similar cultivar, Expect. In the Strip_cultivar and Strip_diversity treatments, we included a 1/8^th^ replacement of Rivera (or Expect) by cultivar Christmas Drumhead. This cultivar is known to be more attractive to several herbivore species and parasitoids (Croijmans et al., 2022, 2024; Poelman, et al., 2009a, 2009b) and was used as a trap crop to lure herbivores and have them culled by natural enemies. Replacement of cabbage cultivars always happened in the central two of four plant rows within the strips, where every fourth Rivera plant was replaced by a Christmas Drumhead plant. The purpose was to maximize the number of Rivera plants that neighbored at least one Christmas Drumhead plant. White cabbage plants were planted at five weeks old, when the wheat or oat already formed a dense cover. Wheat or oat were harvested halfway in the cabbage growing season and replaced by green manure (see composition in Tables S5 - S6).

The invertebrate community of white cabbage is well described and has a set of specialist herbivorous insects that only occur on brassicas. The community of herbivores consisted mainly, but not exclusively, of two specialist flea beetles, *Phyllotreta atra* and *P. undulata* (Coleoptera: Chrysomelidae); two specialist caterpillars, *Plutella xylostella* (Lepidoptera: Plutellidae) and *Pieris rapae* (Lepidoptera: Pieridae); two generalist caterpillars, *Mamestra brassicae* and *Autographa gamma* (Lepidoptera: Noctuidae); a specialist aphid, *Brevicoryne brassicae* (Hemiptera: Aphididae); a generalist aphid, *Myzus persicae* (Hemiptera: Aphididae); a specialist whitefly, *Aleyrodes proletella* (Hemiptera: Aleyrodidae); and thrips, likely *Thrips tabaci* (Thysanoptera: Thripidae). For a full list of herbivores see Table S2. Slugs and snails were also observed, but as these were mostly active at night and early in the morning, their abundances were closely linked to the time of observation. Therefore, we excluded slugs and snails from the analyses.

### Cabbage-invertebrate community monitoring

Monitoring of the invertebrate community involved repeated sampling of the same plants throughout the growing season. Per experimental strip eight cabbage plants were selected equidistantly throughout the strip, excluding the first and last ten meters of each strip. Two cabbage plants were chosen per plant row on each half of the strip length, and within each half the order of the rows was randomized. Within Strip_cultivar, we also sampled eight Christmas Drumhead plants per strip in a similar way as the other cabbage plants. As there was only one replicate of Strip_diversity per field and to increase the sample size of Strip at the reference field, we chose to double the sampling effort in these strips. In the monoculture we sampled cabbage plants in a similar fashion as in the strip treatments, leaving four plant rows free between rows in which we took samples, creating a similar distance as between samples of different strips within the strip treatments. See Table S7 for a dissemination of the number of replications.

Invertebrates on the selected cabbage plants were counted in 3 to 4 rounds per location per year, which were roughly four weeks apart. The first two of these rounds were considered early season, from June 8^th^ to August 5^th^; the second two rounds were considered late season, from August 10^th^ to September 28^th^. After these sampling rounds, the selected cabbage plants were harvested for yield and damage assessment (see below). During monitoring, all invertebrates on the cabbage were identified to the lowest taxonomical unit possible without having to remove the invertebrates, and counted. In addition to monitoring herbivore communities, we measured two variables related to plant size. Diameter of the plant was measured using measuring tape as the greatest distance between two leaf tips on a plant, and the number of leaves was counted.

### Cabbage yield and damage assessment

All sampled cabbage plants were harvested in one operation per location per year and stored in a cooler until fresh harvestable weight and damage assessment. Sampled cabbage harvest happened shortly before the first strip level harvest (organic cabbage harvest usually happens manually in two or three rounds). Cabbage heads were separated from the wrapper leaves and stalk, and inspected for any chewing herbivore injury. Injured head leaves were peeled off until no injury was visible anymore. The wrapper leaves, damaged head leaves and the clean head were weighted separately. Fresh harvestable weight was measured as the total fresh weight of the cabbage head, including injured head leaves. We did not assess marketable yield (the fresh weight after removal of injured leaves) as we believe that the total fresh weight of the cabbage head gives a better indication of how consumptive behavior of herbivores throughout the season affected eventual crop biomass. Such marketable yield would be mostly influenced by the removal of injured leaves caused by late-season herbivory. We considered damage as the proportion of the total cabbage head fresh weight that we removed due to injury and denote this variable as proportion damaged head weight. Taking the ratio avoids an inherent correlation between total cabbage head weight and the weight of the injured leaves, as heavier cabbage heads also have heavier leaves.

### Statistics

All statistical analyses were done in R, version 4.2.2.

To assess the effects of cropping systems on cabbage head fresh weight and proportion damaged head weight, we used generalized linear models using the glmmTMB package (Brooks et al., 2017). For both response variables we used cropping system treatment, placement within the strip (edge or center) and year as fixed effects and field and strip number as random effects. We also included all interactions among fixed variables and performed backward selection using Akaike Information Criterion (AIC) to see which of these interactions increased fit of the model. If taking out an interaction variable reduced the AIC, it was removed from the final model. Because the Strip_diversity treatment was only present in Wageningen, we chose to compare treatments for both locations separately. As there were significant field effects, we separated the two sections of the incomplete block design into distinct analyses: (1) monoculture versus Strip on the reference field, (2) all strip treatments on the other fields. For cabbage head fresh weight, we used a Gaussian distribution. For proportion damaged head weight, the distribution did not fit well with a Gaussian distribution. Therefore, we transformed the proportion to percentage by multiplying the value by 100 and then rounded the value up to the nearest integer. This data transformation allowed us to fit a model with a negative binomial distribution. Because a large portion of heads was undamaged, we encountered zero-inflation in the data, which we corrected for in the final model. We tested model assumptions using the “testResiduals” function of the dHARMA package (Hartig, 2020).

To assess the effect of early plant variables and herbivores on the fresh weight of cabbage heads, and to examine how these effects differ among the treatments, we used piecewise structural equation modelling (pSEM), using the “piecewiseSEM” package (Lefcheck, 2016). We made three models of increasing complexity: (1) model without herbivores (Fig. S1a), (2) model with herbivore abundance and richness (Fig. S1b), and (3) model with four herbivore groups (Fig. S1c) (for more details on the SEM models see text S1). In all pSEM models, we used year and location as fixed factors in all component models. For all three models we first conceived a model that fit with the hypotheses. Thereafter we checked the fit of this initial model on the full dataset using Fisher’s C. Using a mix of back- and forward selection, we added paths that were vital for the model to fit well. We similarly checked the fit on subsets for each year, each location, and each crop configuration. We removed non-significant paths if removal did not reduce fit of the model fitted on all data and fitted on any of the subsets. The eventual accepted model was one that fit with the whole datasets and all individual subsets. We tested each component model within the pSEM for model assumptions using the “testResiduals” function of the dHARMA package (Hartig, 2020).

## Results

We intensively monitored a total of 1,308 cabbage plants throughout the growing season over the course of three years at two experimental field locations (Wageningen and Lelystad). Here, we aggregated insect-community data as early- or late-season, i.e. before and during cabbage head formation, respectively. After harvest, we assessed fresh harvestable weight by measuring fresh weight of individual cabbage heads (including injured leaves), and damage by calculating the percentage of fresh weight lost due to the manual removal of leaves injured by chewing herbivores. In total, we identified and counted 116,478 herbivorous insects, which consisted of 43% phloem feeders (mainly the aphid *Brevicoryne brassicae* and whiteflies (Aleyrodidae)), 26% leaf chewing insects (mainly the lepidopterans *Plutella xylostella* and *Pieris rapae*), 20% thrips (Thysanoptera) and 11% flea beetles (*Phyllotreta atra* and *P. undulata*) (Table S2).

### Cabbage yield and damage

Cabbage plants grown in strip cropping with six crops had higher yield than those grown in the other, less diverse treatments (Fig. 2a). The effect of treatments with two crops or crop mixtures on yield depended on the year (Fig. S2, Table S8). At Wageningen, no difference was observed between strip cropping with one (Strip) or two (Strip_cultivar) cabbage cultivars (Fig. S2). However, at Lelystad, Strip_cultivar outperformed Strip in 2019 and 2021, with the increase in yield of the main cultivar (Rivera) even compensating for its 1/8^th^ replacement by the second, unmarketable cultivar (Christmas Drumhead) (Fig. S2). The treatment that included legumes (Strip_additive), did not differ from the treatment without legumes (Strip) (Fig. S2).

**Fig. 2.**
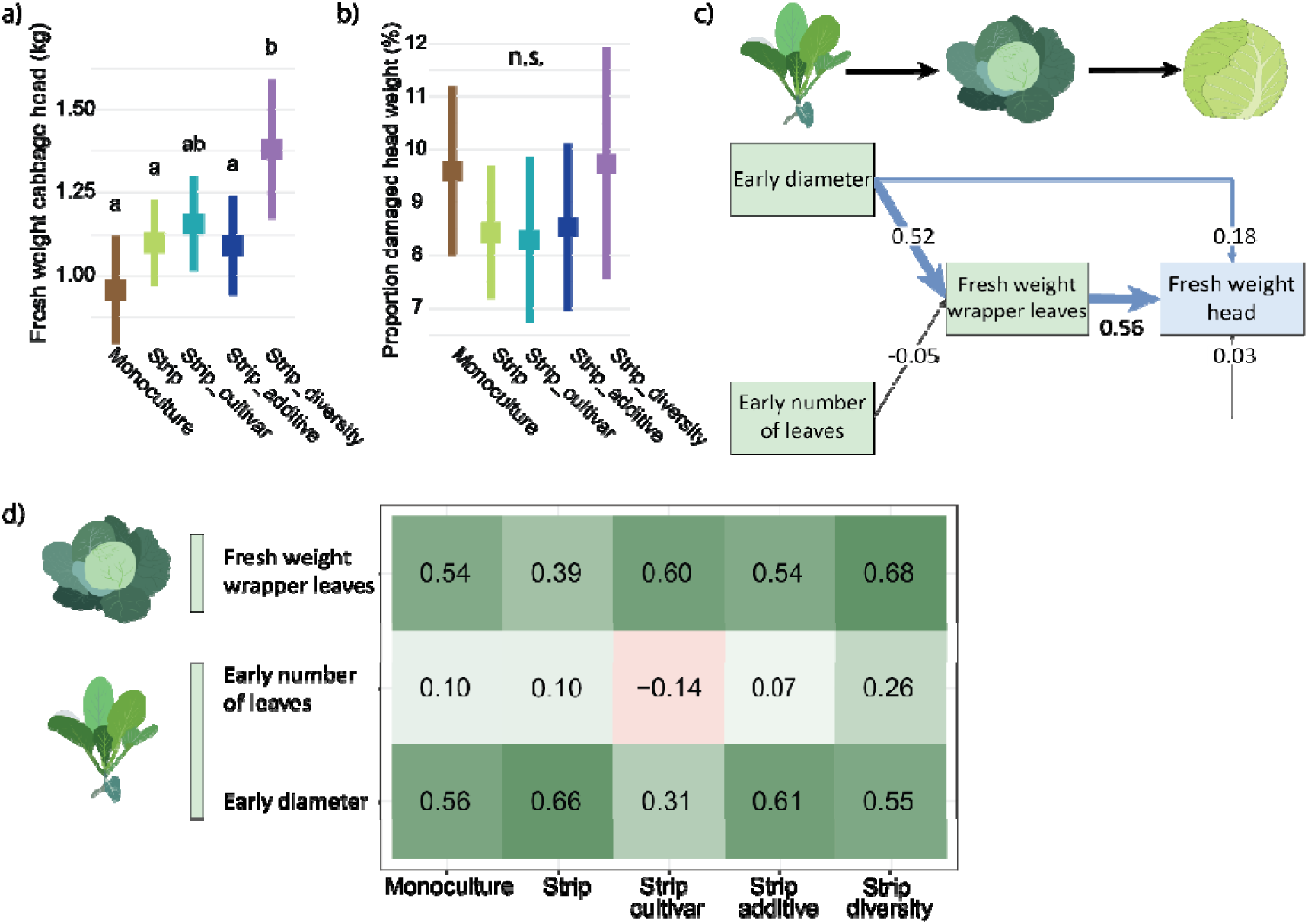
The effect of cropping system and plant size variables on fresh weight of the cabbage head. The overall effect of cropping system on fresh weight of the whole cabbage head **(a)** and the proportion of fresh weight lost through removal of leaves injured by chewing herbivores (damage) **(b)**, averaged across locations and years. Colors indicate different crop configurations. Compact letter display was used to indicate significant differences between crop configurations, whereas “n.s.” indicates no significant effect among crop configurations. The squares indicate estimated means; the vertical lines show the 95% confidence interval. **(c)** Structural equation model (SEM) fitted on the full data set. Standardized parameter estimates are given on each arrow and arrow width indicates the size of the standardized parameter estimates. Arrow color indicates the sign of the parameter estimate (blue = positive, black = not significant). Green squares indicate plant variables related to plant size and the blue square indicates the plant variable related to crop yield. **(d)** Total standardized effect of the three predictor variables on fresh weight of the whole cabbage head, per crop configuration. Cabbage plants were grown in five treatments (Fig. 1). The color gradient indicates the size and sign of the effect. A separation in direct and indirect effects of variables on fresh weight head are given in Table S22.

Overall, cropping system had no effect on the proportion of cabbage head weight removed due to herbivore injury, i.e., on damage (Fig. 2b). However, the effect of cropping system on damage differed between years and locations (Fig. S3, Table S9). In 2019 at Lelystad, damage was lower in strip cropping with one cultivar and no legumes in the rotation (Strip), than in the monoculture or in both more diverse strip treatments (Strip_cultivar and Strip_additive) (Fig. S3). In 2019 at Wageningen, damage was lower in the strip treatment with legumes in the rotation (Strip_additive), compared to the two treatments that included two cabbage cultivars (Strip_cultivar and Strip_diversity) (Fig. S3). However, in 2020 and 2021, there was no effect of cropping system on damage at either location (Fig. S3).

### Herbivore abundance and richness

Both early and late season, insect herbivore abundance tended to be higher in treatments with more than two crops or with two cultivars, than in monocultures (Fig. 3b, S4, Tables S10, S11). Herbivore richness was generally higher in the treatment with six crops compared to any of the other crop treatments (Fig. 3c, S5, Tables S12, S13). Early leaf chewer abundance was higher in all strip treatments than in monocultures at Lelystad in 2019 and 2020, when leaf chewer abundances were high (Fig. S6, Tables S14, S15). However, when leaf chewer abundances were low (Wageningen in all years and Lelystad in 2021), only the most diverse treatment (Strip_diversity) had higher leaf chewer abundances than the other treatments. There was no consistent effect of treatment on early and late phloem sucker abundance, and early thrips (Fig. S7, S8, Table S16 - S18). However, during an outbreak of late-season thrips at Lelystad in 2019, there were significantly more thrips in the strip treatments than in the monoculture (Fig. S8, Table S19). Both early and late flea beetle abundances were generally lower in the monoculture than in the strip treatments at Lelystad, but at Wageningen late flea beetle abundance was higher in the monoculture in two years (Fig. S9, Table S20, S21).

**Fig. 3.**
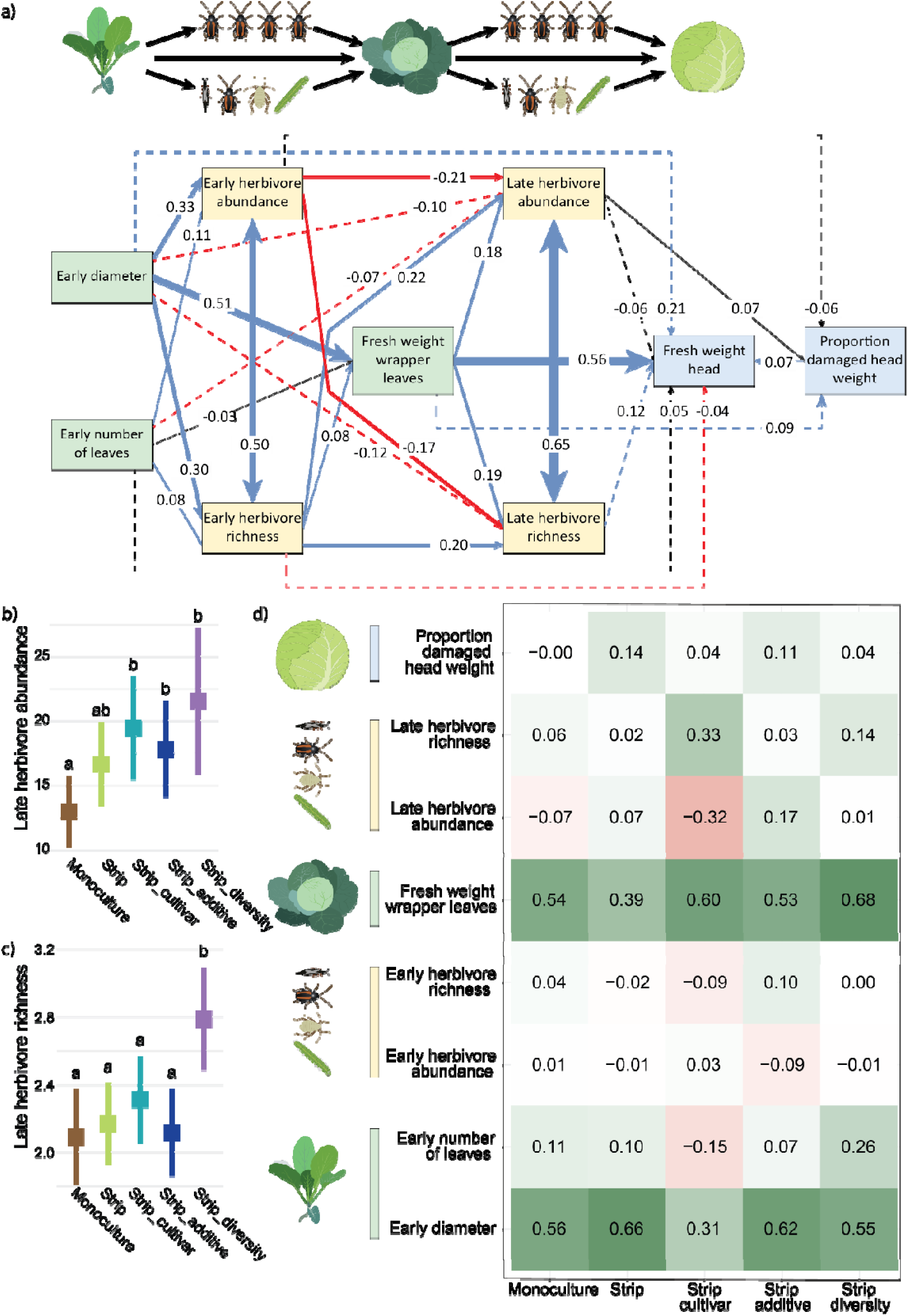
The effect of herbivore abundance and richness on fresh weight and damage of the cabbage head. **(a)** Structural equation model (SEM) fitted on the full data set. Standardized parameter estimates are given on each arrow and arrow width also indicates the size of the standardized parameter estimates. Arrow color indicates the sign of the parameter estimate (blue = positive, red = negative, black = not significant). Dotted arrows indicate paths that were included during the fitting process. Double headed arrows represent relations for which we could not presume a biologically meaningful causal relation, and which were included by their correlated error structure. Green squares indicate plant variables related to plant growth, yellow squares indicate herbivore variables and blue squares indicate plant variables related to crop quantity and quality. **(b,c)** Overall effect of crop configurations on late herbivore abundance **(b)** and richness **(c)**, averaged across locations and years. **(d)** Total standardized effect of the predictor variables on fresh weight of the whole cabbage head per crop configuration. Cabbage plants were grown in five treatments (Fig. 1). The color gradient indicates the size and sign of the effect. A separation in direct and indirect effects of variables on fresh weight head and proportion damaged head weight are given in Tables S23.

The two cabbage cultivars within the strip treatment with two cabbage cultivars (Strip_cultivar) differed in several herbivore variables. At Wageningen, herbivore abundance and richness were generally higher on Christmas Drumhead, whereas the cultivars did not differ significantly at Lelystad (Fig. S4, S5, Tables S10-S13). Leaf chewer and thrips abundance did not differ consistently between the two cultivars for both locations (Fig. S6, S8, Tables S14, S15, S18, S19). However, phloem suckers and flea beetles were regularly more abundant on Christmas Drumhead at Wageningen, whereas flea beetles were usually more abundant on Rivera plants at Lelystad (Fig. S7, S9, Table S16, S17, S20, S21).

### Mixing cultivars, but not crops, reduces predictability of fresh harvestable weight from early-season plant size

First, we examined how well plant size before and during cabbage head formation predicts fresh harvestable weight and how cropping systems alter these predictions. For this purpose, we fitted piecewise SEM on the full dataset and made separate models per treatment to assess differences in the pathways. For all SEM models, we first tested the fit of the hypothesized model. When fit was insufficient, based on the Fisher’s C value, we tested alternative models to reach a model that best fitted the data and thereby explored which pathways best predicted fresh harvestable weight. Early plant size had a strong positive indirect effect on fresh harvestable weight via increasing late plant size, but also a direct positive effect on fresh harvestable weight (Fig. 2c, Table S22). This direct positive effect suggests that large plant size early in the season also positively affects harvestable weight irrespective of its effect on plant growth, for example by reducing herbivore abundances during crop development. The effect of early plant size on fresh harvestable weight was consistent for all treatments, except for the simplest strip treatment with two cabbage cultivars (Strip_cultivar) (Fig. 2d, S10). Here, the positive effect of early plant diameter on fresh harvestable weight was lower than in other treatments, and there was a distinct negative effect of increased early number of leaves. These results show that the addition of a second cultivar within the cropping system reduces how well early plant size predicts crop yield, an effect that seems to be alleviated by including a higher diversity of crops.

### Herbivores differently affect fresh harvestable weight depending on cropping system

Next, we examined how plant size and herbivore abundance and richness at two plant developmental stages affected fresh harvestable weight and damage. We again used piecewise SEM with separate models for the treatments. Early plant size correlated positively with early, but negatively with late herbivore abundance and richness (Fig. 3a). The variation in herbivore abundances and richness, in turn, did not lead to variation in yield and damage in all treatments, except for strip cropping with two cultivars (Fig. 3a,d, S11, Table S23, S24). In the simplest strip treatment with two cultivars (Strip_cultivar), higher late herbivore abundance reduced fresh harvestable weight and increased damage, whereas late herbivore richness had a positive effect on fresh harvestable weight (Fig. 3a,d, S11). Interestingly, in both the simplest strip treatment (Strip) and the treatment that included legumes (Strip_additive), greater damage was associated with greater fresh harvestable weight (Fig. 3d, Table S24). These results show that herbivore abundance does not consistently correlate with negative effects on fresh harvestable weight and damage. Also, we found that cropping system treatments alter the strength and sign of the specific effects of herbivores on crop quantity and quality, especially when the treatment included two cultivars (Fig. 3d, S12). Intriguingly, the direct effect of early plant size on fresh harvestable weight was like that in the model that did not include any effects of herbivores (Fig. 2c, 3a, Tables S22, S23), possibly due to early-plant-size mediated changes in specific herbivore groups, instead of overall herbivore abundance.

### Herbivore groups differently affect crop yield among cropping systems

Lastly, we examined how plant size and the abundance of four herbivore groups before and during cabbage head formation affected fresh harvestable weight and damage, and how these effects varied among crop configurations. We again used piecewise SEM with separate models for the treatments. Early plant size correlated positively with early abundance of most herbivore groups, but the herbivores had mostly low correlations with late plant size (Fig. 4a), yield (Fig. 4b) or damage (Fig. S13). Important exceptions were that in the simplest strip treatment (Strip) and in the simplest treatment with two cultivars (Strip_cultivar) early leaf chewers were negatively correlated with yield (Fig. 4b), and in both treatments that included two cultivars (Strip_cultivar and Strip_diversity) high abundances of early chewers were associated with lower damage (Fig. S13, Table S26). Late herbivore abundances were affected by a combination of early and late plant size, and the abundances of distinct early herbivore groups (Fig. 4a). In turn, the effect of late herbivores on fresh harvestable weight differed among the herbivore groups and the treatments, whereas the variation in late herbivore abundances did not lead to variation in damage whatsoever. In the monoculture, late thrips negatively correlated with fresh harvestable weight, whereas late flea beetles correlated positively (Fig. 4b). In both the least and most diverse strip treatments (Strip, Strip_diversity) late herbivores only had marginal effects on fresh harvestable weight (Fig. 4b). In the simplest treatment with two cabbage cultivars (Strip_cultivar) late thrips were associated with reduced fresh harvestable weight (Fig. 4b). In the treatment with legumes (Strip_additive), late leaf chewers negatively affected fresh harvestable weight, whereas flea beetles were associated with higher fresh harvestable weight (Fig. 4b). The direct effect of early plant size on fresh harvestable weight was again similar to previous models, despite the inclusion of herbivore groups (Fig. 2c, 3a, 4a, Table S25). These results show that (1) different herbivore groups have different effects on crop yield and damage, (2) the effects of specific herbivores differ depending on the time of arrival and the cropping system (Fig. 5), (3) early herbivores only have minor effects on plant growth, and (4) early herbivores can have stronger effects on crop quality than late herbivores, despite not being present whilst the harvestable plant parts are growing.

**Fig. 4.**
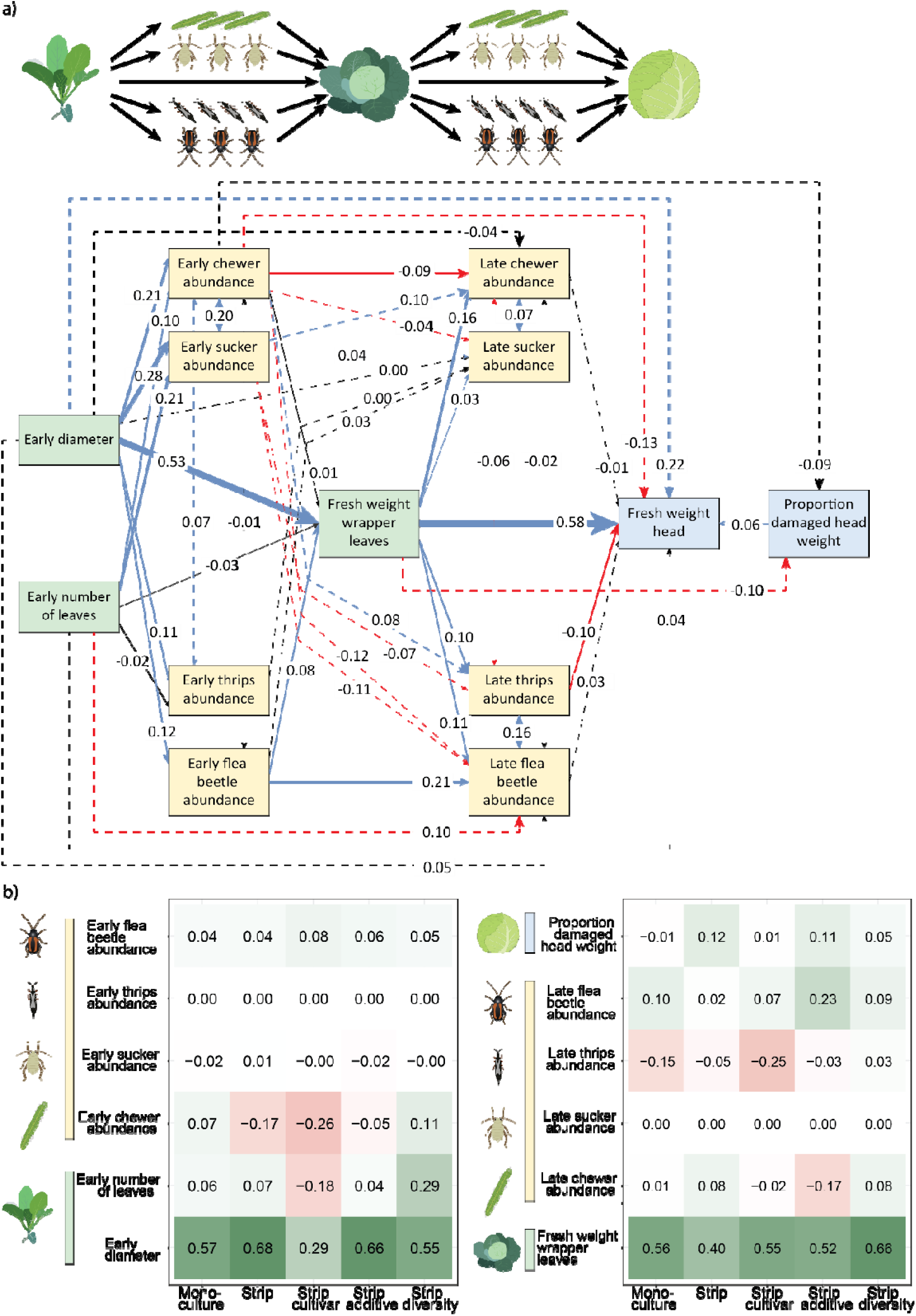
The effect of herbivore groups on fresh weight of the cabbage head. **(a)** Structural equation model (SEM) fitted on the full data set. Standardized parameter estimates are given on each arrow and arrow width also indicates the size of the standardized parameter estimates. Arrow color indicates the sign of the parameter estimate (blue = positive, red = negative, black = not significant). Dotted arrows indicate paths that were included during the fitting process. Double headed arrows represent relations for which we could not presume a biologically meaningful causal relation, and which were included by their correlated error structure. Green squares indicate plant variables related to plant growth, yellow squares indicate herbivore variables and blue squares indicate plant variables related to crop quantity and quality. **(b)** Total standardized effect of the predictor variables on fresh weight of the whole cabbage head per crop configuration. Cabbage plants were grown in five treatments (Fig. 1). The color gradient indicates the size and sign of the effect. A separation in direct and indirect effects of variables on fresh weight head and proportion damaged head weight are given in Tables S25.

**Fig. 5.**
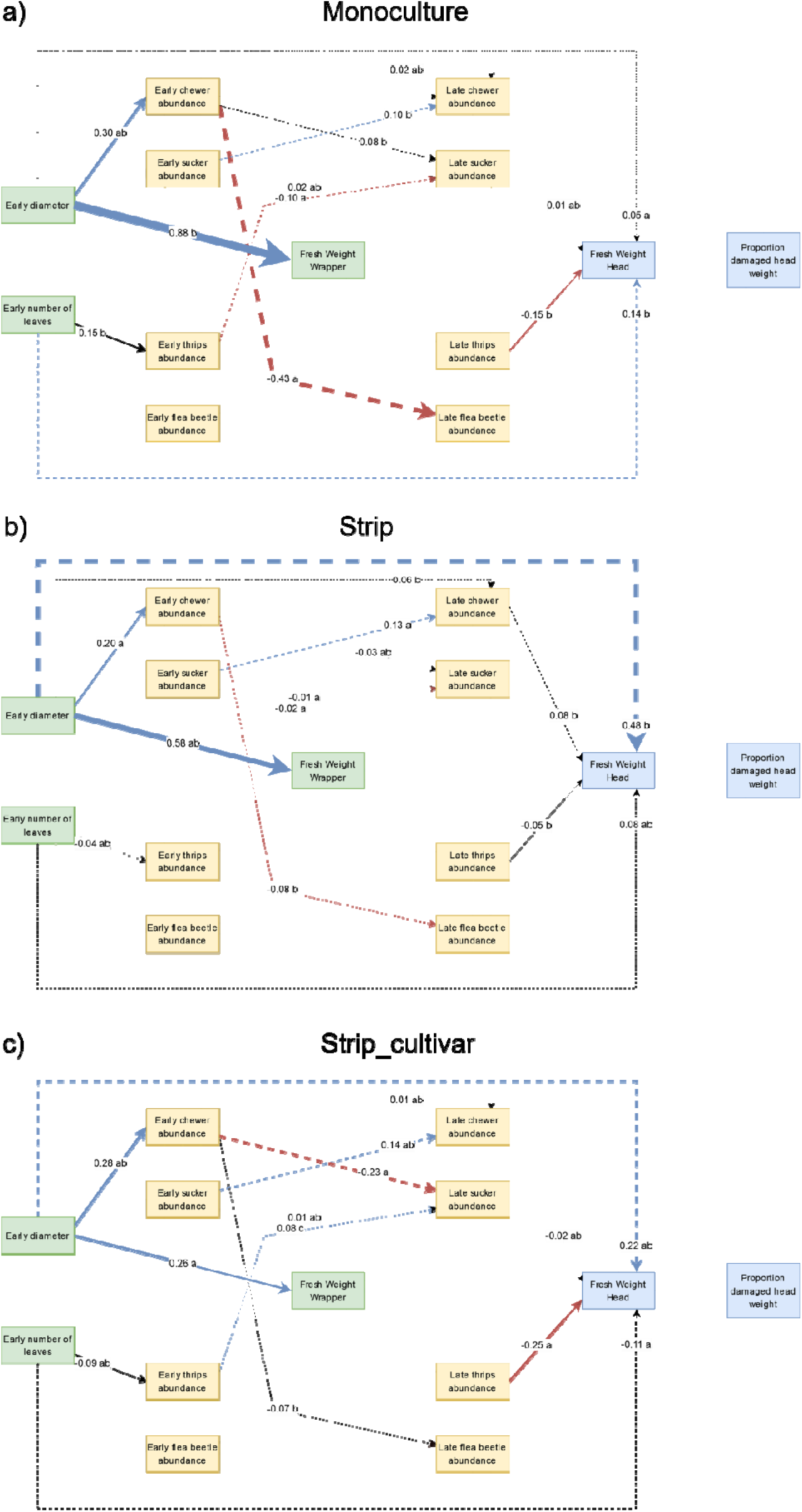
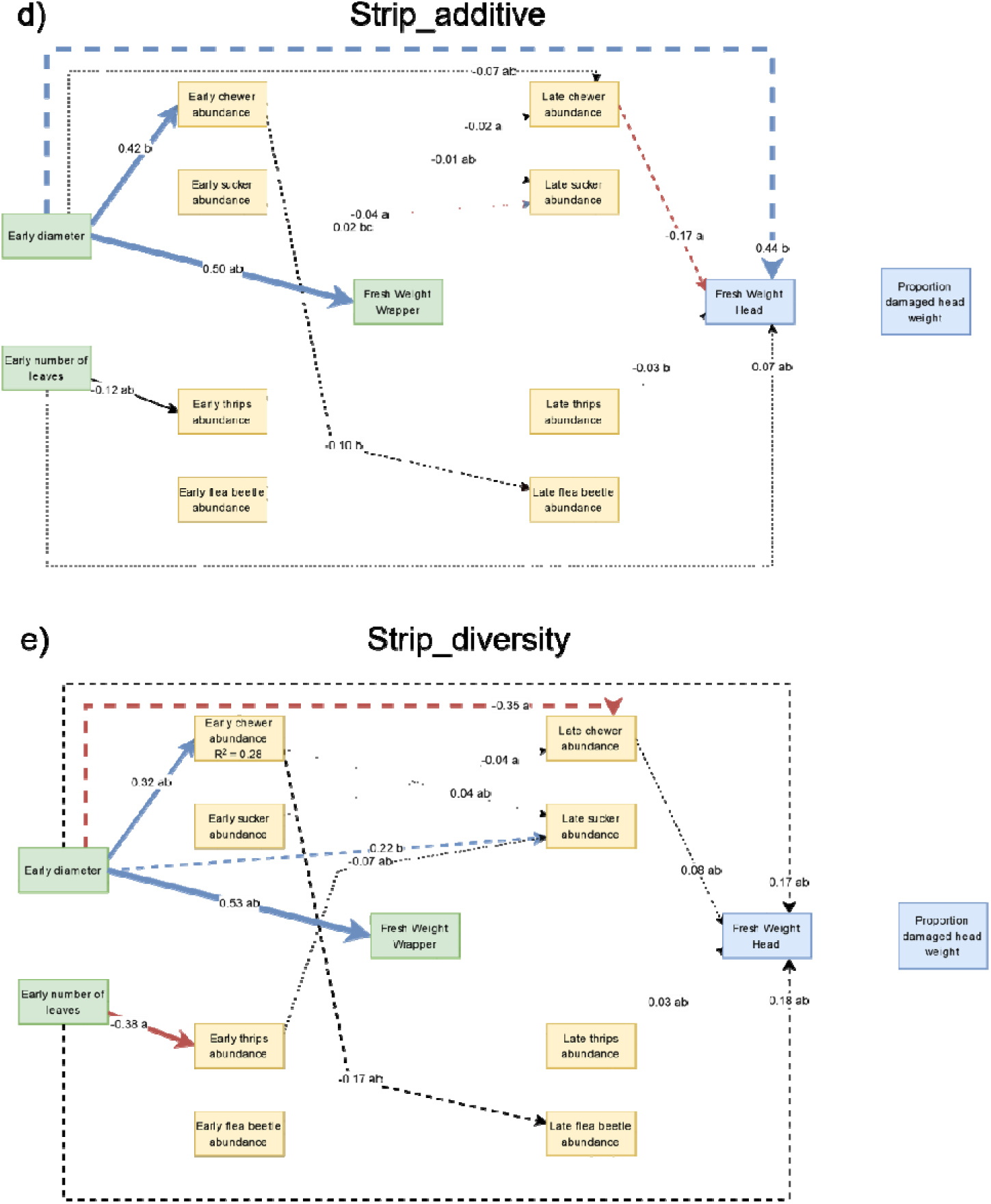
Comparison of pSEM models with herbivore groups per treatment. Cabbage plants were grown in five treatments (Fig. 1). Standardized parameter estimates are given on each arrow and arrow width also indicates the size of the standardized parameter estimates. Arrow color indicates the sign of the parameter estimate (blue = positive, red = negative, black = not significant). Compact letter display was used to indicate significant differences for paths among crop configurations, where the paths among two crop configurations were considered significantly different when their confidence intervals did not overlap. Arrows with reduced opacity were not significantly different among all crop configurations. Dotted arrows indicate paths that were included during the fitting process. Double headed arrows represent relations for which we could not presume a biologically meaningful causal relation, and which were included by their correlated error structure. Green squares indicate plant variables related to plant growth, yellow squares indicate herbivore variables and blue squares indicate plant variables related to crop quantity and quality. A separation in direct and indirect effects of variables on fresh weight head and proportion damaged head weight are given in Tables S25 & S26.

## Discussion

Cropping systems alter how herbivores affect crop yield, as simply looking at the abundance of herbivores as a proxy for pest control foregoes differences in survival of and injury by herbivores. Our results show minor effects of cropping systems on herbivore abundances, but herbivores did show distinct cropping-system-dependent effects on fresh harvestable weight. For example, the most diverse strip treatment (Strip_diversity) had the heaviest cabbage heads, whilst also having the highest abundance of herbivores. These seemingly contrasting results fit with previous findings in the same experimental fields in which, despite higher egg abundances, the survival of a cabbage herbivore was lower (Karssemeijer et al., 2023), abundance and richness of both ground-dwelling and aerial predators were higher (Croijmans et al., unpublished; Cuperus et al., 2023) and parasitism by a natural enemy of a cabbage herbivore was higher (Croijmans et al., 2024) in more diverse cropping systems. As a result, fewer adult herbivores of cabbage (Cuperus et al., 2023), lower foliar injury (Juventia et al., 2021) and more injury-free product were observed in more diverse cropping systems (Carrillo-Reche et al., 2023), although this did not translate into a significant increase in yield (Carrillo- Reche et al., 2023; Ditzler et al., 2023). Thus, the higher abundance of herbivorous insects in more diverse cropping systems might be successfully controlled by naturally occurring predators, which would reduce the impact of these herbivores on crop yield and quality. The increase in fresh harvestable weight in the most diverse strip crop configuration might thus not be caused by a reduction in the abundance of herbivores compared to less-diverse cropping systems, but by other biotic or abiotic factors. Nitrogen availability under different fertilization regimes (Han et al., 2022), competition or niche complementarity among crop species (Bourke et al., 2021), changes in arable flora (Kremen & Miles, 2012), soil microbial communities under different crop rotations (Viketoft et al., 2021), belowground herbivores (Karssemeijer et al., 2023) and disease prevalence (Ditzler et al., 2021) are all factors that may alter plant growth and which were not measured in this study. Our results indicate that highly diverse strip-cropping systems facilitate beneficial interactions that allow herbivores to coexist on crops without causing economic damage, whilst simultaneously increasing crop yield.

Early plant size was a good predictor of fresh harvestable weight in most cropping systems, but inclusion of two cultivars within the system reduced the strength of this prediction. Cabbage is known as a bad competitor early in the season and cropping systems where companion plants are planted or sown before cabbages reduce cabbage yields (Carrillo-Reche et al., 2023). Our results reinforce that early plant growth is vital for crop yield. Cropping systems designed to enhance cabbage productivity should aim to avoid competition or damage to freshly planted cabbage plants, whereas negative trade-offs for cabbage can be accepted later in the season. In cultivar mixtures, plant growth might be more unpredictable as it also depends on the cultivar composition of the neighboring plants, which changes competitive interactions compared to a cropping system with only one cultivar (Bourke et al., 2021). Moreover, although we mostly did not observe higher herbivore abundances in this cropping system, we did observe the strongest negative effects of herbivores on fresh harvestable weight in the cropping system with two cultivars. Both leaf chewers and thrips negatively affected fresh harvestable weight, but chewers did so only early and thrips only late in the season. For thrips, this difference between early and late abundance might be due to thrips only reaching damage thresholds late in the season, during an observed outbreak. These observations might indicate that our choice for a cultivar that is attractive to both herbivores and their natural enemies (Croijmans et al., 2022, 2024; Poelman, et al., 2009a, 2009b) might have backfired. We expected this cultivar to serve as a trap crop that would facilitate beneficial tri-trophic interactions by bringing herbivore and natural enemy together.

However, this might also reduce the chance that natural enemies would visit the cash cultivar when herbivores still end up there, and this is reinforced by non-host herbivores that can also increase parasitoid recruitment by the attractive cultivar (Croijmans et al., 2022). In the most diverse strip cropping system (Strip_diversity), which also included both cultivars, this reduction in predation might be mediated by increased predation facilitated by the higher crop diversity (Croijmans et al., 2024). Thus, the attractive cultivar might interfere with biological control by naturally occurring natural enemies on the cash cultivar, which would risk a negative effect on yield. This higher risk to herbivore damage comes on top of a reduction in yield due to the replacement of 1/8^th^ of the cabbages by the attractive, unmarketable cultivar. Any crop management choice will have its own suite of alterations to herbivore-mediated effects on crop yield, which we need to understand better if we are to design cropping systems that enhance pest control.

Early chewers reduced crop damage in the simplest strip configuration (Strip) and strip cropping with two cabbage cultivars (Strip_cultivar), partly through decreased abundances of late chewers and flea beetles, but also through other, unexplained causes. Potentially, the presence of early herbivores not only reduced the abundance of late herbivores, but also their performance or survival (Ohgushi, 2005). Early herbivores affect the herbivore community via plant-mediated interactions, which has cascading effects on crop yield and quality (Poelman et al., 2023). Herbivory can change secondary metabolites, which alter attraction or apparency of the host plant to later herbivores and their natural enemies (Croijmans et al., 2022; Van Zandt & Agrawal, 2004). Furthermore, herbivores can affect the performance of later herbivores through complementary or antagonistic defensive pathways (Mertens et al., 2021). Such plant-mediated effects of early herbivores on later herbivore presence and performance can have cascading effects on yield and especially on crop quality (Stam et al., 2014). These indirect, community-driven effects on crop yield are complex by nature, but necessary to understand when designing ecology-based alternatives to intensive agriculture.

Our study on white cabbage shows that key players from farm to fork might be too worried about insect herbivores and the crop injury they cause for three key reasons. Firstly, herbivores had mostly minor effects on crop yield, despite not using any pesticides. Non-target effects of pesticides can reduce the abundance of natural enemies (Hallmann et al., 2014). Even targeted pesticide applications, including those in integrated pest management strategies, interfere with prey-predator cycles (Janssen & van Rijn, 2021; Keasar et al., 2023) and in doing so ignore the largely unknown effects of environmental context on herbivore-crop interactions, as we showed in this study. New approaches that include natural enemies (Keasar et al., 2023) reveal greater acknowledgement for the potential of cropping systems to self-regulate damage levels below economic thresholds. Secondly, for leafy vegetables like white cabbage, marketable yield can be increased by breaking false notions of product quality (Dara, 2019). In our study, the biggest losses in crop biomass were generated by the removal of leaves slightly injured by herbivores, not by the injury caused by the herbivores themselves. Thirdly, herbivores play an integral role in food webs and their structural removal in agricultural systems is a bottom-up cause for the loss of (insect) biodiversity (Beaumelle et al., 2023; Vidal & Murphy, 2018). Our work shows that, using agroecological management, there is potential for sharing our crops with insect herbivores without incurring major losses in yield.

Agriculture finds itself at the crossroads of continuing preventative measures against herbivorous pests or adopting more ecological approaches to maintaining crop yield. Many preventative measures are no longer societally accepted, but farmers, their collectives and governments are still hesitant towards phasing out pesticides (Bakker et al., 2021). Diversification of crops by strip cropping offers strong potential to control insect herbivores via ecological processes. Such ecological solutions to agricultural problems need an ecological mindset, that considers the complexities of plant responses and herbivore communities. Future academic research should focus on unraveling the mechanisms underlying these complexities. For the short term, our work shows that crop diversification with six or more crops reduces the impact of herbivory and increases crop yield, all while being manageable under current, organic farming practices.

## Acknowledgements

First, we are grateful to Hilde Faber, Kyra Vervoorn and Rick Wijnberg for their contribution to data collection. This work could not have been completed without the help of many colleagues and students, for this we thank Maria-Franca Dekkers, Stella Juventia, Wessel Rijkens, József Takács, Alessia Vitiello, Max Wantulla and Els van de Zande. We greatly appreciate the seeds of Rivera cv. that were sponsored by Bejo Zaden B.V., the Netherlands. Thanks go to Olivia Elsenpeter, Esther Hofkamp, Titouan le Noc and the Unifarm staff, and in particular Andries Siepel and Peter van der Zee, for maintaining the Wageningen fields; and to the team at Wageningen Open Fields, and in particular Joost Rijk and Laurens van Run, for maintaining the Lelystad fields. Lastly, we greatly appreciate the valuable feedback from Marcel Dicke, Zoë Delamore, and Walter Rossing on earlier versions of this manuscript.

## Statements

### Author contributions

*Luuk Croijmans:* Conceptualization, Methodology, Formal analysis, Investigation, Data curation, Writing – original draft, Writing – review & editing, Visualization, Supervision. *Daan Mertens:* Conceptualization, Methodology, Formal analysis, Writing – review & editing. *Dirk F. van Apeldoorn:* Conceptualization, Writing – review & editing, Visualization, Supervision, Project administration, Funding acquisition. *Yufei Jia:* Investigation, Data curation. *Nelson Ríos Hernández:* Investigation. *Erik H. Poelman:* Conceptualization, Methodology, Writing – original draft, Writing – review & editing, Supervision, Project administration, Funding acquisition.

### Ethics approval

This research was not reviewed by an institutional or governmental regulatory body as the work was performed on invertebrates.

### Availability of data and materials

Data will be made publicly available upon acceptance.

## Supplementary text

### Text S1. Additional information on SEM models

For the first model we included three predictors for fresh weight of the cabbage head (Fig. S1a). We expected that early plant size, measured as plant diameter and number of leaves, would positively affect late plant size, measured as fresh weight of the wrapper leaves. This in turn would positively affect fresh weight of the cabbage head. We also expected that early plant size would have its own effect on fresh weight of the cabbage head, separately from simply having the head start in size. Both fresh weight of the wrapper leaves and of the head were log transformed, and Gaussian distribution was used for all models within the pSEM structure.

For the second model, we supplemented the first model with four variables for early and late herbivore abundance and richness, and we added proportion damaged head weight (Fig. S1b). The effect of herbivory on crop yield is dependent on the developmental stage of the plant and plant growth is affected by herbivory (Tiffin, 2002). Simultaneously, larger plant size leads both to larger biomass of harvestable plant parts and aggregation of herbivores, either by chance or as an active choice in food-plant selection by herbivores (Castells et al., 2016; Van Zandt & Agrawal, 2004). Thus, at the plant level, crop biomass regularly correlates positively to herbivore abundances, via plant size throughout the growing season. Therefore, we expected that early plant size would positively affect early herbivore abundance and richness. In turn, early herbivore abundance and richness would reduce late plant size. Also, we expected late herbivore abundance and richness to positively affect proportion damaged head weight, which in turn would be negatively related with fresh weight of the cabbage head. Again, fresh weight of the wrapper and the head, and both early and late herbivore abundances were log transformed, and Gaussian distribution was used for all models within the pSEM structure.

Considering that herbivore species distinctly interact throughout the growing season (Mertens et al., 2021), and due to the variation in the kind and intensity of damage caused by individuals of different herbivore species (Boege & Marquis, 2005), we included the abundance of four herbivore groups in the final model: leaf chewers (mostly lepidopteran larvae), phloem suckers (aphids and whiteflies), thrips and flea beetles (Fig. S1c). We expected again that early plant size would positively affect early abundance of all four herbivore groups, and these herbivores would subsequently negatively affect late plant size. Late herbivores would then again be positively affected by late plant size, but negatively affected cabbage head fresh weight. As the damage that we measured was chewing damage, we expected both late chewers and flea beetles to have a direct positive effect on proportion damaged head weight and thus indirectly a negative effect on fresh weight of the cabbage head. We expected late phloem suckers and thrips to have a direct negative effect on fresh weight of the cabbage head. We could not reliably predict how herbivore groups would respond to conspecifics within and between time periods. Therefore, we assumed that the abundance of early herbivore groups would have a negative effect on the same groups later in the season, due to induced plant defenses. We then let pSEM exploratively fill in the other interactions among herbivore groups. Again, fresh weight of the wrapper and the head, and early phloem sucker abundance were log transformed, and for these models Gaussian distribution was used. For all other herbivore abundances and for proportion damaged head weight we used negative binomial distribution.

