## Supplemental figures for "Herbivore prevalence poorly predicts yield in diverse cropping systems"

**Supplementary figures**

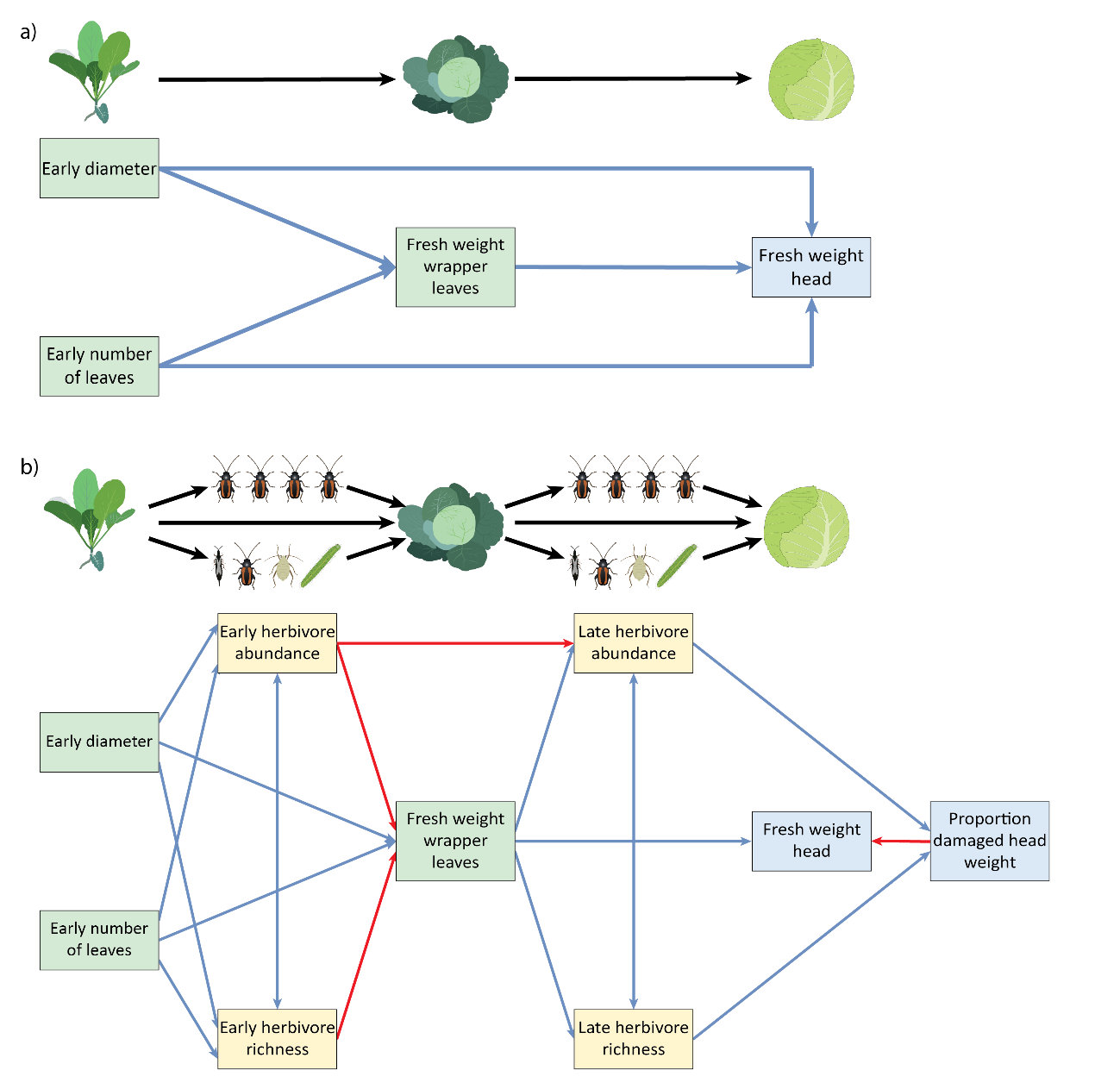

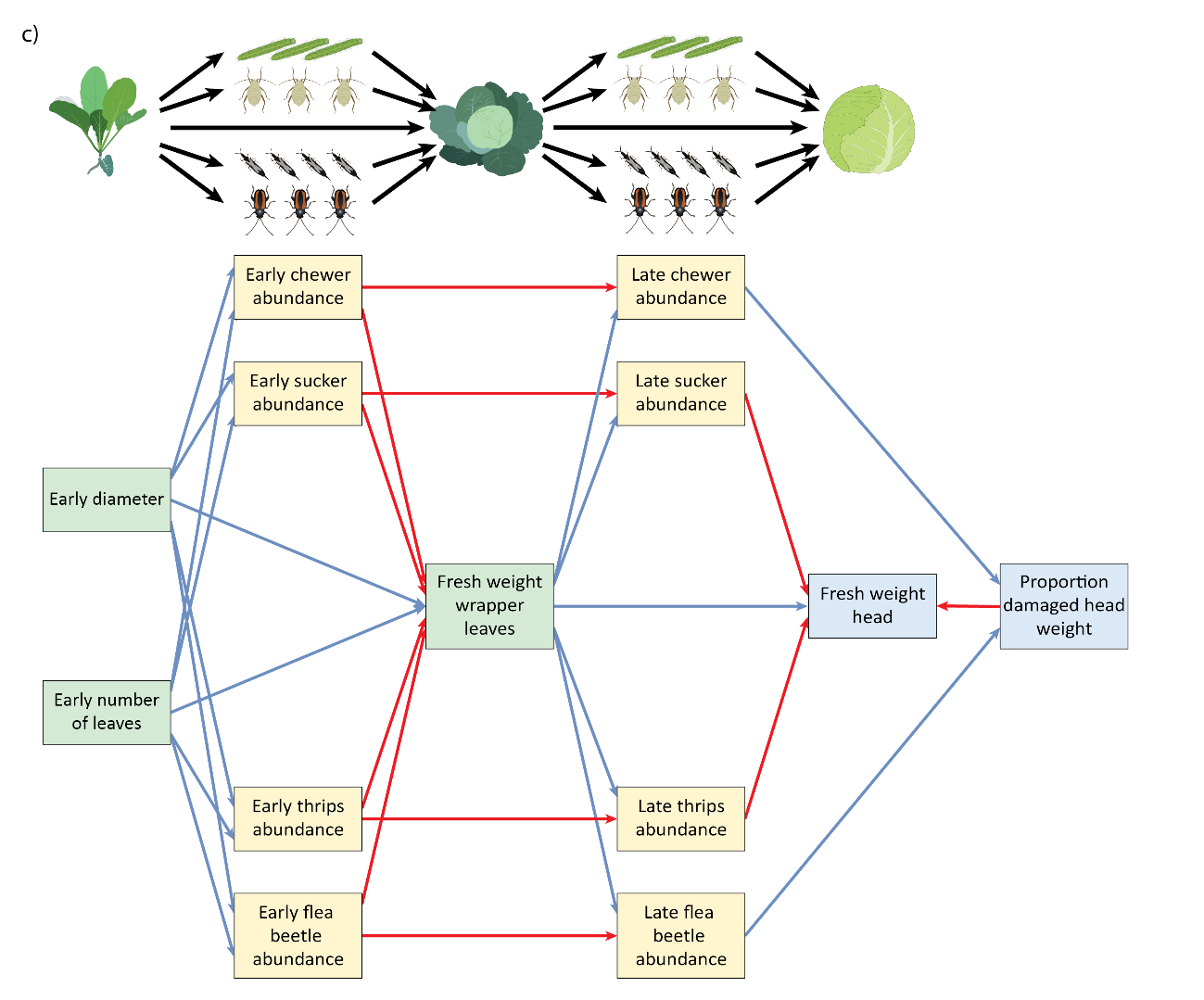

**Fig. S1. Hypothesized effects of plant size and herbivore variables on damage and fresh weight of the cabbage head.** These diagrams show the hypothesized structural equation models to test hypotheses on the relationship of plant size, herbivore abundance and cabbage fresh weight. Included in these models were plant variables for early plant size (diameter and number of leaves) and late plant size (fresh weight wrapper leaves) and **(a)** no further herbivore variables, or **(b)** early and late herbivore abundance and richness, and damage (proportion damaged head weight), or **(c)** early and late leaf chewer, phloem sucker, thrips and flea beetle abundance, and damage. Arrow color indicates the sign of the parameter estimate (blue = positive, red = negative, black = not significant). Green squares indicate plant variables related to plant growth, yellow squares indicate herbivore variables and blue squares indicate plant variables related to crop quantity and quality.

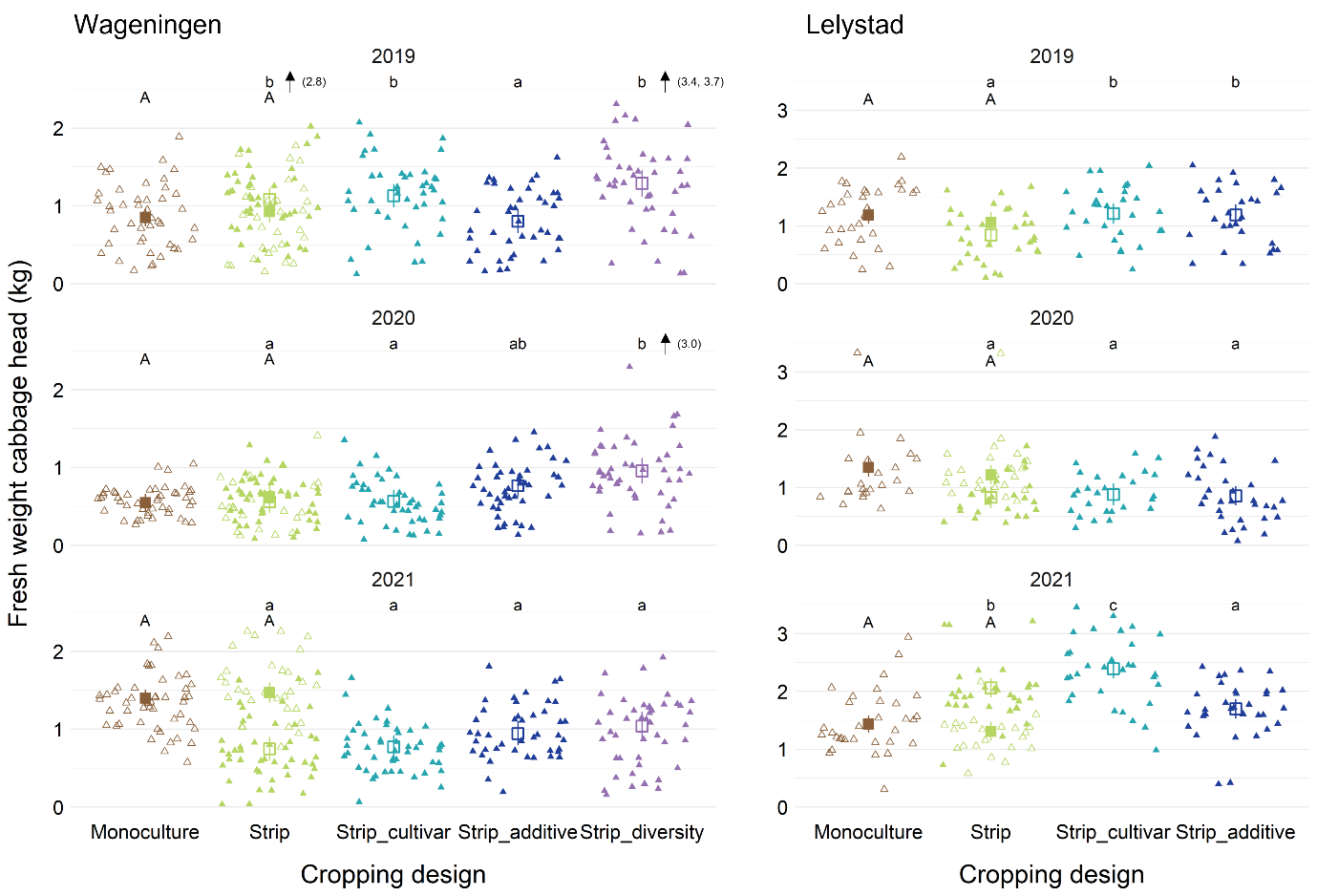

**Fig. S2. Effect of cropping system on fresh weight of the cabbage head.** Cabbage plants were grown in five treatments: monoculture, strip cropping of one cultivar of cabbage with one cultivar of wheat or oat (Strip), strip cropping of two cultivars of cabbage and with two cultivars of wheat or oat (Strip_cultivar), strip cropping of one cultivar of cabbage with a mixture of wheat or oat and faba bean (Strip_additive), and strip cropping of cabbage with wheat or oat, barley, pumpkin, grass-clover and potato including legumes in the grassy crops and two cultivars per crop (Strip_diversity, only at Wageningen). Colors indicate different crop configurations. Results are separated for the three years (rows) and two locations (columns). Each triangle indicates a single cabbage head. Because the reference field in 2021 showed marked differences with the other fields, we used two models: (1) monoculture versus Strip in the reference field (observations included in this analysis are indicated as open triangles) and (2) the strip configurations in the other fields (observations included in this analysis are indicated as filled triangles). The capital letters indicate significant differences between monoculture and strip cropping, the small letters indicate differences among strip configurations. The squares indicate estimated means and the vertical line within these shows the 95% confidence interval. Arrows indicate outliers and the values in brackets to the right indicate the values of these outliers.

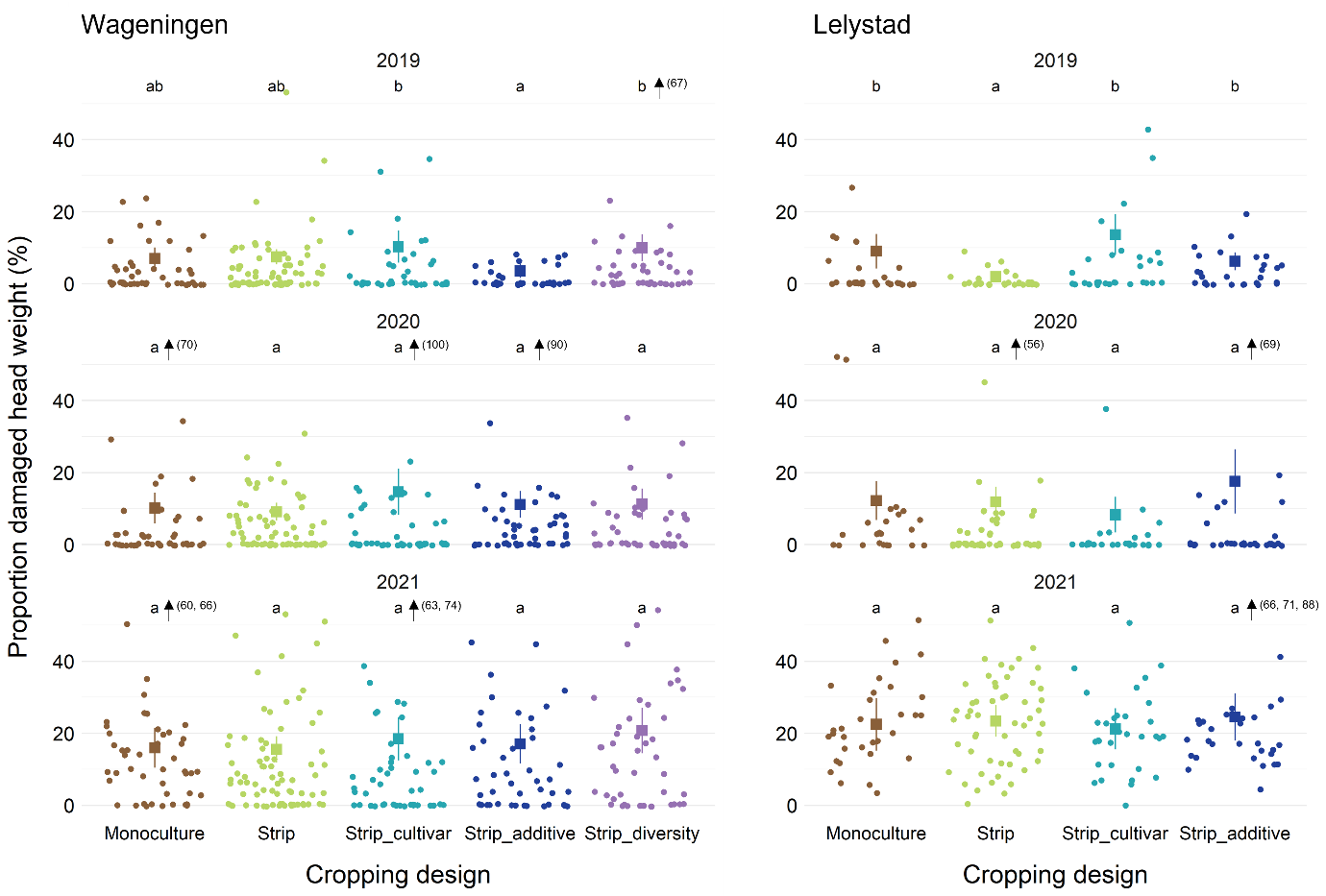
**Fig. S3. Effect of cropping system on proportion damaged head weight.** Cabbage plants were grown in five treatments: monoculture, strip cropping of one cultivar of cabbage with one cultivar of wheat or oat (Strip), strip cropping of two cultivars of cabbage with two cultivars of wheat or oat (Strip_cultivar), strip cropping of one cultivar of cabbage with a mixture of wheat or oat and faba bean (Strip_additive), and strip cropping of cabbage with wheat or oat, barley, pumpkin, grass-clover and potato including legumes in the grassy crops and two cultivars per crop (Strip_diversity, only at Wageningen). Results are separated for the three years (rows) and two locations (columns). Colors indicate different crop configurations. Each dot indicates a single cabbage head. Compact letter display was used to indicate significant differences between crop configurations within combinations of years and locations. The squares indicate estimated means and the vertical line around these shows the 95% confidence interval. Arrows indicate outliers and the values in brackets to the right indicate the values of these outliers.

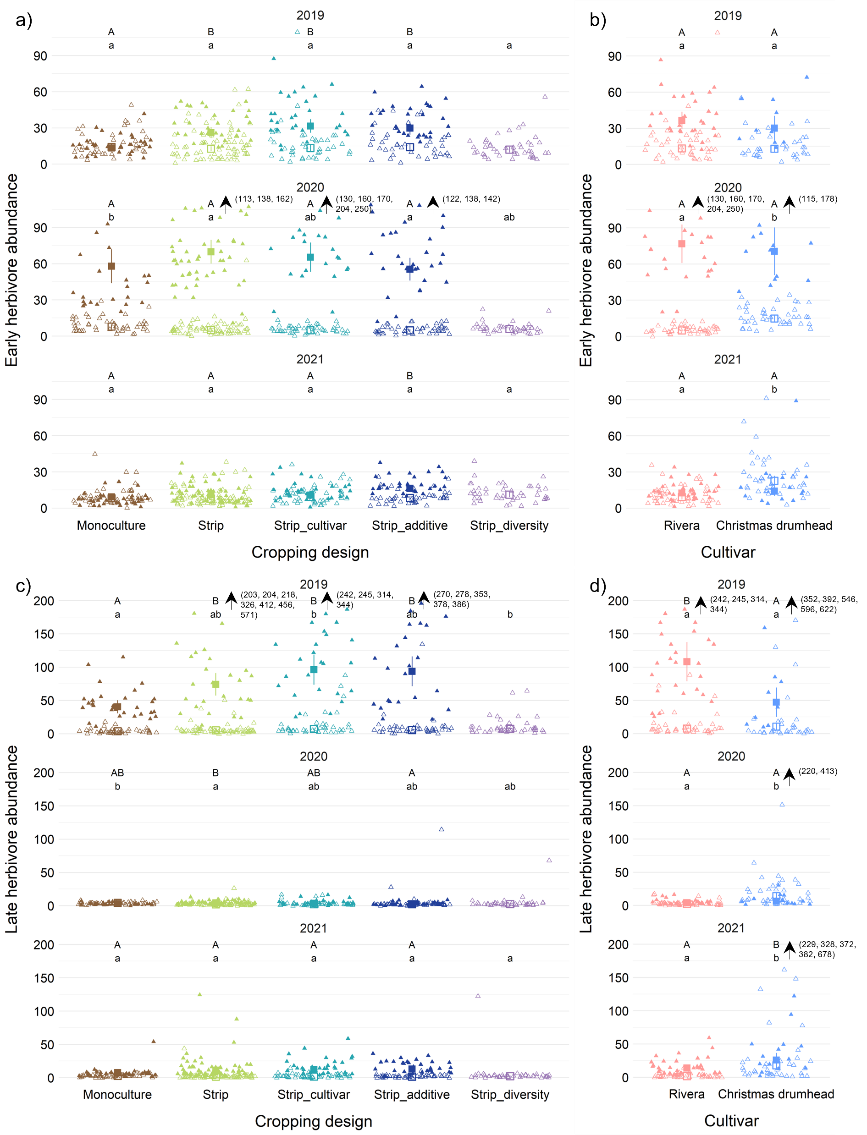

**Fig. S4. Effect of cropping system (a, c) and cultivar within Strip_ cultivar (b, d) on early (a, b) and late (c, d) herbivore abundance.** Cabbage plants were grown in five treatments: monoculture, strip cropping of one cultivar of cabbage with one cultivar of wheat or oat (Strip), strip cropping of two cultivars of cabbage with two cultivars of wheat or oat (Strip_cultivar), strip cropping of one cultivar of cabbage with a mixture of wheat or oat and faba bean (Strip_additive), and strip cropping of cabbage with wheat or oat, barley, pumpkin, grass-clover and potato including legumes in the grassy crops and two cultivars per crop (Strip_diversity, only at Wageningen). Results are separated for the three years (rows). Colors indicate different crop configurations (a, c) or different cultivars (b, d). Open triangles indicate single cabbage heads from Wageningen, closed triangles from Lelystad. Compact letter display was used to indicate significant differences per location and year, where the small letters indicate differences at Wageningen and the capital letters indicate difference at Lelystad. The squares indicate estimated means and the vertical line around these shows the 95% confidence interval. Arrows indicate outliers and the values in brackets to the right indicate the values of these outliers.

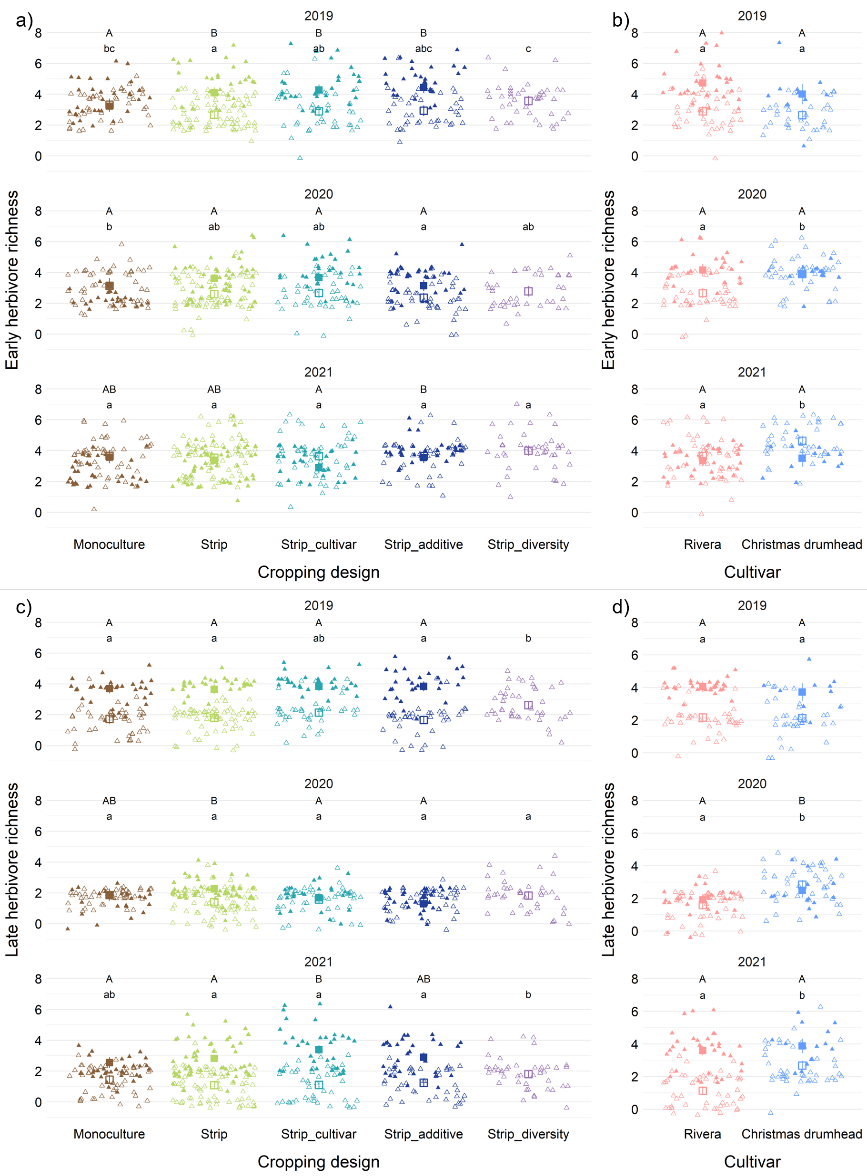

**Fig. S5. Effect of cropping system (a, c) and cultivar within Strip_ cultivar (b, d) on early (a, b) and late (c, d) herbivore richness.** Cabbage plants were grown in five treatments: monoculture, strip cropping of one cultivar of cabbage with one cultivar of wheat or oat (Strip), strip cropping of two cultivars of cabbage with two cultivars of wheat or oat (Strip_cultivar), strip cropping of one cultivar of cabbage with a mixture of wheat or oat and faba bean (Strip_additive), and strip cropping of cabbage with wheat or oat, barley, pumpkin, grass-clover and potato including legumes in the grassy crops and two cultivars per crop (Strip_diversity, only at Wageningen). Results are separated for the three years (rows). Colors indicate different crop configurations (a, c) or different cultivars (b, d). Open triangles indicate single cabbage heads from Wageningen, closed triangles from Lelystad. Compact letter display was used to indicate significant differences per location and year, where the small letters indicate differences at Wageningen and the capital letters indicate difference at Lelystad. The squares indicate estimated means and the vertical line around these shows the 95% confidence interval.

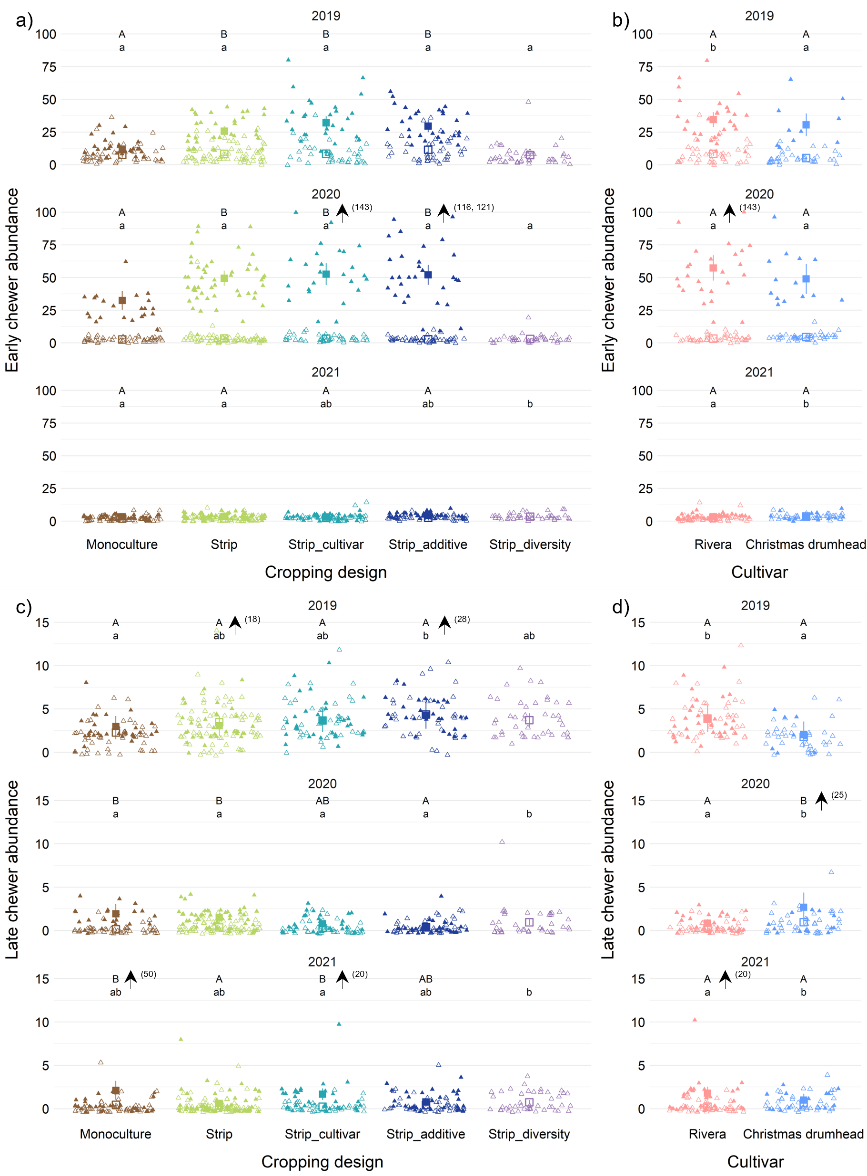

**Fig. S6. Effect of cropping system (a, c) and cultivar within Strip_ cultivar (b, d) on early (a, b) and late (c, d) leaf chewer abundance.** Cabbage plants were grown in five treatments: monoculture, strip cropping of one cultivar of cabbage with one cultivar of wheat or oat (Strip), strip cropping of two cultivars of cabbage with two cultivars of wheat or oat (Strip_cultivar), strip cropping of one cultivar of cabbage with a mixture of wheat or oat and faba bean (Strip_additive), and strip cropping of cabbage with wheat or oat, barley, pumpkin, grass-clover and potato including legumes in the grassy crops and two cultivars per crop (Strip_diversity, only at Wageningen). Results are separated for the three years (rows). Colors indicate different crop configurations (a, c) or different cultivars (b, d). Open triangles indicate single cabbage heads from Wageningen, closed triangles from Lelystad. Compact letter display was used to indicate significant differences per location and year, where the small letters indicate differences at Wageningen and the capital letters indicate difference at Lelystad. The squares indicate estimated means and the vertical line around these shows the 95% confidence interval. Arrows indicate outliers and the values in brackets to the right indicate the values of these outliers.

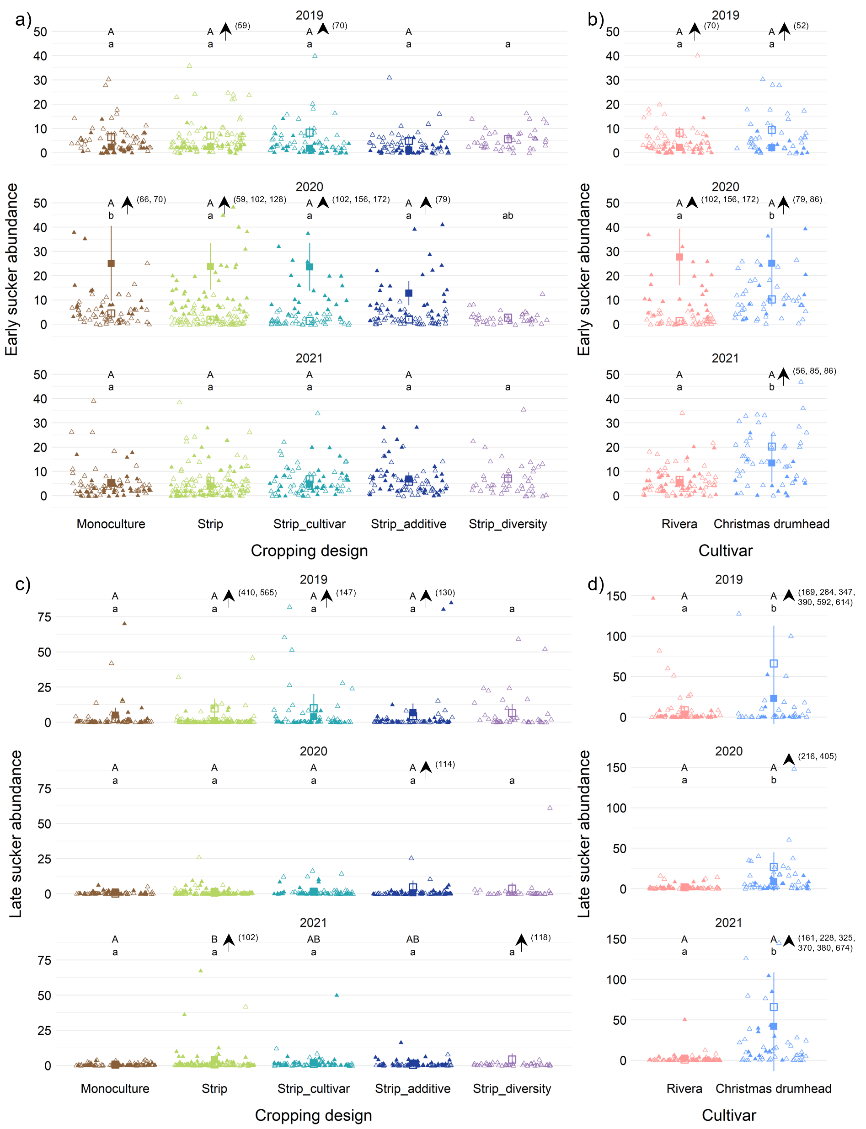

**Fig. S7. Effect of cropping systems (a, c) and cultivar within Strip_ cultivar (b, d) on early (a, b) and late (c, d) phloem sucker abundance.** Cabbage plants were grown in five treatments: monoculture, strip cropping of one cultivar of cabbage with one cultivar of wheat or oat (Strip), strip cropping of two cultivars of cabbage with two cultivars of wheat or oat (Strip_cultivar), strip cropping of one cultivar of cabbage with a mixture of wheat or oat and faba bean (Strip_additive), and strip cropping of cabbage with wheat or oat, barley, pumpkin, grass-clover and potato including legumes in the grassy crops and two cultivars per crop (Strip_diversity, only at Wageningen). Results are separated for the three years (rows). Colors indicate different crop configurations (a, c) or different cultivars (b, d). Open triangles indicate single cabbage heads from Wageningen, closed triangles from Lelystad. Compact letter display was used to indicate significant differences per location and year, where the small letters indicate differences at Wageningen and the capital letters indicate difference at Lelystad. The squares indicate estimated means and the vertical line around these shows the 95% confidence interval. Arrows indicate outliers and the values in brackets to the right indicate the values of these outliers.

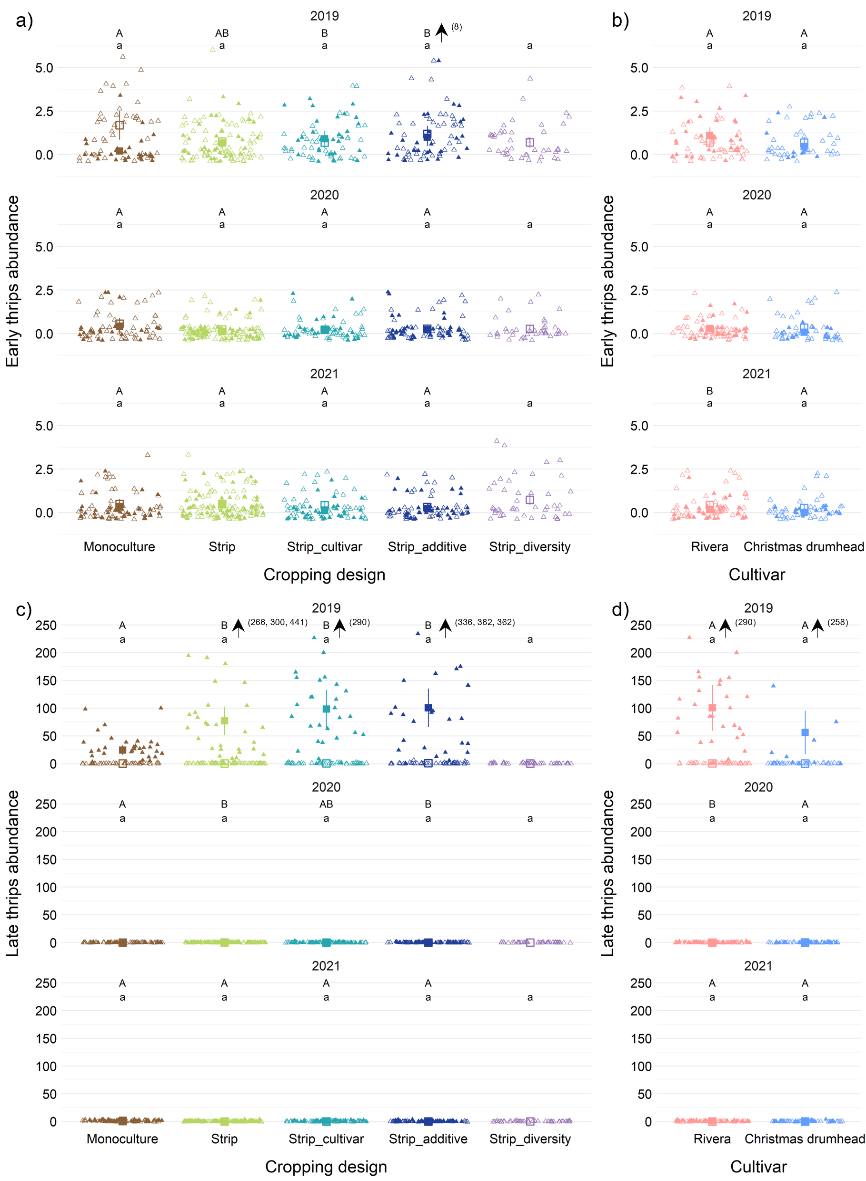

**Fig. S8. Effect of cropping systems (a, c) and cultivar within Strip_ cultivar (b, d) on early (a, b) and late (c, d) thrips abundance.** Cabbage plants were grown in five treatments: monoculture, strip cropping of one cultivar of cabbage with one cultivar of wheat or oat (Strip), strip cropping of two cultivars of cabbage with two cultivars of wheat or oat (Strip_cultivar), strip cropping of one cultivar of cabbage with a mixture of wheat or oat and faba bean (Strip_additive), and strip cropping of cabbage with wheat or oat, barley, pumpkin, grass-clover and potato including legumes in the grassy crops and two cultivars per crop (Strip_diversity, only at Wageningen). Results are separated for the three years (rows). Colors indicate different crop configurations (a, c) or different cultivars (b, d). Open triangles indicate single cabbage heads from Wageningen, closed triangles from Lelystad. Compact letter display was used to indicate significant differences per location and year, where the small letters indicate differences at Wageningen and the capital letters indicate difference at Lelystad. The squares indicate estimated means and the vertical line around these shows the 95% confidence interval. Arrows indicate outliers and the values in brackets to the right indicate the values of these outliers.

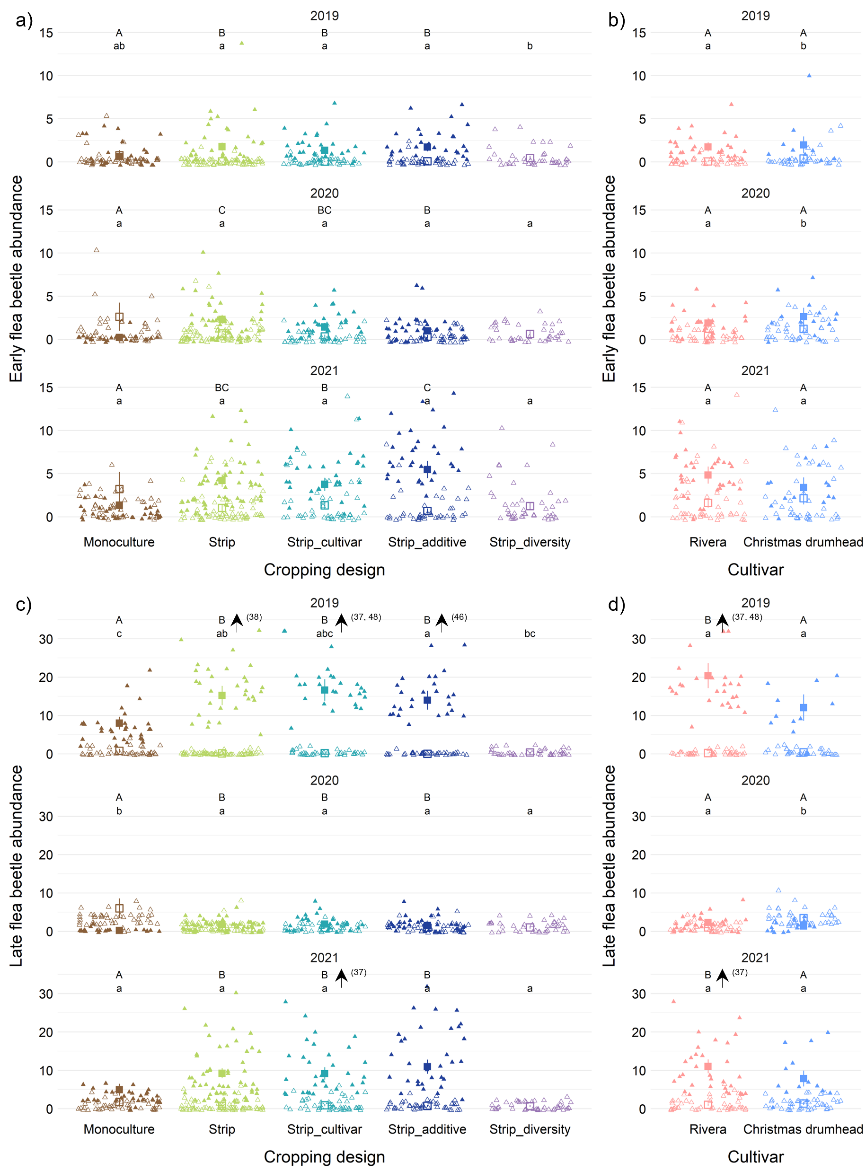

**Fig. S9. Effect of cropping systems (a, c) and cultivar within Strip_ cultivar (b, d) on early (a, b) and late (c, d) flea beetle abundance.** Cabbage plants were grown in five treatments: monoculture, strip cropping of one cultivar of cabbage with one cultivar of wheat or oat (Strip), strip cropping of two cultivars of cabbage with two cultivars of wheat or oat (Strip_cultivar), strip cropping of one cultivar of cabbage with a mixture of wheat or oat and faba bean (Strip_additive), and strip cropping of cabbage with wheat or oat, barley, pumpkin, grass-clover and potato including legumes in the grassy crops and two cultivars per crop (Strip_diversity, only at Wageningen). Results are separated for the three years (rows). Colors indicate different crop configurations (a, c) or different cultivars (b, d). Open triangles indicate single cabbage heads from Wageningen, closed triangles from Lelystad. Compact letter display was used to indicate significant differences per location and year, where the small letters indicate differences at Wageningen and the capital letters indicate difference at Lelystad. The squares indicate estimated means and the vertical line around these shows the 95% confidence interval. Arrows indicate outliers and the values in brackets to the right indicate the values of these outliers.

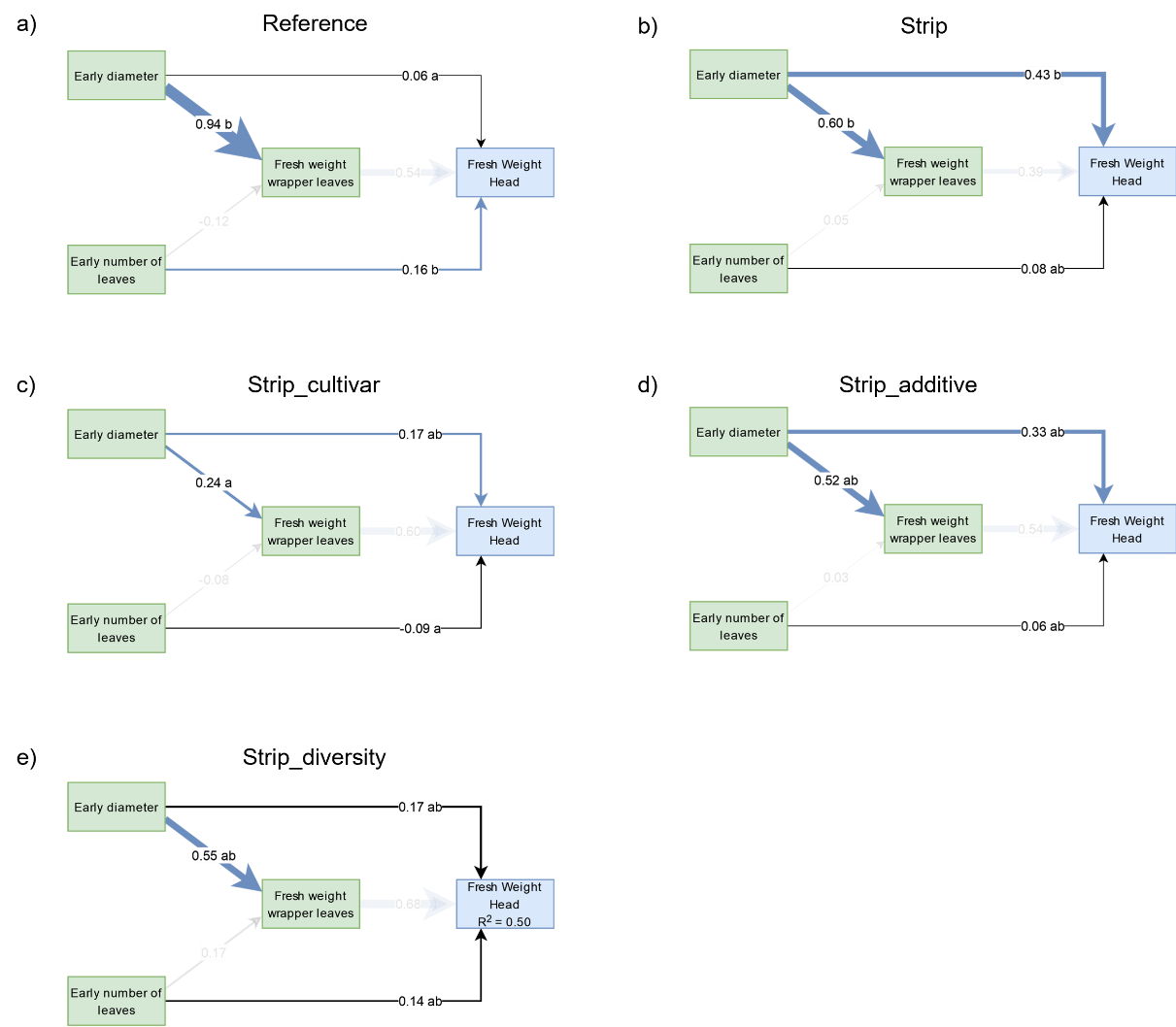

**Fig. S10. Comparison of pSEM models without herbivores per cropping system treatment.** Cabbage plants were grown in five treatments: monoculture, strip cropping of one cultivar of cabbage with one cultivar of wheat or oat (Strip), strip cropping of two cultivars of cabbage with two cultivars of wheat or oat (Strip_cultivar), strip cropping of one cultivar of cabbage with a mixture of wheat or oat and faba bean (Strip_additive), and strip cropping of cabbage with wheat or oat, barley, pumpkin, grass-clover and potato including legumes in the grassy crops and two cultivars per crop (Strip_diversity, only at Wageningen). Standardized parameter estimates are given on each arrow and arrow width also indicates the size of the standardized parameter estimates. Arrow color indicates the sign of the parameter estimate (blue = positive, red = negative, black = not significant). Compact letter display was used to indicate significant differences for paths among crop configurations, where the paths among two crop configurations were considered significantly different when their confidence intervals did not overlap. Arrows with reduced opacity were not significantly different among all crop configurations. Green squares indicate plant variables related to plant growth and blue squares indicate plant variables related to crop quantity. A separation in direct and indirect effects of variables on fresh weight head and proportion damaged head weight are given in Table S22.

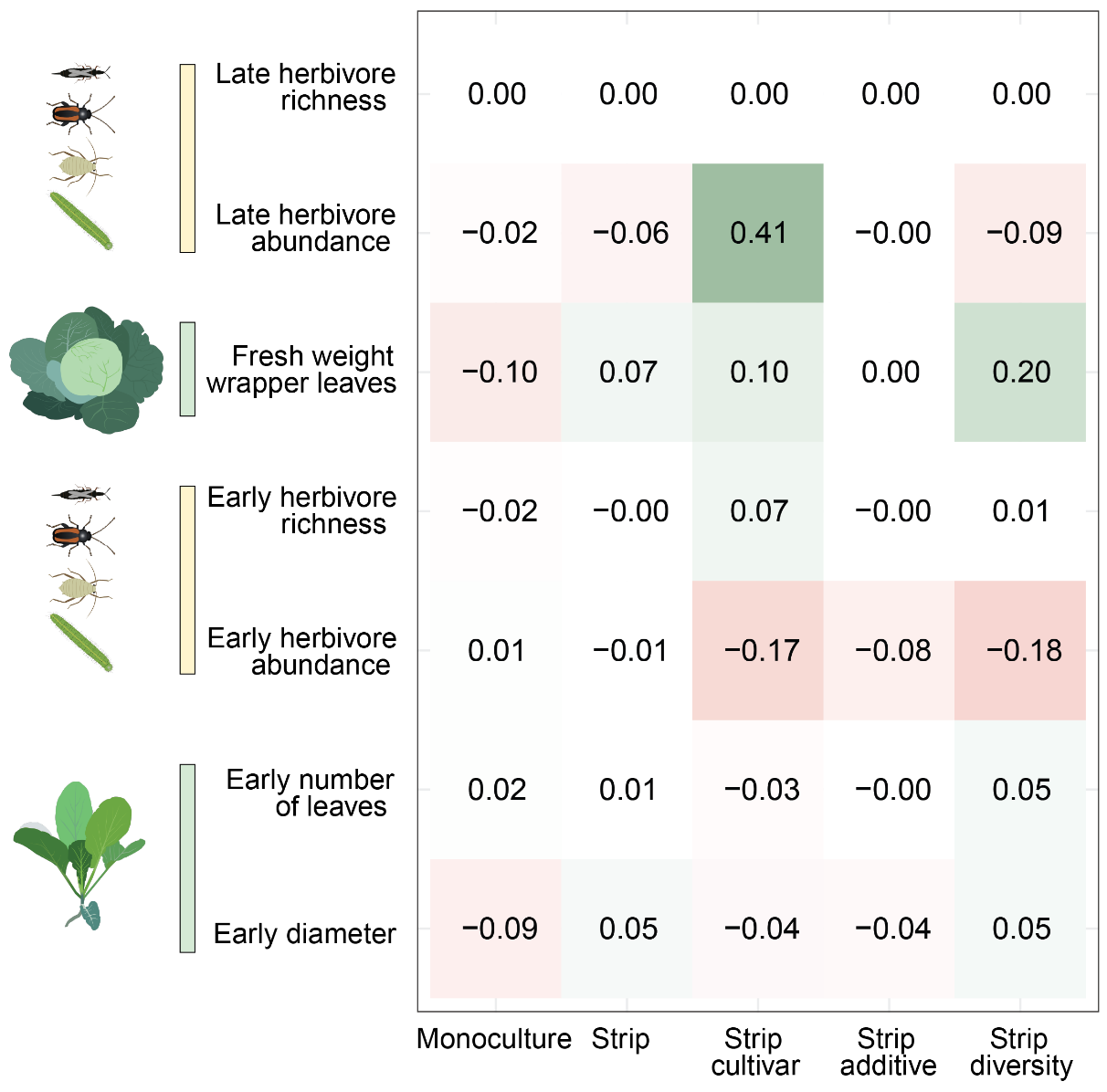

**Fig. S11. The effect of herbivore abundance and richness on damage of the cabbage head.** Total standardized effect of the predictor variables on proportion damaged head weight per crop configuration. Cabbage plants were grown in five treatments: monoculture, strip cropping of one cultivar of cabbage with one cultivar of wheat or oat (Strip), strip cropping of two cultivars of cabbage with two cultivars of wheat or oat (Strip_cultivar), strip cropping of one cultivar of cabbage with a mixture of wheat or oat and faba bean (Strip_additive), and strip cropping of cabbage with wheat or oat, barley, pumpkin, grass-clover and potato including legumes in the grassy crops and two cultivars per crop (Strip_diversity, only at Wageningen). The color gradient indicates the size and sign of the effect. A separation in direct and indirect effects of variables on fresh weight head and proportion damaged head weight are given in Tables S24.

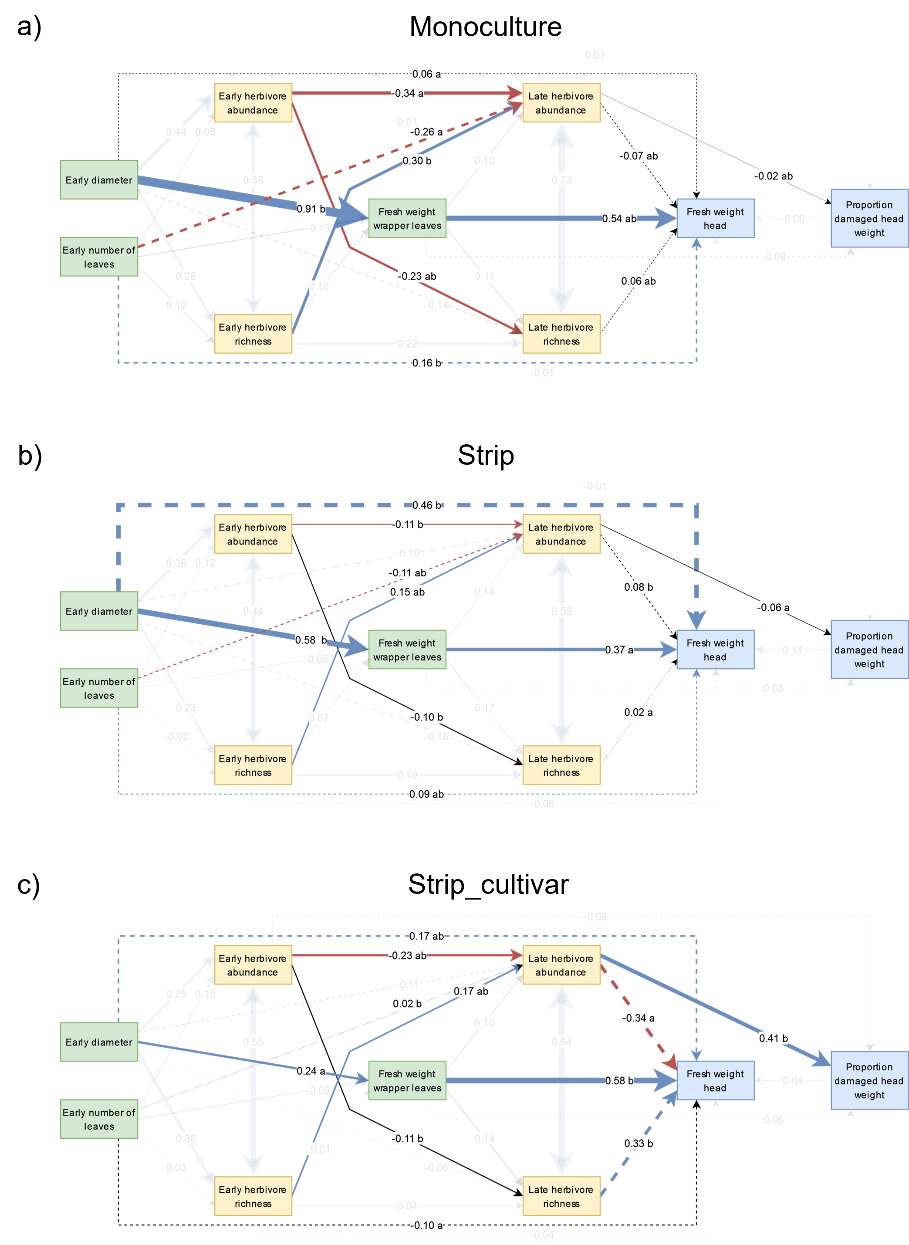

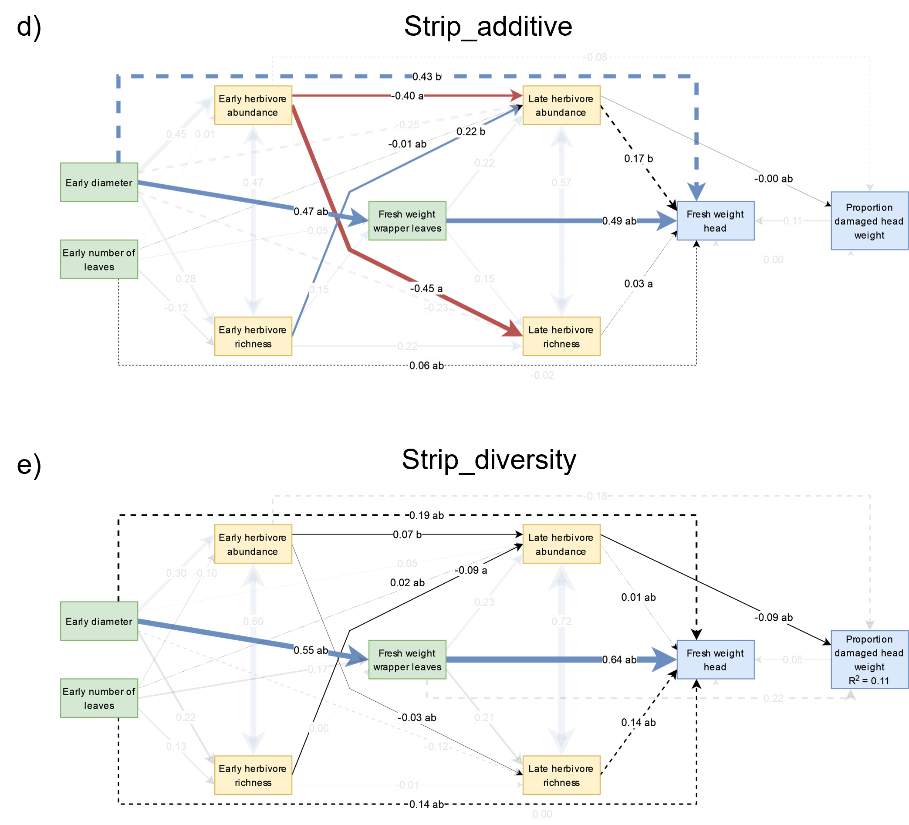

**Fig. S12. Comparison of pSEM models with herbivore abundance and richness per crop configuration.** Cabbage plants were grown in five treatments: monoculture, strip cropping of one cultivar of cabbage with one cultivar of wheat or oat (Strip), strip cropping of two cultivars of cabbage with two cultivars of wheat or oat (Strip_cultivar), strip cropping of one cultivar of cabbage with a mixture of wheat or oat and faba bean (Strip_additive), and strip cropping of cabbage with wheat or oat, barley, pumpkin, grass-clover and potato including legumes in the grassy crops and two cultivars per crop (Strip_diversity, only at Wageningen). Standardized parameter estimates are given on each arrow and arrow width also indicates the size of the standardized parameter estimates. Arrow color indicates the sign of the parameter estimate (blue = positive, red = negative, black = not significant). Compact letter display was used to indicate significant differences for paths among crop configurations, where the paths among two crop configurations were considered significantly different when their confidence intervals did not overlap. Arrows with reduced opacity were not significantly different among all crop configurations. Dotted arrows indicate paths that were included during the fitting process. Double headed arrows represent relations for which we could not presume a biologically meaningful causal relation, and which were included by their correlated error structure. Green squares indicate plant variables related to plant growth, yellow squares indicate herbivore variables and blue squares indicate plant variables related to crop quantity and quality. A separation in direct and indirect effects of variables on fresh weight head and proportion damaged head weight are given in Tables S23 & S24.

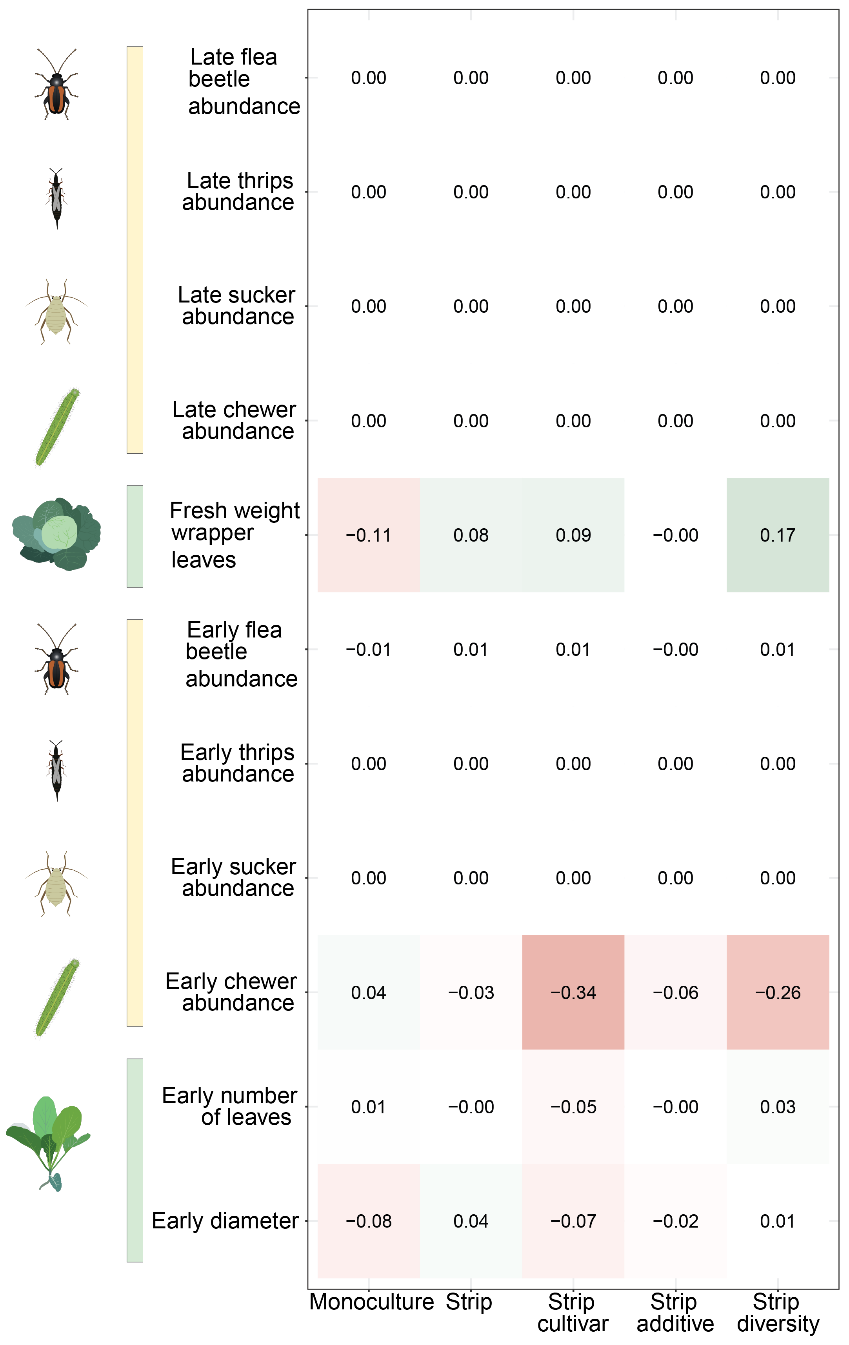

**Fig. S13. The effect of herbivore groups on damage of the cabbage head.** Total standardized effect of the predictor variables on proportion damaged head weight per crop configuration. Cabbage plants were grown in five treatments: monoculture, strip cropping of one cultivar of cabbage with one cultivar of wheat or oat (Strip), strip cropping of two cultivars of cabbage with two cultivars of wheat or oat (Strip_cultivar), strip cropping of one cultivar of cabbage with a mixture of wheat or oat and faba bean (Strip_additive), and strip cropping of cabbage with wheat or oat, barley, pumpkin, grass-clover and potato including legumes in the grassy crops and two cultivars per crop (Strip_diversity, only at Wageningen). The color gradient indicates the size and sign of the effect. A separation in direct and indirect effects of variables on fresh weight head and proportion damaged head weight are given in Tables S26.

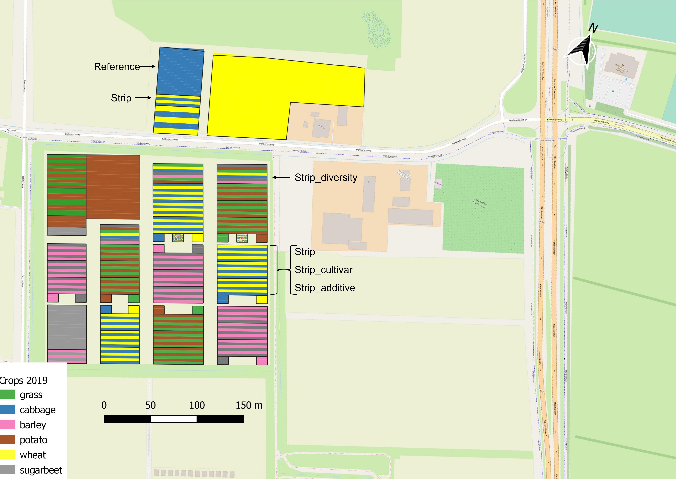

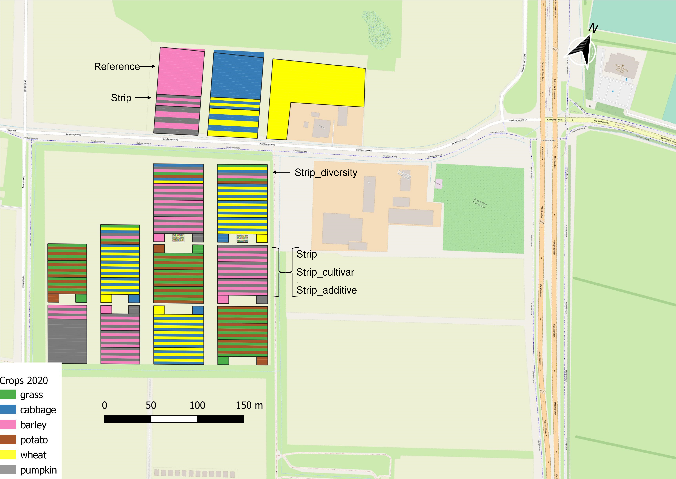

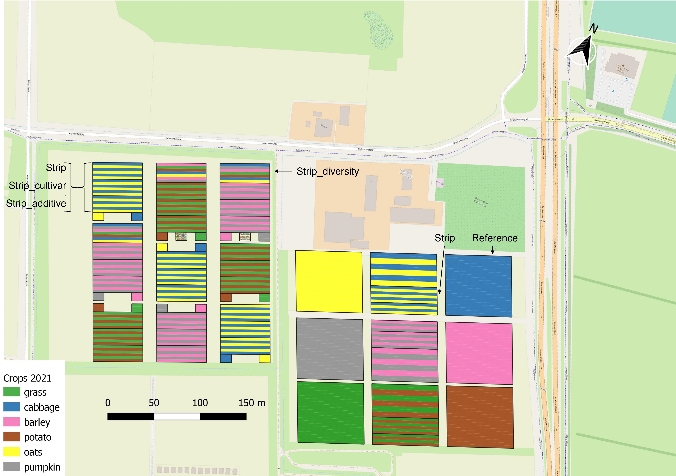

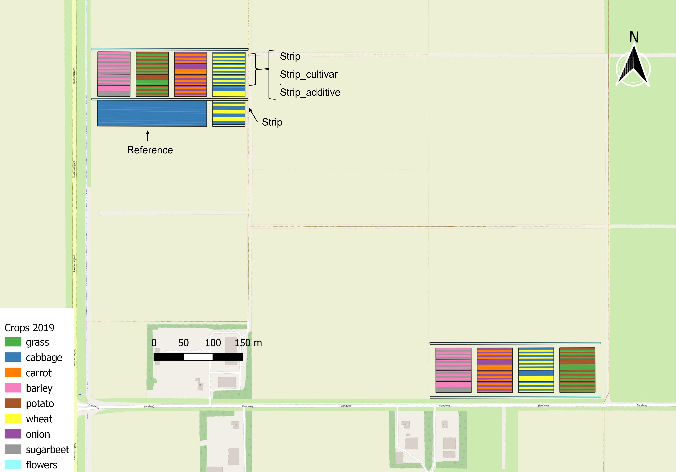

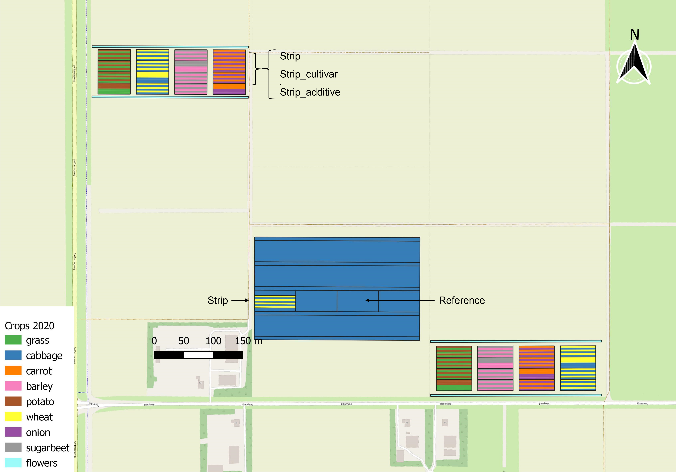

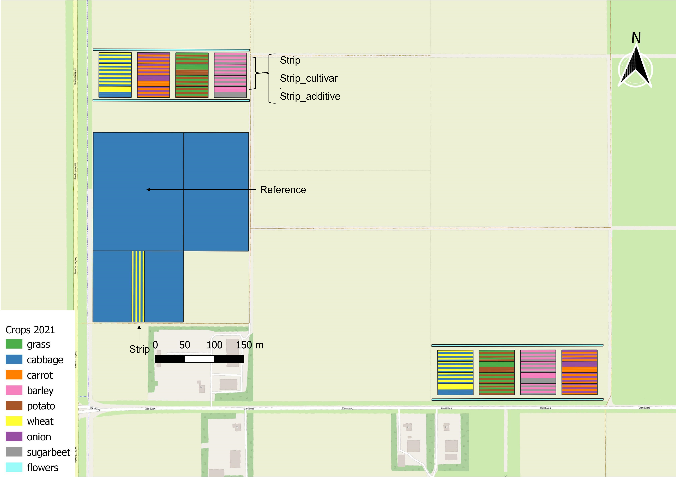

**Fig. S14. Maps of field lay-out of Wageningen (a, b, c) and Lelystad (d, e, f) per year.** Bright colours indicate the crops, dark outlines show the borders of different crop configurations. Cabbage plants were grown in five treatments: monoculture, strip cropping of one cultivar of cabbage with one cultivar of wheat or oat (Strip), strip cropping of two cultivars of cabbage with two cultivars of wheat or oat (Strip_cultivar), strip cropping of one cultivar of cabbage with a mixture of wheat or oat and faba bean (Strip_additive), and strip cropping of cabbage with wheat or oat, barley, pumpkin, grass-clover and potato including legumes in the grassy crops and two cultivars per crop (Strip_diversity, only at Wageningen).
