## Supplemental tables for "Herbivore prevalence poorly predicts yield in diverse cropping systems"

**Table S1.** Checklist of all flora included in some form in this study.

| **English name** | **Latin / Scientific name** |
| --- | --- |
| **Main Crops** | |
| White Cabbage | *Brassica oleracea* var. capitata |
| Wheat | *Triticum aestivum* |
| Oat | *Avena sativa* |
| Spelt | *Triticum spelta* |
| Pumpkin | *Cucurbita maxima* |
| Barley | *Hordeum vulgare* |
| Rye | *Secale cereale* |
| Potato | *Solanum tuberosum* |
| Italian rye-grass | *Lolium multiforum* |
| English rye-grass | *Lolium perenne* |
| Tall fescue | *Festuca arundinaceae* |
| Sugar beet | *Beta vulgaris* |
| Carrot | *Daucus carota* |
| Onion | *Allium cepa* |
| **Secondary crops** | |
| Alfalfa | *Medicago sativa* |
| Caraway | *Carum carvi* |
| Chicory | *Cichorium intybus* |
| Plantago | *Plantago sp.* |
| Red clover | *Trifolium pratense* |
| Faba bean | *Vicia faba* |
| Pea | *Pisum sativum* |
| **Green manure** | |
| Bristle oat | *Avena strigosa* |
| Buckwheat | *Fagopyrum esculentum* |
| Borage | *Borago officinalis* |
| Camelina | *Camelina sativa* |
| Flax | *Linum usitatissimum* |
| Fodder radish | *Raphanus sativus* subsp. *oleiferus* |
| Horseradish | *Armoracia rusticana* |
| Niger | *Guizotia abyssinica* |
| Phacelia | *Phacelia tanacetifolia* |
| Persian clover | *Trifolium resupinatum* |
| Serradella | *Ornithopus sativus* |
| Sunflower | *Helianthus annuus* |
| Vetch | *Vicia sp.* |
| White clover | *Trifolium repens* |
| White mustard | *Sinapis alba* |
| **Flower mixture (on the edges of Strip_additive in Wheat/Oat and Barley)** | |
| Camomile | *Matricaria chamomilla* |
| Common catchfly | *Silene gallica* |
| Corn marigold | *Glebionis segetum* |
| Cowcockle | *Vaccaria hispanica* |
| False mayweed | *Tripleurospermum maritimum* |
| Night-flowering catchfly | *Silene noctiflora* |
| Staggerweed | *Stachys arvensis* |
| **Perennial flower strips (only at Broekemahoeve)** | |
| Buckwheat | *Fagopyrum esculentum* |
| Common chicory | *Cichorium intybus* |
| Common poppy | *Papaver rhoeas* |
| Common yarrow | *Achillea millefolium* |
| Corn marigold | *Glebionis segetum* |
| Cornflower | *Centaurea cyanus* |
| Cow parsley | *Anthriscus sylvestris* |
| Dill | *Anethum graveolens* |
| Fennel | *Foeniculum vulgare* |
| Greater ammi | *Ammi majus* |
| Parsnip | *Pastinaca sativa* |
| Wild angelica | *Angelica sylvestris* |

**Table S2.** Checklist of all herbivores included in this study.

| **English name** | **Scientific name** | **Family** | **Order** | **Herbivore class** |
| --- | --- | --- | --- | --- |
| Diamondback moth | *Plutella xylostella* | Plutellidae | Lepidoptera | Chewer |
| Small white | *Pieris rapae* | Pieridae | Lepidoptera | Chewer |
| Large white | *Pieris brassicae* | Pieridae | Lepidoptera | Chewer |
| Cabbage moth | *Mamestra brassicae* | Noctuidae | Lepidoptera | Chewer |
| Silver Y | *Autographa gamma* | Noctuidae | Lepidoptera | Chewer |
| Armyworm | *Spodoptera sp.* | Noctuidae | Lepidoptera | Chewer |
| Garden pebble | *Evergestis forficalis* | Crambidae | Lepidoptera | Chewer |
| Turnip sawfly | *Athalia rosea* | Tenthredinidae | Hymenoptera | Chewer |
| Cabbage weevil | *Ceutorhynchus obstrictus* | Curculionidae | Coleoptera | Chewer |
| Turnip flea beetle | *Phyllotreta atra* | Chrysomelidae | Coleoptera | Flea beetle |
| Small striped flea beetle | *Phyllotreta undulata* | Chrysomelidae | Coleoptera | Flea beetle |
| Thrips | *Thrips sp.* | Thripidae | Thysanoptera | Thrips |
| Cabbage aphid | *Brevicoryne brassicae* | Aphididae | Hemiptera | Suckers |
| Green peach aphid | *Myzus persicae* | Aphididae | Hemiptera | Suckers |
| Red peach aphid | *Myzus persicae* | Aphididae | Hemiptera | Suckers |
| Black bean aphid | *Aphis fabae* | Aphididae | Hemiptera | Suckers |
| Cabbage whitefly | *Aleyrodes proletella* | Aleyrodidae | Hemiptera | Suckers |
| Cicada | *Auchenorryhyncha* | N/A | Hemiptera | Suckers |

**Table S3.** Major management practices performed in Lelystad (Broekemahoeve) over the course of the three years period for cabbage and wheat. Blackened cells indicate that the practice was done in this treatment. For wheat, “Reference” refers to the wheat strips that were placed next to the cabbage monoculture reference. For cabbage, “Reference” refers both to the monoculture reference and the strips on the same field. “Strip”, “Strip_cultivar” and “Strip_additive” refer to different strip cropping designs of cabbage and wheat with respectively no further additions (Strip), mix cropping of two cultivars of cabbage and wheat within the strips (Strip_cultivar) and mix gropping of faba beans within wheat (Strip_additive). Separate tables describe management practices for **(a)** white cabbage in 2019, **(b)** wheat in 2019, **(c)** white cabbage in 2020, **(d)** wheat in 2020, **(e)** white cabbage in 2021 and **(f)** wheat in 2021.

**Table S3a.** Management practices for white cabbage in 2019.

| Date | Reference | Strip | Strip_cultivar | Strip_additive | Management practice |
| --- | --- | --- | --- | --- | --- |
| Autumn, 2018 |  |  |  |  | **Fertilization** with farm-yard manure (25 t/ha) |
| End of March |  |  |  |  | **Fertilization** with slurry manure (35 m^3^/ha) |
| End of March |  |  |  |  | **Tillage** to loosen soil |
| 16-05-2019 |  |  |  |  | Mechanically **planting** Rivera cv. |
| 16-05-2019 +/- |  |  |  |  | Manually **planting** Christmas Drumhead cv. |
| From 23-05-2019 |  |  |  |  | **Weed control** by harrowing  (every 2-3 weeks until plant canopy closed) |
| From 30-05-2019 |  |  |  |  | **Weed control** by hoeing  (alternately with harrowing, until plant canopy closed) |
| September - November |  |  |  |  | Manual **harvest** |
| After harvest |  |  |  |  | **Crop residue destruction** using flail mower |
| After harvest |  |  |  |  | **Tillage** to loosen soil |

**Table S3b.** Management practices for wheat in 2019.

| Date | Strips  (Reference field) | Strip | Strip_cultivar | Strip_additive | Management practice |
| --- | --- | --- | --- | --- | --- |
| March |  |  |  |  | **Sowing** spring wheat (180 kg/ha) |
| April or May |  |  |  |  | **Fertilization** with slurry manure (30 m^3^/ha)iu |
| May - June |  |  |  |  | **Weed control** by harrowing (5 times) |
| May - June |  |  |  |  | **Weed control** by hoeing |
| 07-08-2019 |  |  |  |  | **Harvest** using combine harvester |
| After harvest |  |  |  |  | **Tillage** to loosen soil with spring tine cultivator |
| After harvest |  |  |  |  | **Sowing** green manure |

**Table S3c.** Management practices for white cabbage in 2020.

| Date | Reference | Strip | Strip_cultivar | Strip_additive | Management practice |
| --- | --- | --- | --- | --- | --- |
| End of 2018 |  |  |  |  | **Sowing** grass-clover |
| October, 2019 |  |  |  |  | **Sowing** green manure |
| 23-03-2020 |  |  |  |  | **Tillage** using flail mower |
| 26-03-2020 |  |  |  |  | **Tillage** using rotary tiller |
| 26-03-2020 |  |  |  |  | **Tillage** using spring tine cultivator |
| April-may |  |  |  |  | **Fertilization** with farm-yard manure (30 t/ha) |
| Before planting |  |  |  |  | **Fertilization** with slurry manure (35 m^3^/ha) |
| 18-05-2020 |  |  |  |  | Mechanically **planting** Rivera cv. |
| 20-05-2020 |  |  |  |  | Manually **planting** Christmas Drumhead cv. |
| May - August |  |  |  |  | **Weed control** by harrowing (2 times) |
| May - August |  |  |  |  | **Weed control** by hoeing (2-3 times) |
| May – June |  |  |  |  | **Irrigation** (10 – 20 mm, 1-2 times) |
| 19-10-2020 |  |  |  |  | Manual **harvest** |
| November |  |  |  |  | **Crop residue destruction** using flail mower |
| November |  |  |  |  | **Tillage** using disc harrow |
| November |  |  |  |  | **Tillage** using duck foot cultivator |

**Table S3d.** Management practices for wheat in 2020.

| Date | Strips  (Reference field) | Strip | Strip_cultivar | Strip_additive | Management practice |
| --- | --- | --- | --- | --- | --- |
| End of 2018 |  |  |  |  | **Sowing** grass-clover |
| August, 2019 |  |  |  |  | **Sowing** grass-clover |
| Before sowing |  |  |  |  | **Tillage** to loosen soil |
| March/April |  |  |  |  | **Sowing** spring wheat |
| March - May |  |  |  |  | **Weed control** by hoeing (2-3 times) |
| 10-08-2020 |  |  |  |  | **Harvest** using combine harvester |
| 08-09-2020 |  |  |  |  | **Harvest** using combine harvester |
| August |  |  |  |  | **Fertilization** with farm-yard manure (30 t/ha) |
| After harvest |  |  |  |  | **Sowing** green manure |

**Table S3e.** Management practices for white cabbage in 2021.

| Date | Reference | Strip | Strip_cultivar | Strip_additive | Management practice |
| --- | --- | --- | --- | --- | --- |
| 08-03-2021 |  |  |  |  | **Tillage** by milling |
| 01-04-2021 |  |  |  |  | **Fertilization** with chicken manure (6 t/ha) |
| 16-04-2021 |  |  |  |  | **Tillage** using spring-tine cultivator |
| 22-04-2021 |  |  |  |  | **Fertilization** with slurry manure (35 t/ha) |
| 20-05-2021 |  |  |  |  | Mechanically **planting** Rivera cv. |
| 26-05-2021 |  |  |  |  | Manually **planting** Christmas Drumhead cv. |
| 31-05-2021 |  |  |  |  | **Weed control** using spring-tine cultivator |
| 07-06-2021 |  |  |  |  | **Weed control** using goose foot cultivator |
| 14-06-2021 |  |  |  |  | **Fertilization** with organic plant fertilizer (400 kg/ha) |
| 15-06-2021 |  |  |  |  | **Weed control** by hoeing |
| 24-06-2021 |  |  |  |  | **Weed control** by hoeing |
| 12-07-2021 |  |  |  |  | **Weed control** by hoeing |
| 07-10-2021 +/- |  |  |  |  | Manual **harvest** |
| November |  |  |  |  | **Crop residue destruction** using flail mower |
| November |  |  |  |  | **Tillage** using disc harrow |
| November |  |  |  |  | **Tillage** using duck foot cultivator |

**Table S3f.** Management practices for wheat in 2021.

| Date | Strips  (Reference field) | Strip | Strip_cultivar | Strip_additive | Management practice |
| --- | --- | --- | --- | --- | --- |
| October, 2020 |  |  |  |  | **Sowing** winter wheat |
| 2^nd^ week of April +/- |  |  |  |  | **Sowing** spring wheat |
| 30-03-2021 |  |  |  |  | **Weed control** by hoeing |
| 01-04-2021 |  |  |  |  | **Fertilization** with chicken manure (6 t/ha) |
| 15-04-2021 |  |  |  |  | **Weed control** by hoeing |
| 21-05-2021 |  |  |  |  | **Weed control** by hoeing |
| 04-08-2021 |  |  |  |  | **Harvest** using combine harvester |
| 21-08-2021 |  |  |  |  | **Harvest** using combine harvester |
| 15-09-2021 +/- |  |  |  |  | **Sowing** green manure |

**Table S4.** Major management practices performed in Wageningen (Droevendaal) over the course of the three years period for cabbage and wheat. Blackened cells indicate that the practice was done in this treatment. “Reference” refers both to the monoculture reference and the strips on the same field. “Strip”, “Strip_cultivar”, “Strip_additive” and “Strip_diversity” refer to different strip cropping designs of cabbage and wheat with respectively no further additions (Strip), mix cropping of two cultivars of cabbage and wheat within the strips (Strip_cultivar), mix gropping of faba beans within wheat (Strip_additive) and strip cropping of six crops including at least two cultivars per crop and legumes within the cereal crops (Strip_diversity). When all the boxes in the column “Reference” were not blackened, then no monoculture reference was available for that crop that year. Separate tables describe management practices for the separate crops in **(a - g)** 2019, **(h – n)** 2020 and **(o – u)** 2021.

**Table S4a.** Management practices for white cabbage in 2019.

| Date | Reference | Strip | Strip_cultivar | Strip_additive | Strip_diveristy | Management practice |
| --- | --- | --- | --- | --- | --- | --- |
| 11-04-2019 |  |  |  |  |  | Mechanically mow Grass-clover of previous year |
| 30-04-2019 |  |  |  |  |  | **Fertilization** using farm-yard manure (20 t/ha) |
| 01-05-2019 |  |  |  |  |  | **Tillage** using rotary tiller |
| 21-05-2019 |  |  |  |  |  | **Tillage** using rotary tiller |
| 23-05-2019 |  |  |  |  |  | Mechanical **planting** of Rivera cv. |
| 24-05-2019 |  |  |  |  |  | **Irrigation** using sprinkler (5 mm) |
| 25-05-2019 |  |  |  |  |  | **Irrigation** using pivot (18 mm) |
| 26-05-2019 |  |  |  |  |  | Manual **planting** of Christmas Drumhead cv. |
| 05-06-2019 |  |  |  |  |  | **Weed control** using fixed-tine cultivator |
| 06-06-2019 |  |  |  |  |  | **Irrigation** using sprinkler (5 mm) |
| 26-06-2019 |  |  |  |  |  | **Weed control** using fixed-tine cultivator and finger weeder |
| 31-06-2019 |  |  |  |  |  | **Weed control** using fixed-tine cultivator and finger weeder |
| 08-08-2019 |  |  |  |  |  | **Fertilization** using organic plant fertilizer (1.5 t/ha) |
| 21-08-2019 |  |  |  |  |  | **Weed control** using duck foot cultivator |
| 13-09-2019 |  |  |  |  |  | Manual **weed control** |
| 15-10-2019 |  |  |  |  |  | Manual **harvest** (1^st^ round) |
| 04-11-2019 |  |  |  |  |  | Manual **harvest** (2^nd^ round) |
| 08-11-2019 |  |  |  |  |  | **Crop residue destruction** using flail mower and incorporation with rotary tiller |
| 23-11-2019 |  |  |  |  |  | **Sowing** winter barley |

**Table S4b.** Management practices for wheat in 2019.

| Date | Reference | Strip | Strip_cultivar | Strip_additive | Strip_diveristy | Management practice |
| --- | --- | --- | --- | --- | --- | --- |
| 05-10-2018 |  |  |  |  |  | **Sowing** winter wheat (and faba bean) |
| 26-10-2018 |  |  |  |  |  | **Weed control** using duck foot cultivator |
| 12-04-2019 |  |  |  |  |  | **Fertilization** using slurry manure (15 m^3^/ha) |
| 16-04-2019 |  |  |  |  |  | **Sowing** spring wheat |
| 01-05-2019 |  |  |  |  |  | **Weed control** using duck foot cultivator |
| 14-05-2019 |  |  |  |  |  | **Weed control** using duck foot cultivator |
| 20-05-2019 |  |  |  |  |  | **Weed control** using duck foot cultivator |
| 25-05-2019 |  |  |  |  |  | **Irrigation** using pivot (18 mm) |
| 31-05-2019 |  |  |  |  |  | **Weed control** using duck foot cultivator |
| 30-06-2019 |  |  |  |  |  | **Harvest** using combine harvester |
| 17-07-2019 |  |  |  |  |  | Manual **weed control** |
| 24-07-2019 |  |  |  |  |  | **Harvest** using combine harvester |
| 31-07-2019 |  |  |  |  |  | **Tillage** using rotary tiller and disc harrower |
| 05-08-2019 |  |  |  |  |  | **Tillage** using rotary tiller and disc harrower |
| 18-09-2019 |  |  |  |  |  | **Sowing** green manure |

**Table S4c.** Management practices for sugar beet in 2019.

| Date | Reference | Strip | Strip_cultivar | Strip_additive | Strip_diveristy | Management practice |
| --- | --- | --- | --- | --- | --- | --- |
| Early spring |  |  |  |  |  | **Tillage** using rotary tiller |
| 30-04-2019 |  |  |  |  |  | **Fertilization** using slurry manure (35 m^3^/ha) |
| 06-05-2019 |  |  |  |  |  | **Sowing** sugar beet |
| 25-05-2019 |  |  |  |  |  | **Irrigation** using pivot (18 mm) |
| June or July |  |  |  |  |  | **Tillage** using rotary tiller to remove sugar beet which failed |
| 15-11-2019 |  |  |  |  |  | **Sowing** grass (and clover) |

**Table S4d.** Management practices for barley in 2019.

| Date | Reference | Strip | Strip_cultivar | Strip_additive | Strip_diveristy | Management practice |
| --- | --- | --- | --- | --- | --- | --- |
| 17-12-2018 |  |  |  |  |  | **Sowing** rye (green manure) |
| 30-04-2019 |  |  |  |  |  | **Fertilization** using slurry manure (15 m^3^/ha) |
| 02-05-2019 |  |  |  |  |  | **Tillage** using rotary tiller |
| 02-05-2019 |  |  |  |  |  | **Sowing** spring barley (and peas) |
| 22-05-2019 |  |  |  |  |  | **Weed control** using spring-tine cultivator |
| 25-05-2019 |  |  |  |  |  | **Irrigation** using pivot (18 mm) |
| 03-06-2019 |  |  |  |  |  | **Weed control** using spring-tine cultivator |
| 20-08-2019 |  |  |  |  |  | **Harvest** using combine harvester |
| 08-11-2019 |  |  |  |  |  | **Sowing** green manure |

**Table S4e.** Management practices for potato in 2019.

| Date | Reference | Strip | Strip_cultivar | Strip_additive | Strip_diveristy | Management practice |
| --- | --- | --- | --- | --- | --- | --- |
| 10-04-2019 |  |  |  |  |  | Manual **harvest** of leek |
| 29-04-2019 |  |  |  |  |  | **Fertilization** with chop-drop grass |
| 30-04-2019 |  |  |  |  |  | **Fertilization** using farm-yard manure (35 t/ha) |
| 01-05-2019 |  |  |  |  |  | **Tillage** with rotary tiller |
| 03-05-2019 |  |  |  |  |  | **Planting** potatoes |
| 25-05-2019 |  |  |  |  |  | **Irrigation** with pivot (18 mm) |
| 04-06-2019 |  |  |  |  |  | **Ridging** |
| 12-06-2019 |  |  |  |  |  | **Ridging** |
| 17-06-2019 |  |  |  |  |  | **Ridging** |
| 04-09-2019 |  |  |  |  |  | **Burning** |
| 12-09-2019 |  |  |  |  |  | **Burning** |
| 18-09-2019 |  |  |  |  |  | Mechanical **harvest** |
| 27-09-2019 |  |  |  |  |  | **Tillage** with subsoiler |
| 27-09-2019 |  |  |  |  |  | **Tillage** with harrow tiller |
| 28-10-2019 |  |  |  |  |  | **Sowing** winter wheat (and faba bean) |

**Table S4f.** Management practices for grass in 2019.

| Date | Reference | Strip | Strip_cultivar | Strip_additive | Strip_diveristy | Management practice |
| --- | --- | --- | --- | --- | --- | --- |
| Spring, 2018 |  |  |  |  |  | **Sowing** grass (and clover) |
| 30-04-2019 |  |  |  |  |  | **Fertilization** using slurry manure (3 t/ha) |
| 06-05-2019 |  |  |  |  |  | **Tillage** with rotary tiller |
| 13-05-2019 |  |  |  |  |  | **Re-sowing** grass (and clover) |
| 25-05-2019 |  |  |  |  |  | **Irrigation** with pivot (18 mm) |
| 30-05-2019 |  |  |  |  |  | **Harvest** by mowing |
| 12-07-2019 |  |  |  |  |  | **Harvest** by mowing |
| 09-10-2019 |  |  |  |  |  | **Harvest** by mowing |
| 29-10-2019 |  |  |  |  |  | **Harvest** by mowing |

**Table S4g.** Management practices for white cabbage in 2020.

| Date | Reference | Strip | Strip_cultivar | Strip_additive | Strip_diveristy | Management practice |
| --- | --- | --- | --- | --- | --- | --- |
| 01-05-2020 |  |  |  |  |  | Mechanically mow grass of previous year |
| 12-05-2020 |  |  |  |  |  | **Fertilization** using manure (35 t/ha) |
| 12-05-2020 |  |  |  |  |  | **Fertilization** using organic plant ferilizer (1.5 t/ha) |
| 13-05-2020 |  |  |  |  |  | **Tillage** using rotary tiller |
| 25-05-2020 |  |  |  |  |  | Mechanical **planting** of Rivera cv. |
| 26-05-2020 |  |  |  |  |  | Manual **planting** of Christmas Drumhead cv. |
| June - August |  |  |  |  |  | **Weed control** using duck foot cultivator (and finger weeder) (done several times) |
| September |  |  |  |  |  | Manual **weed control** (one round of hand-weeding) |
| 15-10-2020 |  |  |  |  |  | Manual **harvest** (1^st^ round) |
| 10-11-2020 |  |  |  |  |  | Manual **harvest** (2^nd^ round) |
| 26-11-2020 |  |  |  |  |  | Manual **harvest** (3^rd^ round) |
| 30-11-2020 |  |  |  |  |  | **Crop residue destruction** using flail mower and incorporation with rotary tiller |
| 30-11-2020 |  |  |  |  |  | **Liming** (2500 kg/ha Miramag) |
| 30-11-2020 |  |  |  |  |  | **Sowing** winter barley (but failed, so resown in April 2021) |

**Table S4h.** Management practices for wheat in 2020.

| Date | Reference | Strip | Strip_cultivar | Strip_additive | Strip_diveristy | Management practice |
| --- | --- | --- | --- | --- | --- | --- |
| 28-10-2019 |  |  |  |  |  | **Sowing** winter wheat (and faba bean) |
| 07-02-2020 |  |  |  |  |  | **Weed control** using spring-tine cultivator |
| 07-03-2020 |  |  |  |  |  | **Weed control** using spring-tine cultivator |
| 19-03-2020 |  |  |  |  |  | **Weed control** using spring-tine cultivator |
| 19-03-2020 |  |  |  |  |  | **Sowing** of wildflower mix |
| 25-03-2020 |  |  |  |  |  | **Weed control** using spring-tine cultivator |
| 26-03-2020 |  |  |  |  |  | **Fertilization** using slurry manure (20 m^3^/ha) |
| 15-04-2020 |  |  |  |  |  | **Weed control** using spring-tine cultivator |
| 04-05-2020 |  |  |  |  |  | **Weed control** using spring-tine cultivator |
| 12-05-2020 |  |  |  |  |  | **Weed control** using spring-tine cultivator |
| 31-07-2020 |  |  |  |  |  | **Harvest** using combine harvester |
| 21-08-2020 |  |  |  |  |  | **Sowing** green manure |
| 30-11-2020 |  |  |  |  |  | **Liming** (2500 kg/ha Miramag) |

**Table S4i.** Management practices for sugar beet / pumpkin in 2020.

| Date | Reference | Strip | Strip_cultivar | Strip_additive | Strip_diveristy | Management practice |
| --- | --- | --- | --- | --- | --- | --- |
| 18-09-2019 |  |  |  |  |  | **Sowing** green manure |
| 15-11-2019 |  |  |  |  |  | **Sowing** grass |
| 26-03-2020 |  |  |  |  |  | **Fertilization** using slurry manure (35 m^3^/ha) |
| 09-04-2020 |  |  |  |  |  | **Tillage** using rotary tiller |
| 10-04-2020 |  |  |  |  |  | **Sowing** sugar beet |
| 29-04-2020 |  |  |  |  |  | **Tillage** using spring-tine cultivator |
| 25-05-2020 |  |  |  |  |  | **Sowing** pumpkin |
| 16-06-2020 |  |  |  |  |  | **Re-sowing** pumpkin (crop failure) |
| July and August |  |  |  |  |  | **Weed control** using duck foot cultivator and fixed tine cultivator (several times) |
| 02-10-2020 |  |  |  |  |  | Manual **harvest** |
| 15-10-2020 |  |  |  |  |  | **Tillage** using disc harrower |
| 05-11-2020 |  |  |  |  |  | **Tillage** using rotary tiller |
| 03-05-2021 |  |  |  |  |  | **Sowing** grass (and clover) |

**Table S4j.** Management practices for barley in 2020.

| Date | Reference | Strip | Strip_cultivar | Strip_additive | Strip_diveristy | Management practice |
| --- | --- | --- | --- | --- | --- | --- |
| 23-11-2019 |  |  |  |  |  | **Sowing** winter barley and peas |
| 19-03-2020 |  |  |  |  |  | **Tillage** using spring-tine cultivator |
| 25-03-2020 |  |  |  |  |  | **Tillage** using spring-tine cultivator |
| 26-03-2020 |  |  |  |  |  | **Fertilization** using slurry manure (15 m^3^/ha) |
| 15-04-2020 |  |  |  |  |  | **Tillage** using spring-tine cultivator |
| 20-07-2020 |  |  |  |  |  | **Harvest** using combine harvester |
| 23-10-2020 |  |  |  |  |  | **Crop residue destruction** using flail mower |

**Table S4k.** Management practices for potato in 2020.

| Date | Reference | Strip | Strip_cultivar | Strip_additive | Strip_diveristy | Management practice |
| --- | --- | --- | --- | --- | --- | --- |
| 21-04-2020 |  |  |  |  |  | **Fertilization** using farm-yard manure (35 t/ha) |
| 26-04-2020 |  |  |  |  |  | **Tillage** with rotary tiller |
| 28-04-2020 |  |  |  |  |  | **Planting** potatoes |
| 15-05-2020 |  |  |  |  |  | **Fertilization** with chop-crop grass (18 t/ha) |
| 23-05-2020 |  |  |  |  |  | **Ridging** |
| 24-06-2020 |  |  |  |  |  | **Fertilization** with chop-crop grass (18 t/ha) |
| 29-07-2020 |  |  |  |  |  | **Burning** (1^st^ time) |
| 31-07-2020 |  |  |  |  |  | **Burning** (2^nd^ time) |
| 25-08-2020 |  |  |  |  |  | Mechanical **harvest** |
| 26-09-2020 |  |  |  |  |  | **Liming** (2500 kg/ha Miramag) |
| 23-10-2020 |  |  |  |  |  | **Tillage** with rotary tiller |
| 05-11-2020 |  |  |  |  |  | **Tillage** with duck foot cultivator |
| 10-11-2020 |  |  |  |  |  | **Sowing** spelt (and faba bean) |

**Table S4l.** Management practices for grass in 2020.

| Date | Reference | Strip | Strip_cultivar | Strip_additive | Strip_diveristy | Management practice |
| --- | --- | --- | --- | --- | --- | --- |
| 15-11-2019 |  |  |  |  |  | **Sowing** grass (and clover) |
| 26-03-2020 |  |  |  |  |  | **Fertilization** using slurry manure (10 m^3^/ha) |
| Throughout season |  |  |  |  |  | **Harvest** by mowing (several times) |
| 26-09-2020 |  |  |  |  |  | **Liming** (2500 kg/ha Miramag) |

**Table S4m.** Management practices for white cabbage in 2021.

| Date | Reference | Strip | Strip_cultivar | Strip_additive | Strip_diveristy | Management practice |
| --- | --- | --- | --- | --- | --- | --- |
| 06-04-2021 |  |  |  |  |  | **Tillage** using rotary tiller |
| 21-04-2021 |  |  |  |  |  | **Fertilization** using farm-yard manure (15 t/ha) |
| 12-05-2021 |  |  |  |  |  | **Tillage** using rotary tiller |
| 18-05-2021 |  |  |  |  |  | **Fertilization** using slurry manure (10 m^3^/ha) |
| 20-05-2021 |  |  |  |  |  | Mechanical **planting** of Rivera cv. |
| 20-05-2021 |  |  |  |  |  | Manual **planting** of Christmas Drumhead cv. |
| 07-06-2021 |  |  |  |  |  | **Weed control** using duck foot cultivator |
| 10-06-2021 |  |  |  |  |  | **Fertilization** using organic plant fertilizer (1 t/ha) |
| 14-06-2021 |  |  |  |  |  | **Weed control** using duck foot cultivator and finger weeder |
| 16-06-2021 |  |  |  |  |  | Manual **weed control** |
| 28-06-2021 |  |  |  |  |  | Manual **weed control** |
| 27-08-2021 |  |  |  |  |  | **Fertilization** using organic plant fertilizer (0.5 t/ha) |
| 05-10-2021 |  |  |  |  |  | Manual **harvest** (1^st^ round) |
| 02-11-2021 |  |  |  |  |  | Manual **harvest** (2^nd^ round) |
| 05-11-2021 |  |  |  |  |  | **Crop residue destruction** using flail mower and incorporation with rotary tiller |
| 10-11-2021 |  |  |  |  |  | **Sowing** winter barley |

**Table S4n.** Management practices for wheat in 2021.

| Date | Reference | Strip | Strip_cultivar | Strip_additive | Strip_diveristy | Management practice |
| --- | --- | --- | --- | --- | --- | --- |
| 10-11-2020 |  |  |  |  |  | **Sowing** spelt (and faba bean) |
| 03-03-2021 |  |  |  |  |  | **Tillage** using rotary tiller to remove Spelt which failed |
| 02-04-2021 |  |  |  |  |  | **Fertilization** using slurry manure (25 t/ha) |
| 12-04-2021 |  |  |  |  |  | **Tillage** using fixed-tine cultivator |
| 14-04-2021 |  |  |  |  |  | **Sowing** oat |
| 23-04-2021 |  |  |  |  |  | **Sowing** of wildflower border |
| 28-04-2021 |  |  |  |  |  | **Weed control** using spring-tine cultivator |
| 10-05-2021 |  |  |  |  |  | **Weed control** using spring-tine cultivator |
| 17-05-2021 |  |  |  |  |  | **Weed control** using spring-tine cultivator |
| 31-05-2021 |  |  |  |  |  | **Weed control** using spring-tine cultivator |
| 25-08-2021 |  |  |  |  |  | **Harvest** using combine harvester |
| 26-08-2021 |  |  |  |  |  | **Crop residue destruction** using flail mower |
| 27-08-2021 |  |  |  |  |  | **Tillage** using rotary tiller |
| 31-08-2021 |  |  |  |  |  | **Sowing** green manure |

**Table S4o.** Management practices for pumpkin in 2021.

| Date | Reference | Strip | Strip_cultivar | Strip_additive | Strip_diveristy | Management practice |
| --- | --- | --- | --- | --- | --- | --- |
| 21-08-2020 |  |  |  |  |  | **Sowing** green manure |
| 21-04-2021 |  |  |  |  |  | **Fertilization** using farm-yard manure (15 t/ha) |
| 12-05-2021 |  |  |  |  |  | **Tillage** using duck foot cultivator |
| 02-06-2021 |  |  |  |  |  | **Tillage** using rotary tiller |
| 09-06-2021 |  |  |  |  |  | **Sowing** pumpkin |
| 10-06-2021 |  |  |  |  |  | **Weed control** using duck foot cultivator |
| 25-06-2021 |  |  |  |  |  | **Weed control** using duck foot cultivator |
| 30-06-2021 |  |  |  |  |  | Manual **weed control** |
| 07-07-2021 |  |  |  |  |  | **Weed control** using duck foot cultivator |
| 08-07-2021 |  |  |  |  |  | **Fertilization** using organic plant fertilizer (0.81 t/ha) and lucerne pellets (0.54 t/ha) |
| 10-07-2021 |  |  |  |  |  | **Weed control** using duck foot cultivator |
| 15-09-2021 |  |  |  |  |  | Manual **harvest** |
| 17-09-2021 |  |  |  |  |  | **Tillage** using rotary tiller |
| 21-09-2021 |  |  |  |  |  | **Sowing** grass (and clover) |

**Table S4p.** Management practices for barley in 2021.

| Date | Reference | Strip | Strip_cultivar | Strip_additive | Strip_diveristy | Management practice |
| --- | --- | --- | --- | --- | --- | --- |
| 30-11-2020 |  |  |  |  |  | **Sowing** winter barley and peas (but failed, so resown in April 2021) |
| 03-03-2021 |  |  |  |  |  | **Tillage** using duck foot cultivator |
| 02-04-2021 |  |  |  |  |  | **Fertilization** using slurry manure (20 m^3^ / ha) |
| 12-04-2021 |  |  |  |  |  | **Sowing** spring barley and peas |
| 28-04-2021 |  |  |  |  |  | **Weed control** using spring-tine cultivator |
| 10-05-2021 |  |  |  |  |  | **Weed control** using spring-tine cultivator |
| 17-05-2021 |  |  |  |  |  | **Weed control** using spring-tine cultivator |
| 31-05-2021 |  |  |  |  |  | **Weed control** using spring-tine cultivator |
| 24-08-2021 |  |  |  |  |  | **Harvest** using combine harvester |
| 26-08-2021 |  |  |  |  |  | **Crop residue destruction** using flail mower |
| 27-08-2021 |  |  |  |  |  | **Tillage** using rotary tiller |
| 31-08-2021 |  |  |  |  |  | **Sowing** green manure |

**Table S4q.** Management practices for potato in 2021.

| Date | Reference | Strip | Strip_cultivar | Strip_additive | Strip_diveristy | Management practice |
| --- | --- | --- | --- | --- | --- | --- |
| 21-04-2021 |  |  |  |  |  | **Fertilization** using farm-yard manure (25 t/ha) |
| 26-04-2021 |  |  |  |  |  | **Tillage** with rotary tiller |
| 28-04-2021 |  |  |  |  |  | **Sowing** potatoes |
| 07-05-2021 |  |  |  |  |  | **Fertilization** with chop-drop grass (20 t/ha) |
| 01-06-2021 |  |  |  |  |  | **Ridging** |
| 08-06-2021 |  |  |  |  |  | **Ridging** |
| 23-06-2021 |  |  |  |  |  | **Ridging** |
| 19-07-2021 |  |  |  |  |  | **Burning** |
| 23-07-2021 |  |  |  |  |  | **Burning** |
| 30-08-2021 |  |  |  |  |  | Mechanical **harvest** |
| 07-09-2021 |  |  |  |  |  | **Tillage** using duck foot cultivator |
| 07-09-2021 |  |  |  |  |  | **Liming** (2500 kg/ha Miramag) |
| 09-09-2021 |  |  |  |  |  | **Sowing** green manure |

**Table S4r.** Management practices for grass in 2021.

| Date | Reference | Strip | Strip_cultivar | Strip_additive | Strip_diveristy | Management practice |
| --- | --- | --- | --- | --- | --- | --- |
| 03-03-2021 |  |  |  |  |  | **Weed control** using spring-tine cultivator |
| 12-04-2021 |  |  |  |  |  | **Weed control** using spring-tine cultivator |
| 03-05-2021 |  |  |  |  |  | **Sowing** grass-clover |
| 24-06-2021 |  |  |  |  |  | **Harvest** by mowing |
| 06-08-2021 |  |  |  |  |  | **Harvest** by mowing |
| 07-09-2021 |  |  |  |  |  | **Liming** (2500 kg/ha Miramag) |
| 14-09-2021 |  |  |  |  |  | **Harvest** by mowing |

**Table S5.** Ground cover and plant mixtures throughout the three years field experiment for Wageningen (Droevendaal). Merged cells indicate the period of a certain ground cover. For the monoculture reference each column indicates either cabbage or wheat at the reference field. For the other cropping designs each column depicts the same strips over the three years. “Unknown” means that this information could not be found within the cropping management logs. Crop cultivars and seeding/planting densities are given whenever known.

| **Reference** | | | | | | |
| --- | --- | --- | --- | --- | --- | --- |
| **Year** | **Month** | **Cabbage Reference 2019** | **Wheat Reference 2019 / Cabbage Reference 2020** | **Wheat Reference 2020** | **Cabbage Reference 2021** | **Wheat Reference 2021** |
| 2019 | Jan | **Oat** | **Oat** | **Oat** | **No crop / bare soil** | **No crop / bare soil** |
|  | Feb |  |  |  |  |  |
|  | Mar |  |  |  | **Summer oat** | **Summer oat** |
|  | Apr |  |  |  |  |  |
|  | May | **No crop / bare soil** | **Spring wheat**  Harenda cv.  (200 kg/ha) | **Spring wheat**  Harenda cv.  (200 kg/ha) |  |  |
|  | Jun | **White cabbage**  Rivera cv.  (30k plants/ha) |  |  |  |  |
|  | Jul |  | **Spring wheat**  Stubble left | **Spring wheat**  Stubble left |  |  |
|  | Aug |  | **No crop / bare soil** | **No crop / bare soil** |  |  |
|  | Sep |  |  |  | **Green manure**  Fodder radish  Bristle oat  Yellow mustard | **Green manure**  Fodder radish  Bristle oat  Yellow mustard |
|  | Oct |  |  |  |  |  |
|  | Nov | **No crop / bare soil** |  | **Winter wheat**  Talent KWS cv.  (200 kg/ha) |  |  |
|  | Dec | **Winter barley**  Cassiopee cv.  (100 kg/ha) |  |  |  |  |
| 2020 | Jan |  |  |  |  |  |
|  | Feb |  |  |  |  |  |
|  | Mar |  |  |  | **Summer barley** | **Summer barley** |
|  | Apr |  |  |  |  |  |
|  | May |  | **White cabbage**  Rivera cv.  (30k plants/ha) |  |  |  |
|  | Jun |  |  |  |  |  |
|  | Jul |  |  |  |  |  |
|  | Aug |  |  | **Winter wheat**  Stubble left | **Summer barley**  Reshoots from before | **No crop / bare soil** |
|  | Sep |  |  |  |  |  |
|  | Oct |  |  | **Oat** |  |  |
|  | Nov |  |  |  |  | **Winter spelt**  **(failed)** |
|  | Dec |  |  |  |  |  |
| 2021 | Jan |  |  |  |  |  |
|  | Feb |  |  |  |  |  |
|  | Mar |  |  |  |  |  |
|  | Apr |  |  |  |  |  |
|  | May |  |  |  |  | **Oat**  Symphony cv.  (200 kg/ha) |
|  | Jun |  |  |  | **White cabbage**  Rivera cv.  (30k plants/ha) |  |
|  | Jul |  |  |  |  |  |
|  | Aug |  |  |  |  |  |
|  | Sep |  |  |  |  | **Green manure**  Rye  (100 kg/ha) |
|  | Oct |  |  |  |  |  |
|  | Nov |  |  |  | **Winter barley**  Cassiopee cv.  (180 kg/ha) |  |
|  | Dec |  |  |  |  |  |

| **Strip** | | | | | | | | | |
| --- | --- | --- | --- | --- | --- | --- | --- | --- | --- |
| **Year** | **Month** | **Strip 1** | **Strip 2** |  | **Strip 3** | **Strip 4** |  | **Strip 5** | **Strip 6** |
| 2019 | Jan | **Leek**  Pluston cv.  (110,000 plants/ha) | **Italian rye grass**  Danergo cv.  (35 kg/ha) |  | **Winter wheat**  Kelvin cv.  (200 kg/ha) | **Italian rye grass**  Danergo cv.  (35 kg/ha) |  | **No crop / bare soil** | **Rye**  Ruben cv.  (100 kg/ha) |
|  | Feb |  |  |  |  |  |  |  |  |
|  | Mar |  |  |  |  |  |  |  |  |
|  | Apr |  |  |  |  |  |  |  |  |
|  | May | **Potato**  Agria cv.  (2750 kg/ha) |  |  |  | **No crop / bare soil** |  | **Sugar beet**  **(failed)** | **Spring barley**  Irina cv.  (200 kg/ha) |
|  | Jun |  |  |  |  | **White cabbage**  Rivera cv.  (30k plants/ha) |  | **No crop / bare soil** |  |
|  | Jul |  |  |  |  |  |  |  |  |
|  | Aug |  |  |  | **No crop / bare soil** |  |  |  |  |
|  | Sep |  |  |  |  |  |  |  | **No crop / bare soil** |
|  | Oct | **No crop / bare soil** |  |  | **Green manure**  Buckwheat  Cv 1  (40 kg/ha) |  |  |  |  |
|  | Nov | **Winter wheat**  Kelvin cv.  (200 kg/ha) |  |  |  | **No crop / bare soil** |  | **Italian rye grass**  Danergo cv.  (35 kg/ha) | **Green manure**  Horseradish  Radetzky cv.  (40 kg/ha) |
|  | Dec |  |  |  |  | **Winter barley**  Cassiopee cv.  (100 kg/ha) |  |  |  |
| 2020 | Jan |  |  |  |  |  |  |  |  |
|  | Feb |  |  |  |  |  |  |  |  |
|  | Mar |  |  |  |  |  |  |  |  |
|  | Apr |  |  |  |  |  |  |  |  |
|  | May |  | **White cabbage**  Rivera cv.  (30k plants/ha) |  | **Sugar beet**  **(failed)** |  |  |  | **Potato**  Agria cv.  (2750 kg/ha) |
|  | Jun |  |  |  | **Pumpkin**  Kaori kuri cv.  (18k seeds/ha) |  |  |  |  |
|  | Jul |  |  |  |  |  |  |  |  |
|  | Aug | **No crop / bare soil** |  |  |  | **Winter barley**  Stubble left |  |  |  |
|  | Sep | **Green manure**  Bristle oat  (60 kg/ha) |  |  |  |  |  |  | **No crop / bare soil** |
|  | Oct |  |  |  | **No crop / bare soil** |  |  |  |  |
|  | Nov |  |  |  |  |  |  |  | **Spelt**  **(failed)**  (200 kg/ha) |
|  | Dec |  | **No crop / bare soil** |  |  |  |  |  |  |
| 2021 | Jan |  |  |  |  |  |  |  |  |
|  | Feb |  |  |  |  |  |  |  |  |
|  | Mar |  |  |  |  |  |  |  | **No crop / bare soil** |
|  | Apr |  | **Spring barley**  RGT planet cv.  (200 kg/ha) |  |  |  |  | **No crop / bare soil** |  |
|  | May |  |  |  | **Italian rye grass**  Danergo cv.  (35 kg/ha) | **Potato**  Agria cv.  (2750 kg/ha) |  |  | **Oat**  Symphony cv.  (200 kg/ha) |
|  | Jun | **Pumpkin**  Kaori kuri cv.  (18k seeds/ha) |  |  |  |  |  | **White cabbage**  Rivera cv.  (30k plants/ha) |  |
|  | Jul |  |  |  |  |  |  |  |  |
|  | Aug |  |  |  |  |  |  |  |  |
|  | Sep |  | **Green manure**  Rye  (100 kg/ha) |  |  | **Green manure**  Buckwheat  Cv 1  (40 kg/ha) |  |  | **Green manure**  Rye  (100 kg/ha) |
|  | Oct | **Italian rye grass**  Danergo cv.  (35 kg/ha) |  |  |  |  |  |  |  |
|  | Nov |  |  |  |  |  |  | **Winter barley**  Cassiopee cv.  (180 kg/ha) |  |
|  | Dec |  |  |  |  |  |  |  |  |

| **Strip_cultivar** | | | | | | | | | |
| --- | --- | --- | --- | --- | --- | --- | --- | --- | --- |
| **Year** | **Month** | **Strip 1** | **Strip 2** |  | **Strip 3** | **Strip 4** |  | **Strip 5** | **Strip 6** |
| 2019 | Jan | **Leek**  Pluston cv.  (55k plants/ha)  Vitation cv.  (55k plants/ha) | **Italian rye grass**  Danergo cv.  (17.5 kg/ha)  **English rye grass**  Timothy cv.  (17.5 kg/ha) |  | **Winter wheat**  Kelvin cv.  (100 kg/ha)  Julius cv.  (100 kg/ha) | **Italian rye grass**  Danergo cv.  (15 kg/ha)  **English rye grass**  Timothy cv.  (15 kg/ha) |  | **No crop / bare soil** | **Rye**  Ruben cv.  (50 kg/ha)  Diamond cv.  (50 kg/ha) |
|  | Feb |  |  |  |  |  |  |  |  |
|  | Mar |  |  |  |  |  |  |  |  |
|  | Apr |  |  |  |  |  |  |  |  |
|  | May | **Potato**  Agria cv.  (950 kg/ha)  Allouette cv.  (950 kg/ha)  Carolus cv.  (950 kg/ha) |  |  |  | **No crop / bare soil** |  | **Sugar beet**  **(failed)** | **Spring barley**  Irina cv.  (100 kg/ha)  Laureate cv.  (100 kg/ha) |
|  | Jun |  |  |  |  | **White cabbage**  Rivera cv.  (26250 plants/ha)  Christmas Drumhead cv.  (3750 plants/ha) |  | **No crop / bare soil** |  |
|  | Jul |  |  |  |  |  |  |  |  |
|  | Aug |  |  |  | **No crop / bare soil** |  |  |  |  |
|  | Sep |  |  |  |  |  |  |  | **No crop / bare soil** |
|  | Oct | **No crop / bare soil** |  |  | **Green manure**  Buckwheat  Cv 1  (20 kg/ha)  Buckwheat  Cv 2  (20 kg/ha) |  |  |  |  |
|  | Nov | **Winter wheat**  Kelvin cv.  (67 kg/ha)  Julius cv.  (67 kg/ha)  KWS talent cv.  (67 kg/ha) |  |  |  | **No crop / bare soil** |  | **Italian rye grass**  Danergo cv.  (17.5 kg/ha)  **English rye grass**  Timothy cv.  (17.5 kg/ha) | **Green manure**  “Nemacontrol”  (30 kg/ha)  *Composition*:  Horseradish mixture |
|  | Dec |  |  |  |  | **Winter barley**  Cassiopee cv.  (200 kg/ha) |  |  |  |
| 2020 | Jan |  |  |  |  |  |  |  |  |
|  | Feb |  |  |  |  |  |  |  |  |
|  | Mar |  |  |  |  |  |  |  |  |
|  | Apr |  |  |  |  |  |  |  |  |
|  | May |  | **White cabbage**  Rivera cv.  (26250 plants/ha)  Christmas Drumhead cv.  (3750 plants/ha) |  | **Sugar beet**  **(failed)** |  |  |  | **Potato**  Agria cv.  (950 kg/ha)  Allouette cv.  (950 kg/ha)  Carolus cv.  (950 kg/ha) |
|  | Jun |  |  |  | **Pumpkin**  Kaori kuri cv.  (9k seeds/ha)  Orange summer cv.  (9k seeds/ha) |  |  |  |  |
|  | Jul |  |  |  |  |  |  |  |  |
|  | Aug | **No crop / bare soil** |  |  |  | **Winter barley**  Stubble left |  |  |  |
|  | Sep | **Green manure**  Bristle oat  (20 kg/ha)  Summer barley  (20 kg/ha)  Summer wheat  (20 kg/ha) |  |  |  |  |  |  | **No crop / bare soil** |
|  | Oct |  |  |  | **No crop / bare soil** |  |  |  |  |
|  | Nov |  |  |  |  |  |  |  | **Spelt**  **(failed)**  (200 kg/ha) |
|  | Dec |  | **No crop / bare soil** |  |  |  |  |  |  |
| 2021 | Jan |  |  |  |  |  |  |  |  |
|  | Feb |  |  |  |  |  |  |  |  |
|  | Mar |  |  |  |  |  |  |  | **No crop / bare soil** |
|  | Apr |  | **Spring barley**  RGT planet cv.  (100 kg/ha)  KWS Irina cv.  (100 kg/ha) |  |  |  |  | **No crop / bare soil** |  |
|  | May |  |  |  | **Italian rye grass**  Danergo cv.  (15 kg/ha)  **English rye grass**  Timothy cv.  (15 kg/ha) | **Potato**  Agria cv.  (950 kg/ha)  Allouette cv.  (950 kg/ha)  Carolus cv.  (950 kg/ha) |  |  | **Oat**  Symphony cv.  (100 kg/ha)  Armani cv.  (100 kg/ha) |
|  | Jun | **Pumpkin**  Kaori kuri cv.  (9k seeds/ha)  Orange summer cv.  (9k seeds/ha) |  |  |  |  |  | **White cabbage**  Rivera cv.  (26250 plants/ha)  Christmas Drumhead cv.  (3750 plants/ha) |  |
|  | Jul |  |  |  |  |  |  |  |  |
|  | Aug |  |  |  |  |  |  |  |  |
|  | Sep |  | **Green manure**  Rye  (60 kg/ha)  Triticale  (60 kg/ha)  Italian rye grass  (10 kg/ha) |  |  | **Green manure**  Buckwheat  Cv 1  (20 kg/ha)  Buckwheat  Cv 2  (20 kg/ha) |  |  | **Green manure**  Rye  (33 kg/ha)  Triticale  (33 kg/ha)  Italian rye grass  (1.7 kg/ha) |
|  | Oct | **Italian rye grass**  Danergo cv.  (17.5 kg/ha)  **English rye grass**  Timothy cv.  (17.5 kg/ha) |  |  |  |  |  |  |  |
|  | Nov |  |  |  |  |  |  | **Winter barley**  Cassiopee cv.  (90 kg/ha)  SU Jule cv.  (90 kg/ha) |  |
|  | Dec |  |  |  |  |  |  |  |  |

| **Strip_additive** | | | | | | | | | |
| --- | --- | --- | --- | --- | --- | --- | --- | --- | --- |
| **Year** | **Month** | **Strip 1** | **Strip 2** |  | **Strip 3** | **Strip 4** |  | **Strip 5** | **Strip 6** |
| 2019 | Jan | **Leek**  Pluston cv.  (110,000 plants/ha) | **Italian rye grass**  Danergo cv.  (35 kg/ha)  **Red clover**  Salino cv.  (5 kg/ha) |  | **Winter wheat**  Kelvin cv.  (200 kg/ha)  **Faba bean**  Tundra cv.  20k seeds/ha) | **Italian rye grass**  Danergo cv.  (35 kg/ha)  **Red clover**  Salino cv.  (5 kg/ha) |  | **No crop / bare soil** | **Rye**  Ruben cv.  (100 kg/ha)  **Vetch**  (40 kg/ha) |
|  | Feb |  |  |  |  |  |  |  |  |
|  | Mar |  |  |  |  |  |  |  |  |
|  | Apr |  |  |  |  |  |  |  |  |
|  | May | **Potato**  Agria cv.  (2750 kg/ha) |  |  |  | **No crop / bare soil** |  | **Sugar beet**  **(failed)** | **Spring barley**  Irina cv.  (200 kg/ha)  **Summer pea**  Avantgarde cv.  (20k seeds / ha) |
|  | Jun |  |  |  |  | **White cabbage**  Rivera cv.  (30k plants/ha) |  | **No crop / bare soil** |  |
|  | Jul |  |  |  |  |  |  |  |  |
|  | Aug |  |  |  | **No crop / bare soil** |  |  |  |  |
|  | Sep |  |  |  |  |  |  |  | **No crop / bare soil** |
|  | Oct | **No crop / bare soil** |  |  | **Green manure**  Buckwheat  (5 kg/ha)  Borage  (0.78 kg/ha)  Flax  (8.75 kg/ha)  Niger  (1 kg/ha)  Phacelia  (0.75 kg/ha)  Persian clover (10 kg/ha)  Serradella  (17.5 kg/ha)  Sunflower  (2.5 kg/ha) |  |  |  |  |
|  | Nov | **Winter wheat**  Kelvin cv.  (200 kg/ha)  **Faba bean**  Tundra cv.  20k seeds/ha) |  |  |  | **No crop / bare soil** |  | **Italian rye grass**  Danergo cv.  (35 kg/ha)  **Red clover**  Salino cv.  (5 kg/ha) | **Green manure**  “Solarigol TR”  (40 kg/ha)  *Composition*:  Fabaceae  (43%)  Brassicaceae (14%)  Flax  Japanese oat  Niger  Vetch  Persian clover  Fodder radish  Camelina |
|  | Dec |  |  |  |  | **Winter barley**  Cassiopee cv.  (200 kg/ha)  **Winter pea**  Balltrap cv.  (20k seeds / ha) |  |  |  |
| 2020 | Jan |  |  |  |  |  |  |  |  |
|  | Feb |  |  |  |  |  |  |  |  |
|  | Mar |  |  |  |  |  |  |  |  |
|  | Apr |  |  |  |  |  |  |  |  |
|  | May |  | **White cabbage**  Rivera cv.  (30k plants/ha) |  | **Sugar beet**  **(failed)** |  |  |  | **Potato**  Agria cv.  (2750 kg/ha) |
|  | Jun |  |  |  | **Pumpkin**  Kaori kuri cv.  (18k seeds/ha) |  |  |  |  |
|  | Jul |  |  |  |  |  |  |  |  |
|  | Aug | **No crop / bare soil** |  |  |  | **Winter barley**  Stubble left |  |  |  |
|  | Sep | **Green manure**  Buckwheat  (5 kg/ha)  Borage  (0.78 kg/ha)  Flax  (8.75 kg/ha)  Niger  (1 kg/ha)  Phacelia  (0.75 kg/ha)  Persian clover (10 kg/ha)  Serradella  (17.5 kg/ha)  Sunflower  (2.5 kg/ha) |  |  |  |  |  |  | **No crop / bare soil** |
|  | Oct |  |  |  | **No crop / bare soil** |  |  |  |  |
|  | Nov |  |  |  |  |  |  |  | **Spelt**  **(failed)**  (200 kg/ha)  **Faba bean**  Tundra cv.  20k seeds/ha) |
|  | Dec |  | **No crop / bare soil** |  |  |  |  |  |  |
| 2021 | Jan |  |  |  |  |  |  |  |  |
|  | Feb |  |  |  |  |  |  |  |  |
|  | Mar |  |  |  |  |  |  |  | **No crop / bare soil** |
|  | Apr |  | **Spring barley**  RGT Planet cv.  (200 kg/ha)  **Summer pea**  Tiberius cv.  (20k seeds/ha) |  |  |  |  | **No crop / bare soil** |  |
|  | May |  |  |  | **Italian rye grass**  Danergo cv.  (35 kg/ha)  **Red clover**  Salino cv.  (5 kg/ha) | **Potato**  Agria cv.  (2750 kg/ha) |  |  | **Oat**  Symphony cv.  (200 kg/ha)  **Faba bean**  Tundra cv.  20k seeds/ha) |
|  | Jun | **Pumpkin**  Kaori kuri cv.  (18k seeds/ha) |  |  |  |  |  | **White cabbage**  Rivera cv.  (30k plants/ha) |  |
|  | Jul |  |  |  |  |  |  |  |  |
|  | Aug |  |  |  |  |  |  |  |  |
|  | Sep |  | **Green manure**  Rye  (100 kg/ha)  Pea  (70 kg/ha) |  |  | **Green manure**  Buckwheat  (5 kg/ha)  Borage  (0.78 kg/ha)  Flax  (8.75 kg/ha)  Niger  (1 kg/ha)  Phacelia  (0.75 kg/ha)  Persian clover  (10 kg/ha)  Serradella  (17.5 kg/ha)  Sunflower  (2.5 kg/ha) |  |  | **Green manure**  Rye  (30 kg/ha)  Vetch  (30 kg/ha)  Buckwheat  (1.7 kg/ha)  Borage  (0.3 kg/ha)  Flax  (2.9 kg/ha)  Niger  (0.3 kg/ha)  Phacelia  (0.3 kg/ha)  Persian clover  (3.3 kg/ha)  Serradella  (5.8 kg/ha)  Sunflower  (0.8 kg/ha) |
|  | Oct | **Italian rye grass**  Danergo cv.  (35 kg/ha)  **Red clover**  Salino cv.  (5 kg/ha) |  |  |  |  |  |  |  |
|  | Nov |  |  |  |  |  |  | **Winter barley**  Cassiopee cv.  (180 kg/ha)  **Winter pea**  Balltrap cv.  (20k seeds/ha) |  |
|  | Dec |  |  |  |  |  |  |  |  |

| **Strip_diversity** | | | | | | | | | |
| --- | --- | --- | --- | --- | --- | --- | --- | --- | --- |
| **Year** | **Month** | **Strip 1** | **Strip 2** |  | **Strip 3** | **Strip 4** |  | **Strip 5** | **Strip 6** |
| 2019 | Jan | **Leek**  Pluston cv.  (55k plants/ha)  Vitation cv.  (55k plants/ha) | **Italian rye grass**  Danergo cv.  (17.5 kg/ha)  **English rye grass**  Timothy cv.  (17.5 kg/ha)  **Red clover**  Salino cv.  (5 kg/ha) |  | **Winter wheat**  Kelvin cv.  (100 kg/ha)  Julius cv.  (100 kg/ha)  **Faba bean**  Tundra cv.  20k seeds/ha) | **Italian rye grass**  Danergo cv.  (15 kg/ha)  **English rye grass**  Timothy cv.  (15 kg/ha)  **Red clover**  Salino cv.  (5 kg/ha) |  | **No crop / bare soil** | **Rye**  Ruben cv.  (50 kg/ha)  Diamond cv.  (50 kg/ha)  **Vetch**  (40 kg/ha) |
|  | Feb |  |  |  |  |  |  |  |  |
|  | Mar |  |  |  |  |  |  |  |  |
|  | Apr |  |  |  |  |  |  |  |  |
|  | May | **Potato**  Agria cv.  (950 kg/ha)  Allouette cv.  (950 kg/ha)  Carolus cv.  (950 kg/ha) |  |  |  | **No crop / bare soil** |  | **Sugar beet**  **(failed)** | **Spring barley**  Irina cv.  (100 kg/ha)  Laureate cv.  (100 kg/ha)  **Summer pea**  Avantgarde cv.  (20k seeds / ha) |
|  | Jun |  |  |  |  | **White cabbage**  Rivera cv.  (26250 plants/ha)  Christmas Drumhead cv.  (3750 plants/ha) |  | **No crop / bare soil** |  |
|  | Jul |  |  |  |  |  |  |  |  |
|  | Aug |  |  |  | **No crop / bare soil** |  |  |  |  |
|  | Sep |  |  |  |  |  |  |  | **No crop / bare soil** |
|  | Oct | **No crop / bare soil** |  |  | **Green manure**  Buckwheat  (5 kg/ha)  Borage  (0.78 kg/ha)  Flax  (8.75 kg/ha)  Niger  (1 kg/ha)  Phacelia  (0.75 kg/ha)  Persian clover (10 kg/ha)  Serradella  (17.5 kg/ha)  Sunflower  (2.5 kg/ha) |  |  |  |  |
|  | Nov | **Winter wheat**  Kelvin cv.  (67 kg/ha)  Julius cv.  (67 kg/ha)  KWS talent cv.  (67 kg/ha)  **Faba bean**  Tundra cv.  20k seeds/ha) |  |  |  | **No crop / bare soil** |  | **Italian rye grass**  Danergo cv.  (17.5 kg/ha)  **English rye grass**  Timothy cv.  (17.5 kg/ha)  **Red clover**  Salino cv.  (5 kg/ha) | **Green manure**  “Solarigol TR”  (40 kg/ha)  *Composition*:  Fabaceae  (43%)  Brassicaceae (14%)  Flax  Japanese oat  Niger  Vetch  Persian clover  Fodder radish  Camelina |
|  | Dec |  |  |  |  | **Winter barley**  Cassiopee cv.  (200 kg/ha)  **Winter pea**  Balltrap cv.  (20k seeds / ha) |  |  |  |
| 2020 | Jan |  |  |  |  |  |  |  |  |
|  | Feb |  |  |  |  |  |  |  |  |
|  | Mar |  |  |  |  |  |  |  |  |
|  | Apr |  |  |  |  |  |  |  |  |
|  | May |  | **White cabbage**  Rivera cv.  (26250 plants/ha)  Christmas Drumhead cv.  (3750 plants/ha) |  | **Sugar beet**  **(failed)** |  |  |  | **Potato**  Agria cv.  (950 kg/ha)  Allouette cv.  (950 kg/ha)  Carolus cv.  (950 kg/ha) |
|  | Jun |  |  |  | **Pumpkin**  Kaori kuri cv.  (9k seeds/ha)  Orange summer cv.  (9k seeds/ha) |  |  |  |  |
|  | Jul |  |  |  |  |  |  |  |  |
|  | Aug | **No crop / bare soil** |  |  |  | **Winter barley**  Stubble left |  |  |  |
|  | Sep | **Green manure**  Buckwheat  (5 kg/ha)  Borage  (0.78 kg/ha)  Flax  (8.75 kg/ha)  Niger  (1 kg/ha)  Phacelia  (0.75 kg/ha)  Persian clover (10 kg/ha)  Serradella  (17.5 kg/ha)  Sunflower  (2.5 kg/ha) |  |  |  |  |  |  | **No crop / bare soil** |
|  | Oct |  |  |  | **No crop / bare soil** |  |  |  |  |
|  | Nov |  |  |  |  |  |  |  | **Spelt**  **(failed)**  (200 kg/ha)  **Faba bean**  Tundra cv.  20k seeds/ha) |
|  | Dec |  | **No crop / bare soil** |  |  |  |  |  |  |
| 2021 | Jan |  |  |  |  |  |  |  |  |
|  | Feb |  |  |  |  |  |  |  |  |
|  | Mar |  |  |  |  |  |  |  | **No crop / bare soil** |
|  | Apr |  | **Spring barley**  RGT Planet cv.  (100 kg/ha)  KWS Irina cv.  (100 kg/ha)  **Summer pea**  Tiberius cv.  (20k seeds / ha) |  |  |  |  | **No crop / bare soil** |  |
|  | May |  |  |  | **Italian rye grass**  Danergo cv.  (15 kg/ha)  **English rye grass**  Timothy cv.  (15 kg/ha)  **Red clover**  Salino cv.  (5 kg/ha) | **Potato**  Agria cv.  (950 kg/ha)  Allouette cv.  (950 kg/ha)  Carolus cv.  (950 kg/ha) |  |  | **Oat**  Symphony cv.  (100 kg/ha)  Armani cv.  (100 kg/ha)  **Faba bean**  Tundra cv.  20k seeds/ha) |
|  | Jun | **Pumpkin**  Kaori kuri cv.  (9k seeds/ha)  Orange summer cv.  (9k seeds/ha) |  |  |  |  |  | **White cabbage**  Rivera cv.  (26250 plants/ha)  Christmas Drumhead cv.  (3750 plants/ha) |  |
|  | Jul |  |  |  |  |  |  |  |  |
|  | Aug |  |  |  |  |  |  |  |  |
|  | Sep |  | **Green manure**  Rye  (60 kg/ha)  Pea  (70 kg/ha)  Italian rye grass  (10 kg/ha) |  |  | **Green manure**  Buckwheat  (5 kg/ha)  Borage  (0.78 kg/ha)  Flax  (8.75 kg/ha)  Niger  (1 kg/ha)  Phacelia  (0.75 kg/ha)  Persian clover  (10 kg/ha)  Serradella  (17.5 kg/ha)  Sunflower  (2.5 kg/ha) |  |  | **Green manure**  Rye  (20 kg/ha)  Italian rye grass  (1.7 kg/ha)  Vetch  (30 kg/ha)  Buckwheat  (1.7 kg/ha)  Borage  (0.3 kg/ha)  Flax  (2.9 kg/ha)  Niger  (0.3 kg/ha)  Phacelia  (0.3 kg/ha)  Persian clover  (3.3 kg/ha)  Serradella  (5.8 kg/ha)  Sunflower  (0.8 kg/ha) |
|  | Oct | **Italian rye grass**  Danergo cv.  (17.5 kg/ha)  **English rye grass**  Timothy cv.  (17.5 kg/ha)  **Red clover**  Salino cv.  (5 kg/ha) |  |  |  |  |  |  |  |
|  | Nov |  |  |  |  |  |  | **Winter barley**  Cassiopee cv.  (90 kg/ha)  SU Jule cv.  (90 kg/ha)  **Winter pea**  Balltrap cv.  (20k seeds / ha) |  |
|  | Dec |  |  |  |  |  |  |  |  |

**Table S6.** Ground cover and plant mixtures throughout the three years field experiment for Lelystad (Broekemahoeve). Merged cells indicate the period of a certain ground cover. For the monoculture reference each column indicates either cabbage or wheat at the reference field. For the other cropping designs each column depicts the same strips over the three years. “Unknown” means that this information could not be found within the cropping management logs. Crop cultivars and seeding/planting densities are given whenever known.

| **Reference** | | | | | | | |
| --- | --- | --- | --- | --- | --- | --- | --- |
| **Year** | **Month** | **Cabbage Reference 2019** | **Wheat reference Strips 2019** | **Cabbage Reference 2020** | **Wheat Reference Strips 2020** | **Cabbage Reference 2021** | **Wheat reference Strips 2021** |
| 2019 | Jan | **Italian rye grass**  **Red clover** | **Italian rye grass**  **Red clover** | **Italian rye grass**  **Red clover** | **Italian rye grass**  **Red clover** | **Unknown** | **Unknown** |
|  | Feb |  |  |  |  |  |  |
|  | Mar |  | **Spring wheat**  Quintus cv.  (150 kg/ha) |  |  |  |  |
|  | Apr |  |  |  |  | **Potato**  Twinner cv.  (22.500 plants/ha)  Twister cv.  (22.500 plants/ha) | **Potato**  Twinner cv.  (22.500 plants/ha)  Twister cv.  (22.500 plants/ha) |
|  | May | **White cabbage**  Rivera cv.  (50,000 plants/ha) |  |  |  |  |  |
|  | Jun |  |  |  |  |  |  |
|  | Jul |  |  |  |  |  |  |
|  | Aug |  |  |  |  |  |  |
|  | Sep |  | **Green manure**  Buckwheat  cv. 1  (80 kg/ha) |  |  | **Italian rye grass**  **Red clover** | **Italian rye grass**  **Red clover** |
|  | Oct |  |  |  |  |  |  |
|  | Nov |  |  |  |  |  |  |
|  | Dec |  |  |  |  |  |  |
| 2020 | Jan |  |  | **Italian rye grass**  **Red clover** | **Italian rye grass**  **Red clover** | **Italian rye grass**  **Red clover** | **Italian rye grass**  **Red clover** |
|  | Feb |  |  |  |  |  |  |
|  | Mar |  |  |  |  |  |  |
|  | Apr |  |  |  | **Spring wheat**  Lennox cv.  (160 kg/ha) |  |  |
|  | May |  |  |  |  |  |  |
|  | Jun |  |  | **White cabbage**  Rivera cv.  (45,000 plants/ha) |  |  |  |
|  | Jul |  |  |  |  |  |  |
|  | Aug |  |  |  |  |  |  |
|  | Sep |  |  |  |  |  |  |
|  | Oct |  |  |  | **No crop / soil bare** |  |  |
|  | Nov |  |  |  |  |  |  |
|  | Dec |  |  |  |  |  |  |
| 2021 | Jan |  |  |  |  | **Italian rye grass**  **Red clover** | **Italian rye grass**  **Red clover** |
|  | Feb |  |  |  |  |  |  |
|  | Mar |  |  |  |  |  |  |
|  | Apr |  |  |  |  |  | **Spring wheat**  Lennox cv.  (160 kg/ha) |
|  | May |  |  |  |  |  |  |
|  | Jun |  |  |  |  | **White cabbage** Rivera cv.  (45,000 plants/ha) |  |
|  | Jul |  |  |  |  |  |  |
|  | Aug |  |  |  |  |  |  |
|  | Sep |  |  |  |  |  |  |
|  | Oct |  |  |  |  |  | **Green manure**  Buckwheat  cv. 1  (80 kg/ha) |
|  | Nov |  |  |  |  | **No crop / soil bare** |  |
|  | Dec |  |  |  |  |  |  |

| **Strip** | | | | | | | | | | | | |
| --- | --- | --- | --- | --- | --- | --- | --- | --- | --- | --- | --- | --- |
| **Year** | **Month** | **Strip 1** | **Strip 2** |  | **Strip 3** | **Strip 4** |  | **Strip 5** | **Strip 6** |  | **Strip 7** | **Strip 8** |
| 2019 | Jan | **Unknown** | **Unknown** |  | **Unknown** | **No crop / soil bare** |  | **Unknown** | **White clover** |  | **No crop / soil bare** | **Italian rye Grass**  (40 kg/ha) |
|  | Feb |  |  |  |  | **Italian rye Grass**  Meroa cv.  (40 kg/ha) |  |  |  |  |  |  |
|  | Mar |  |  |  | **No crop / soil bare** |  |  |  | **No crop / soil bare** |  | **Spring wheat**  Quintus cv.  (150 kg/ha) |  |
|  | Apr | **Spring barley**  Irina cv.  (150 kg/ha) |  |  | **Potato**  Ditta cv.  (45,000 plants / ha) |  |  | **Onion**  Hylander cv.  (1,000,000 seeds/ha) | **Carrot (J10)**  Nerac cv.  (1,800,000 seeds/ha)  **Buckwheat (J8)**  Kora cv.  (80 kg/ha) |  |  |  |
|  | May |  | **Sugar beet**  Anarosa cv.  (90,000 zaden / ha) |  |  |  |  |  |  |  |  | **White cabbage**  Rivera cv.  (50,000 plants/ha) |
|  | Jun |  |  |  |  |  |  |  |  |  |  |  |
|  | Jul |  |  |  |  |  |  |  |  |  |  |  |
|  | Aug |  |  |  |  |  |  |  |  |  |  |  |
|  | Sep | **Green manure**  Bristle oat  cv. 1  (70 kg/ha) |  |  | **No crop / soil bare** |  |  | **Green manure**  Fodder radish  Radetsky cv.  (40 kg/ha) |  |  | **Green manure**  Buckwheat  cv. 1  (80 kg/ha) |  |
|  | Oct |  |  |  |  |  |  |  |  |  |  |  |
|  | Nov |  | **Italian rye Grass**  Meroa cv.  (40 kg/ha) |  |  |  |  |  | **No crop / soil bare** |  |  |  |
|  | Dec |  |  |  |  |  |  |  |  |  |  | **No crop / soil bare** |
| 2020 | Jan | **Green manure**  Buckwheat  (50 kg/ha) | **Italian rye Grass**  Meroa cv.  (40 kg/ha) |  | **No crop / soil bare** | **Italian rye Grass**  Meroa cv.  (40 kg/ha) |  | **Green manure**  Fodder radish  Radetsky cv.  (40 kg/ha) | **No crop / soil bare** |  | **Green manure**  Fodder radish  (50 kg/ha) | **No crop / soil bare** |
|  | Feb |  |  |  |  |  |  |  |  |  |  |  |
|  | Mar |  |  |  |  |  |  |  |  |  |  |  |
|  | Apr | **Potato**  Ditta cv.  (45,000 plants/ha) |  |  | **Spring wheat**  Lennox cv.  (160 kg/ha) |  |  |  | **Spring barley**  Irina cv.  (120 kg/ha) |  |  |  |
|  | May |  |  |  |  | **No crop / soil bare** |  | **Sugar beet**  Myrtille cv.  (115,000 seeds/ha) |  |  | **Onion**  Hylander cv.  (1,000,000 seeds/ha) | **Carrot**  Nerac cv.  (1,800,000 seeds/ha) |
|  | Jun |  |  |  |  | **White cabbage**  Rivera cv.  (45,000 plants/ha) |  |  |  |  |  |  |
|  | Jul |  |  |  |  |  |  |  |  |  |  |  |
|  | Aug |  |  |  | **No crop / soil bare** |  |  |  |  |  |  |  |
|  | Sep | **No crop / soil bare** |  |  |  |  |  |  | **Green manure**  Winter rye  Dankowskie Diament  (130 kg/ha) |  |  |  |
|  | Oct |  |  |  |  |  |  | **No crop / soil bare** |  |  | **No crop / soil bare** |  |
|  | Nov | **Winter wheat**  Moschus cv.  (175 kg/ha) |  |  |  | **No crop / bare soil** |  | **Italian rye grass**  Meroa cv.  (40 kg/ha) |  |  |  | **No crop / bare soil** |
|  | Dec |  |  |  |  |  |  |  |  |  |  |  |
| 2021 | Jan | **Winter wheat**  Moschus cv.  (175 kg/ha) | **Italian rye Grass**  Meroa cv.  (40 kg/ha) |  | **No crop / soil bare** | **No crop / soil bare** |  | **Italian rye grass**  Meroa cv.  (40 kg/ha) | **Green manure**  Winter rye  (90 kg/ha) |  | **No crop / soil bare** | **No crop / bare soil** |
|  | Feb |  |  |  |  |  |  |  |  |  |  |  |
|  | Mar |  |  |  |  |  |  |  |  |  |  |  |
|  | Apr |  | **No crop / soil bare** |  |  |  |  |  | **No crop / soil bare** |  |  | **Spring barley**  Irina cv.  (120 kg/ha) |
|  | May |  |  |  | **Onion**  Hylander cv.  (1,000,000 seeds/ha) |  |  |  |  |  | **Sugar beet**  Myrtille cv.  (115,000 seeds/ha) |  |
|  | Jun |  | **White cabbage** Rivera cv.  (45,000 plants/ha) |  |  | **Carrot**  Nerac cv.  (1,800,000 seeds/ha) |  |  | **Potato**  Ditta cv.  (45,000 plants/ha) |  |  |  |
|  | Jul |  |  |  |  |  |  |  |  |  |  |  |
|  | Aug | **No crop / soil bare** |  |  |  |  |  |  |  |  |  | **No crop / soil bare** |
|  | Sep | **Green manure**  Buckwheat  cv. 1  (80 kg/ha) |  |  |  |  |  |  | **No crop / soil bare** |  |  | **Green manure**  Winter rye  Dankowskie Diament cv.  (130 kg/ha) |
|  | Oct |  |  |  | **Green manure**  Winter rye  (130 kg/ha) |  |  |  | **Winter wheat**  Moschus cv.  (175 kg/ha) |  |  |  |
|  | Nov |  | **Green manure**  Bristle oat  (90 kg/ha) |  |  | **Winter barley**  (120 kg/ha) |  |  |  |  | **Italian rye grass**  Meroa cv.  (40 kg/ha) |  |
|  | Dec |  |  |  |  |  |  |  |  |  |  |  |

| **Strip_cultivar** | | | | | | | | | | | | |
| --- | --- | --- | --- | --- | --- | --- | --- | --- | --- | --- | --- | --- |
| **Year** | **Month** | **Strip 1** | **Strip 2** |  | **Strip 3** | **Strip 4** |  | **Strip 5** | **Strip 6** |  | **Strip 7** | **Strip 8** |
| 2019 | Jan | **Unknown** | **Unknown** |  | **Unknown** | **No crop / soil bare** |  | **Unknown** | **White clover** |  | **No crop / soil bare** | **Italian rye Grass**  (40 kg/ha) |
|  | Feb |  |  |  |  | **Italian rye Grass**  Meroa cv.  (20 kg/ha)  **Tall fescue**  SWAJ cv.  (20 kg/ha)) |  |  |  |  |  |  |
|  | Mar |  |  |  | **No crop / soil bare** |  |  |  | **No crop / soil bare** |  | **Spring wheat**  Lennox cv.  (80 kg/ha)  Harenda cv.  (80 kg/ha) |  |
|  | Apr | **Spring barley**  Irina cv.  (60 kg/ha)  Planet cv.  (60 kg/ha) |  |  | **Potato**  Ditta cv.  (22,500 plants/ha)  Allouette cv.  (22,500 plants/ha) |  |  | **Onion**  Hylander cv.  (500,000 seeds/ha)  Hytech cv.  (500,000 seeds/ha) | **Carrot (J10)**  Nerac cv.  (900,000 seeds/ha)  Romance cv.  (900,000 seeds/ha  **Buckwheat (J8)**  Kora cv.  (40 kg/ha)  Panda cv.  (40 kg/ha) |  |  |  |
|  | May |  | **Sugar beet**  Anarosa cv.  (45,000 zaden / ha)  ? cv.  (45,000 zaden / ha) |  |  |  |  |  |  |  |  | **White cabbage**  Rivera cv.  (38,750 plants/ha)  Christmas Drumhead cv.  (5,625 plants/ha) |
|  | Jun |  |  |  |  |  |  |  |  |  |  |  |
|  | Jul |  |  |  |  |  |  |  |  |  |  |  |
|  | Aug |  |  |  |  |  |  |  |  |  |  |  |
|  | Sep | **Green manure**  Bristle oat  cv. 1  (35 kg/ha)  Bristle oat  cv. 2  (35 kg/ha) |  |  | **No crop / soil bare** |  |  | **Green manure**  Nemacontrol terralife DSV  (40 kg/ha) |  |  | **Green manure**  Buckwheat  cv. 1  (40 kg/ha)  Buckwheat  cv. 2  (40 kg/ha) |  |
|  | Oct |  |  |  |  |  |  |  |  |  |  |  |
|  | Nov |  | **Italian rye Grass**  Meroa cv.  (20 kg/ha)  **Tall fescue**  SWAJ cv.  (20 kg/ha)) |  |  |  |  |  | **No crop / soil bare** |  |  |  |
|  | Dec |  |  |  |  |  |  |  |  |  |  | **No crop / soil bare** |
| 2020 | Jan | **Green manure**  Buckwheat  (50 kg/ha) | **Italian rye Grass**  Meroa cv.  (30 kg/ha)  **Tall fescue**  SWAJ cv.  (10 kg/ha) |  | **No crop / soil bare** | **Italian rye Grass**  Meroa cv.  (20 kg/ha)  **Tall fescue**  SWAJ cv.  (20 kg/ha) |  | **Green manure**  Nemacontrol terralife DSV  (40 kg/ha) | **No crop / soil bare** |  | **Green manure**  Fodder radish  (50 kg/ha) | **No crop / soil bare** |
|  | Feb |  |  |  |  |  |  |  |  |  |  |  |
|  | Mar |  |  |  |  |  |  |  |  |  |  |  |
|  | Apr | **Potato**  Ditta cv.  (22,500 plants/ha)  Allouette cv.  (22,500 plants/ha) |  |  | **Spring wheat**  Lennox cv.  (80 kg/ha)  Harenda cv.  (80 kg/ha) |  |  |  | **Spring barley**  Irina cv.  (60 kg/ha)  Planet cv.  (60 kg/ha) |  |  |  |
|  | May |  |  |  |  | **No crop / soil bare** |  | **Sugar beet**  Myrtille cv.  (57,500 seeds/ha)  Daphna cv.  (57,500 seeds/ha) |  |  | **Onion**  Hylander cv.  (500,000 seeds/ha)  Hytech cv.  (500,000 seeds/ha) | **Carrot**  Nerac cv.  (900,000 seeds/ha)  Romance cv.  (900,000 seeds/ha) |
|  | Jun |  |  |  |  | **White cabbage**  Rivera cv.  (38,750 plants/ha)  Christmas Drumhead cv.  (5,625 plants/ha) |  |  |  |  |  |  |
|  | Jul |  |  |  |  |  |  |  |  |  |  |  |
|  | Aug |  |  |  | **No crop / soil bare** |  |  |  |  |  |  |  |
|  | Sep | **No crop / soil bare** |  |  |  |  |  |  | **Green manure**  Winter rye  Dankowskie Diament  (65 kg/ha)  Winter barley  Kosmos cv. (60 kg/ha) |  |  |  |
|  | Oct |  |  |  |  |  |  | **No crop / soil bare** |  |  | **No crop / soil bare** |  |
|  | Nov | **Winter wheat**  Moschus cv.  (44 kg/ha)  Extase cv.  (44 kg/ha)  Kelvin cv.  (44 kg/ha)  Talent cv.  (44 kg/ha) |  |  |  | **No crop / bare soil** |  | **Italian rye Grass**  Meroa cv.  (30 kg/ha)  **Tall fescue**  SWAJ cv.  (10 kg/ha) |  |  |  | **No crop / bare soil** |
|  | Dec |  |  |  |  |  |  |  |  |  |  |  |
| 2021 | Jan | **Winter wheat**  Moschus cv.  (44 kg/ha)  Extase cv.  (44 kg/ha)  Kelvin cv.  (44 kg/ha)  Talent cv.  (44 kg/ha) | **Italian rye Grass**  Meroa cv.  (30 kg/ha)  **Tall fescue**  SWAJ cv.  (10 kg/ha) |  | **No crop / soil bare** | **No crop / soil bare** |  | **Italian rye Grass**  Meroa cv.  (30 kg/ha)  **Tall fescue**  SWAJ cv.  (10 kg/ha) | **Green manure**  Winter rye  Dankowskie Diament  (65 kg/ha)  Winter barley  Kosmos cv. (60 kg/ha) |  | **No crop / soil bare** | **No crop / bare soil** |
|  | Feb |  |  |  |  |  |  |  |  |  |  |  |
|  | Mar |  |  |  |  |  |  |  |  |  |  |  |
|  | Apr |  | **No crop / soil bare** |  |  |  |  |  | **No crop / soil bare** |  |  | **Spring barley**  Irina cv.  (60 kg/ha)  Planet cv.  (60 kg/ha) |
|  | May |  |  |  | **Onion**  Hylander cv.  (500,000 seeds/ha)  Hytech cv.  (500,000 seeds/ha) |  |  |  |  |  | **Sugar beet**  Myrtille cv.  (57,500 seeds/ha)  Daphna cv.  (57,500 seeds/ha) |  |
|  | Jun |  | **White cabbage**  Rivera cv.  (38,750 plants/ha)  Christmas Drumhead cv.  (5,625 plants/ha) |  |  | **Carrot**  Nerac cv.  (900,000 seeds/ha)  Romance cv.  (900,000 seeds/ha) |  |  | **Potato**  Ditta cv.  (22,500 plants/ha)  Allouette cv.  (22,500 plants/ha) |  |  |  |
|  | Jul |  |  |  |  |  |  |  |  |  |  |  |
|  | Aug | **No crop / soil bare** |  |  |  |  |  |  |  |  |  | **No crop / soil bare** |
|  | Sep | **Green manure**  Buckwheat  cv. 1  (40 kg/ha)  Buckwheat  cv. 2  (40 kg/ha) |  |  |  |  |  |  | **No crop / soil bare** |  |  | **Green manure**  Winter rye  Dankowskie Diament  (65 kg/ha)  Winter barley  Kosmos cv. (60 kg/ha) |
|  | Oct |  |  |  | **Green manure**  Winter rye  (65 kg/ha)  Winter barley  (60 kg/ha) |  |  |  | **Winter wheat**  Moschus cv.  (44 kg/ha)  Extase cv.  (44 kg/ha)  Kelvin cv.  (44 kg/ha)  Talent cv.  (44 kg/ha) |  |  |  |
|  | Nov |  | **Green manure**  Bristle oat  cv. 1  (45 kg/ha)  cv. 2  (45 kg/ha) |  |  | **No crop / soil bare** |  |  |  |  | **Italian rye Grass**  Meroa cv.  (30 kg/ha)  **Tall fescue**  SWAJ cv.  (10 kg/ha) |  |
|  | Dec |  |  |  |  |  |  |  |  |  |  |  |

| **Strip_additive** | | | | | | | | | | | | |
| --- | --- | --- | --- | --- | --- | --- | --- | --- | --- | --- | --- | --- |
| **Year** | **Month** | **Strip 1** | **Strip 2** |  | **Strip 3** | **Strip 4** |  | **Strip 5** | **Strip 6** |  | **Strip 7** | **Strip 8** |
| 2019 | Jan | **Unknown** | **Unknown** |  | **Unknown** | **No crop / soil bare** |  | **Unknown** | **White clover** |  | **No crop / soil bare** | **Italian rye Grass**  (40 kg/ha) |
|  | Feb |  |  |  |  | **Italian rye Grass**  Meroa cv.  (20 kg/ha)  **Tall fescue**  SWAJ cv.  (8 kg/ha)  **Alfalfa**  Marshal cv.  (8 kg/ha)  **Red clover**  Rozeta cv.  (8 kg/ha)  **White clover**  Nemuniai cv.  (3 kg/ha)  **Chicory**  (1 kg/ha)  **Plantago**  (1 kg/ha)  **Caraway**  (1 kg/ha) |  |  |  |  |  |  |
|  | Mar |  |  |  | **No crop / soil bare** |  |  |  | **No crop / soil bare** |  | **Spring wheat**  Lennox cv.  (90 kg/ha)  **Faba bean**  Tiffany cv.  (90 kg/ha)  **Flower mixture** |  |
|  | Apr | **Spring barley**  Irina cv.  (120 kg/ha)  **Pea**  Balltrapp cv.  (60 kg/ha)  **Flower mixture** |  |  | **Potato**  Ditta cv.  (45,000 plants / ha) |  |  | **Onion**  Hylander cv.  (1,000,000 seeds/ha) | **Carrot (J10)**  Nerac cv.  (1,800,000 seeds/ha)  **Buckwheat (J8)**  Kora cv.  (80 kg/ha)  **Phacelia (J8)**  **White clover (J8)** |  |  |  |
|  | May |  | **Sugar beet**  Anarosa cv.  (90,000 zaden / ha) |  |  |  |  |  |  |  |  | **White cabbage**  Rivera cv.  (50,000 plants/ha) |
|  | Jun |  |  |  |  |  |  |  |  |  |  |  |
|  | Jul |  |  |  |  |  |  |  |  |  |  |  |
|  | Aug |  |  |  |  |  |  |  |  |  |  |  |
|  | Sep | **Green manure**  Solanum terralife DSV  (50 kg/ha) |  |  | **No crop / soil bare** |  |  | **Green manure**  Solarigol TR  (40 kg/ha) |  |  | **Green manure**  Buckwheat  cv. 1  (20 kg/ha)  Phacelia  (10 kg/ha)  Persian clover  (10 kg/ha) |  |
|  | Oct |  |  |  |  |  |  |  |  |  |  |  |
|  | Nov |  | **Grass mixture (see next page)** |  |  |  |  |  | **No crop / soil bare** |  |  |  |
|  | Dec |  |  |  |  |  |  |  |  |  |  | **No crop / soil bare** |
| 2020 | Jan | **Green manure**  Buckwheat  (50 kg/ha) | **Italian rye Grass**  Meroa cv.  (20 kg/ha)  **Tall fescue**  SWAJ cv.  (8 kg/ha)  **Alfalfa**  Marshal cv.  (8 kg/ha)  **Red clover**  Rozeta cv.  (8 kg/ha)  **White clover**  Nemuniai cv.  (3 kg/ha)  **Chicory**  (1 kg/ha)  **Plantago**  (1 kg/ha)  **Caraway**  (1 kg/ha) |  | **No crop / soil bare** | **Grass mixture (see previous page)** |  | **Green manure**  Solarigol TR  (40 kg/ha) | **No crop / soil bare** |  | **Green manure**  Fodder radish  (50 kg/ha) | **No crop / soil bare** |
|  | Feb |  |  |  |  |  |  |  |  |  |  |  |
|  | Mar |  |  |  |  |  |  |  |  |  |  |  |
|  | Apr | **Potato**  Ditta cv.  (45,000 plants/ha) |  |  | **Spring wheat**  Lennox cv.  (90 kg/ha)  **Faba bean**  Tiffany cv.  (90 kg/ha)  **Flower mixture** |  |  |  | **Spring barley**  Irina cv.  (120 kg/ha)  **Pea**  Balltrapp cv.  (60 kg/ha)  **Flower mixture** |  |  |  |
|  | May |  |  |  |  | **No crop / soil bare** |  | **Sugar beet**  Myrtille cv.  (115,000 seeds/ha) |  |  | **Onion**  Hylander cv.  (1,000,000 seeds/ha) | **Carrot**  Nerac cv.  (1,800,000 seeds/ha) |
|  | Jun |  |  |  |  | **White cabbage**  Rivera cv.  (45,000 plants/ha) |  |  |  |  |  |  |
|  | Jul |  |  |  |  |  |  |  |  |  |  |  |
|  | Aug |  |  |  | **No crop / soil bare** |  |  |  |  |  |  |  |
|  | Sep | **No crop / soil bare** |  |  |  |  |  |  | **Green manure**  Winter rye  Dankowskie Diament  (65 kg/ha)  Vetch  Hungvillosa cv.  (40 kg/ha)  White Mustard  Maryna cv.  (5 kg/ha) |  |  |  |
|  | Oct |  |  |  |  |  |  | **No crop / soil bare** |  |  | **No crop / soil bare** |  |
|  | Nov | **Winter wheat**  Moschus cv.  (175 kg/ha)  **Faba bean**  Tundra cv.  (40 kg/ha)  **Flower mixture** |  |  |  | **No crop / bare soil** |  | **Grass mixture (see next page)** |  |  |  | **No crop / bare soil** |
|  | Dec |  |  |  |  |  |  |  |  |  |  |  |
| 2021 | Jan | **Winter wheat**  Moschus cv.  (175 kg/ha)  **Faba bean**  Tundra cv.  (40 kg/ha)  **Flower mixture** | **Grass mixture (see previous page)** |  | **No crop / soil bare** | **No crop / soil bare** |  | **Italian rye Grass**  Meroa cv.  (20 kg/ha)  **Tall fescue**  SWAJ cv.  (8 kg/ha)  **Alfalfa**  Marshal cv.  (8 kg/ha)  **Red clover**  Rozeta cv.  (8 kg/ha)  **White clover**  Nemuniai cv.  (3 kg/ha)  **Chicory**  (1 kg/ha)  **Plantago**  (1 kg/ha)  **Caraway**  (1 kg/ha) | **Green manure**  Winter rye  (90 kg/ha) |  | **No crop / soil bare** | **No crop / bare soil** |
|  | Feb |  |  |  |  |  |  |  |  |  |  |  |
|  | Mar |  |  |  |  |  |  |  |  |  |  |  |
|  | Apr |  | **No crop / soil bare** |  |  |  |  |  | **No crop / soil bare** |  |  | **Spring barley**  Irina cv.  (120 kg/ha)  **Pea**  Balltrapp cv.  (60 kg/ha)  **Flower mixture** |
|  | May |  |  |  | **Onion**  Hylander cv.  (1,000,000 seeds/ha) |  |  |  |  |  | **Sugar beet**  Myrtille cv.  (115,000 seeds/ha) |  |
|  | Jun |  | **White cabbage** Rivera cv.  (45,000 plants/ha) |  |  | **Carrot**  Nerac cv.  (1,800,000 seeds/ha) |  |  | **Potato**  Ditta cv.  (45,000 plants/ha) |  |  |  |
|  | Jul |  |  |  |  |  |  |  |  |  |  |  |
|  | Aug | **No crop / soil bare** |  |  |  |  |  |  |  |  |  | **No crop / soil bare** |
|  | Sep | **Green manure**  Buckwheat  cv. 1  (20 kg/ha)  Phacelia  (10 kg/ha)  Persian clover  (10 kg/ha) |  |  |  |  |  |  | **No crop / soil bare** |  |  | **Green manure**  Winter rye  Dankowskie Diament cv.  (65 kg/ha)  Vetch  Hungvillosa cv.  (40 kg/ha)  White Mustard  Maryna cv.  (5 kg/ha) |
|  | Oct |  |  |  | **Green manure**  Winter rye  Dankowskie Diament cv.  (65 kg/ha)  Vetch  Hungvillosa cv.  (40 kg/ha)  White Mustard  Maryna cv.  (5 kg/ha) |  |  |  | **Winter wheat**  Moschus cv.  (175 kg/ha)  **Faba bean**  Tundra cv.  (40 kg/ha)  **Flower mixture** |  |  |  |
|  | Nov |  | **Green manure**  Bristle oat  (70 kg/ha)  Vetch  Hungvillosa cv.  (90 kg/ha) |  |  | **Winter barley**  (120 kg/ha)  **Pea**  Balltrapp cv.  (33 kg/ha)  **Flower mixture** |  |  |  |  | **Grass mixture**  **(see column strip 5)** |  |
|  | Dec |  |  |  |  |  |  |  |  |  |  |  |

**Table S7.** Number of replications per location and year. Per cropping design we indicate the number of fields, the number of strips (of 4 plant rows) and the total number of samples (individual cabbages).

| Location | Year | Rounds | Fields | Strips | Total samples |
| --- | --- | --- | --- | --- | --- |
| Lelystad | 2019 | 3 | Monoculture: 1  Strip: 2  Strip_cultivar: 2  Strip_additive: 2 | Monoculture: 4  Strip: 4  Strip_cultivar: 4  Strip_additive: 4 | Monoculture: 31  Strip: 31  Strip_cultivar: 29  Strip_additive: 28 |
|  | 2020 | 4 | Monoculture: 1  Strip: 3  Strip_cultivar: 2  Strip_additive: 2 | Monoculture: 4  Strip: 7  Strip_cultivar: 4  Strip_additive: 4 | Monoculture: 22  Strip: 49  Strip_cultivar: 26  Strip_additive: 32 |
|  | 2021 |  |  |  | Monoculture: 32  Strip: 54  Strip_cultivar: 32  Strip_additive: 31 |
| Wageningen | 2019 | 4 | Monoculture: 1  Strip: 4  Strip_cultivar: 3  Strip_additive: 3  Strip_diversity: 3 | Monoculture: 6  Strip: 8  Strip_cultivar: 6  Strip_additive: 6  Strip_diversity: 3 | Monoculture: 44  Strip: 73  Strip_cultivar: 41  Strip_additive: 41  Strip_diversity: 43 |
|  | 2020 |  |  |  | Monoculture: 47  Strip: 72  Strip_cultivar: 46  Strip_additive: 46  Strip_diversity: 40 |
|  | 2021 |  | Monoculture: 1  Strip: 4  Strip_cultivar: 3  Strip_additive: 3  Strip_diversity: 3 | Monoculture: 6  Strip: 10  Strip_cultivar: 6  Strip_additive: 6  Strip_diversity: 3 | Monoculture: 48  Strip: 76  Strip_cultivar: 47  Strip_additive: 44  Strip_diversity: 44 |

**Table S8.** Effect of cropping design on *cabbage head weight*. Results from ANOVA using the final models when using all data, only the data from the reference field at both locations separately and of the other fields at both locations separately. Bold numbers indicate significant effects (α = 0.05).

| Dataset | Random effects | Predictor | Df | Chi sq | P |
| --- | --- | --- | --- | --- | --- |
| All data | Field | Cropping design | 4 | 14.0 | **0.007** |
|  | Field : Strip_ID | Year | 2 | 234.4 | **<0.001** |
|  |  | Location | 1 | 15.0 | **<0.001** |
|  |  | Cropping design : Year | 8 | 26.2 | **<0.001** |
|  |  | Year : Location | 2 | 47.5 | **<0.001** |
| Wageningen | Strip_ID | Cropping design | 1 | 1.06 | 0.302 |
| Monoculture field |  | Placement | 1 | 0.68 | 0.410 |
|  |  | Year | 2 | 142.2 | **<0.001** |
| Wageningen | Strip_ID | Cropping design | 3 | 21.2 | **<0.001** |
| Excl. Monoculture |  | Placement | 1 | 11.1 | **<0.001** |
|  |  | Year | 2 | 69.4 | **<0.001** |
|  |  | Field | 2 | 41.1 | **<0.001** |
|  |  | Cropping design : Year | 6 | 28.0 | **<0.001** |
|  |  | Year : Field | 4 | 8.72 | 0.068 |
| Lelystad | Strip_ID | Cropping design | 1 | 2.42 | 0.120 |
| Monoculture field |  | Placement | 1 | 0.01 | 0.939 |
|  |  | Year | 2 | 6.60 | **0.037** |
| Lelystad | Strip_ID | Cropping design | 2 | 17.9 | **<0.001** |
| Excl. Monoculture |  | Placement | 1 | 3.86 | **0.050** |
|  |  | Year | 2 | 313.6 | **<0.001** |
|  |  | Field | 1 | 2.64 | 0.104 |
|  |  | Cropping design : Year | 4 | 26.7 | **<0.001** |

**Table S9.** Effect of cropping design on *proportion damaged head weight*. Results from ANOVA using the final models when using data from both locations combined and both locations separately. Bold numbers indicate significant effects (α = 0.05).

| Dataset | Random effects | Predictor | Df | Chi sq | P |
| --- | --- | --- | --- | --- | --- |
| Both locations |  | Cropping design | 3 | 5.24 | 0.155 |
| (Excl. Strip_diversity) |  | Year | 2 | 119.8 | **<0.001** |
|  |  | Location | 1 | 7.43 | **0.006** |
|  |  | Cropping design : Year | 6 | 12.4 | 0.053 |
|  |  | Cropping design : Location | 3 | 2.91 | 0.406 |
|  |  | Year : Location | 2 | 3.95 | 0.139 |
|  |  | Cropping design : Year : Location | 6 | 21.7 | **0.001** |
| Wageningen |  | Cropping design | 4 | 7.40 | 0.116 |
|  |  | Placement | 1 | 4.05 | **0.044** |
|  |  | Year | 2 | 60.6 | **<0.001** |
|  |  | Field | 3 | 5.35 | 0.148 |
|  |  | Cropping design : Year | 8 | 8.84 | 0.356 |
| Lelystad |  | Cropping design | 3 | 1.30 | 0.730 |
|  |  | Placement | 1 | 0.03 | 0.864 |
|  |  | Year | 2 | 79.4 | **<0.001** |
|  |  | Field | 2 | 0.79 | 0.672 |
|  |  | Cropping design : Year | 6 | 29.6 | **<0.001** |

**Table S10.** Effect of cropping design and cabbage cultivar on *log transformed early herbivore abundance*. Results from ANOVA using the final models when using data from both locations separately. Bold numbers indicate significant effects (α = 0.05).

| Dataset | Random effects | Predictor | Df | Chi sq | P |
| --- | --- | --- | --- | --- | --- |
| Wageningen |  | Cropping design | 4 | 8.19 | 0.085 |
|  |  | Placement | 1 | 10.2 | **0.001** |
|  |  | Year | 2 | 213.2 | **<0.001** |
|  |  | Field | 3 | 14.0 | **0.003** |
|  |  | Cropping design : Year | 8 | 20.2 | **0.009** |
| Lelystad |  | Cropping design | 3 | 46.9 | **<0.001** |
|  |  | Placement | 1 | 11.4 | **<0.001** |
|  |  | Year | 2 | 848.6 | **<0.001** |
|  |  | Field | 2 | 33.4 | **<0.001** |
|  |  | Cropping design : Year | 6 | 35.2 | **<0.001** |
| Wageningen |  | Cultivar | 1 | 76.7 | **<0.001** |
| Strip_cultivar |  | Year | 2 | 36.4 | **<0.001** |
|  |  | Field | 2 | 10.6 | **0.005** |
|  |  | Cultivar : Year | 2 | 36.0 | **<0.001** |
| Lelystad |  | Cultivar | 1 | 0.16 | 0.692 |
| Strip_cultivar |  | Year | 2 | 223.4 | **<0.001** |
|  |  | Field | 1 | 1.56 | 0.211 |
|  |  | Cultivar : Year | 2 | 1.42 | 0.491 |

**Table S11.** Effect of cropping design and cabbage cultivar on *log transformed late herbivore abundance*. Results from ANOVA using the final models when using data from both locations separately. Bold numbers indicate significant effects (α = 0.05).

| Dataset | Random effects | Predictor | Df | Chi sq | P |
| --- | --- | --- | --- | --- | --- |
| Wageningen |  | Cropping design | 4 | 15.8 | **0.003** |
|  |  | Placement | 1 | 8.25 | **0.004** |
|  |  | Year | 2 | 178.3 | **<0.001** |
|  |  | Field | 3 | 13.8 | **0.003** |
|  |  | Cropping design : Placement | 4 | 1.15 | 0.886 |
|  |  | Cropping design : Year | 8 | 32.8 | **<0.001** |
| Lelystad |  | Cropping design | 3 | 36.7 | **<0.001** |
|  |  | Placement | 1 | 2.90 | 0.088 |
|  |  | Year | 2 | 1136.1 | **<0.001** |
|  |  | Field | 2 | 14.4 | **<0.001** |
|  |  | Cropping design : Year | 6 | 21.3 | **0.002** |
| Wageningen |  | Cultivar | 1 | 77.3 | **<0.001** |
| Strip_cultivar |  | Year | 2 | 5.27 | 0.072 |
|  |  | Field | 2 | 9.14 | **0.010** |
|  |  | Cultivar : Year | 2 | 21.3 | **<0.001** |
| Lelystad |  | Cultivar | 1 | 0.65 | 0.422 |
| Strip_cultivar |  | Year | 2 | 276.5 | **<0.001** |
|  |  | Field | 1 | 6.39 | **0.011** |
|  |  | Cultivar : Year | 2 | 17.3 | **<0.001** |

**Table S12.** Effect of cropping design and cabbage cultivar on *early herbivore richness*. Results from ANOVA using the final models when using data from both locations separately. Bold numbers indicate significant effects (α = 0.05).

| Dataset | Random effects | Predictor | Df | Chi sq | P |
| --- | --- | --- | --- | --- | --- |
| Wageningen |  | Cropping design | 4 | 26.7 | **<0.001** |
|  |  | Placement | 1 | 3.94 | **0.047** |
|  |  | Year | 2 | 109.2 | **<0.001** |
|  |  | Field | 3 | 26.7 | **<0.001** |
|  |  | Cropping design : Year | 8 | 12.3 | 0.136 |
| Lelystad |  | Cropping design | 3 | 15.4 | **0.001** |
|  |  | Placement | 1 | 6.75 | **0.009** |
|  |  | Year | 2 | 45.7 | **<0.001** |
|  |  | Field | 2 | 54.2 | **<0.001** |
|  |  | Cropping design : Year | 6 | 31.3 | **<0.001** |
| Wageningen |  | Cultivar | 1 | 26.7 | **<0.001** |
| Strip_cultivar |  | Year | 2 | 64.9 | **<0.001** |
|  |  | Field | 2 | 17.5 | **<0.001** |
|  |  | Cultivar : Year | 2 | 21.5 | **<0.001** |
| Lelystad |  | Cultivar | 1 | 1.20 | 0.273 |
| Strip_cultivar |  | Year | 2 | 25.9 | **<0.001** |
|  |  | Field | 1 | 0.02 | 0.897 |
|  |  | Cultivar : Year | 2 | 3.02 | 0.221 |

**Table S13.** Effect of cropping design and cabbage cultivar on *late herbivore richness*. Results from ANOVA using the final models when using data from both locations separately. Bold numbers indicate significant effects (α = 0.05).

| Dataset | Random effects | Predictor | Df | Chi sq | P |
| --- | --- | --- | --- | --- | --- |
| Wageningen |  | Cropping design | 4 | 53.5 | **<0.001** |
|  |  | Placement | 1 | 0.83 | 0.363 |
|  |  | Year | 2 | 71.8 | **<0.001** |
|  |  | Field | 3 | 26.0 | **<0.001** |
|  |  | Cropping design : Year | 8 | 19.0 | **0.015** |
| Lelystad |  | Cropping design | 3 | 12.2 | **0.007** |
|  |  | Placement | 1 | 0.03 | 0.872 |
|  |  | Year | 2 | 301.3 | **<0.001** |
|  |  | Field | 2 | 66.6 | **<0.001** |
|  |  | Cropping design : Year | 6 | 36.7 | **<0.001** |
| Wageningen |  | Cultivar | 1 | 66.5 | **<0.001** |
| Strip_cultivar |  | Year | 2 | 6.08 | **0.048** |
|  |  | Field | 2 | 22.6 | **<0.001** |
|  |  | Cultivar : Year | 2 | 31.9 | **<0.001** |
| Lelystad |  | Cultivar | 1 | 1.63 | 0.201 |
| Strip_cultivar |  | Year | 2 | 111.9 | **<0.001** |
|  |  | Field | 1 | 28.6 | **<0.001** |
|  |  | Cultivar : Year | 2 | 4.85 | 0.089 |

**Table S14.** Effect of cropping design and cabbage cultivar on *early chewer abundance*. Results from ANOVA using the final models when using data from both locations separately. Bold numbers indicate significant effects (α = 0.05).

| Dataset | Random effects | Predictor | Df | Chi sq | P |
| --- | --- | --- | --- | --- | --- |
| Wageningen |  | Cropping design | 4 | 3.75 | 0.441 |
|  |  | Placement | 1 | 0.70 | 0.401 |
|  |  | Year | 2 | 350.3 | **<0.001** |
|  |  | Field | 3 | 0.87 | 0.832 |
|  |  | Cropping design : Year | 8 | 20.1 | **0.010** |
| Lelystad |  | Cropping design | 3 | 92.1 | **<0.001** |
|  |  | Placement | 1 | 6.66 | **0.010** |
|  |  | Year | 2 | 1650.3 | **<0.001** |
|  |  | Field | 2 | 10.6 | **0.005** |
|  |  | Cropping design : Year | 6 | 24.2 | **<0.001** |
| Wageningen |  | Cultivar | 1 | 0.09 | 0.763 |
| Strip_cultivar |  | Year | 2 | 59.4 | **<0.001** |
|  |  | Field | 2 | 4.17 | 0.124 |
|  |  | Cultivar : Year | 2 | 19.2 | **<0.001** |
| Lelystad |  | Cultivar | 1 | 1.08 | 0.299 |
| Strip_cultivar |  | Year | 2 | 533.2 | **<0.001** |
|  |  | Field | 1 | 1.87 | 0.172 |
|  |  | Cultivar : Year | 2 | 0.75 | 0.688 |

**Table S15.** Effect of cropping design and cabbage cultivar on *late chewer abundance*. Results from ANOVA using the final models when using data from both locations separately. Bold numbers indicate significant effects (α = 0.05).

| Dataset | Random effects | Predictor | Df | Chi sq | P |
| --- | --- | --- | --- | --- | --- |
| Wageningen |  | Cropping design | 4 | 17.1 | **0.002** |
|  |  | Placement | 1 | 16.0 | **<0.001** |
|  |  | Year | 2 | 581.0 | **<0.001** |
|  |  | Field | 3 | 22.7 | **<0.001** |
|  |  | Cropping design : Year | 8 | 30.4 | **<0.001** |
| Lelystad |  | Cropping design | 3 | 5.55 | 0.136 |
|  |  | Placement | 1 | 4.59 | **0.032** |
|  |  | Year | 2 | 90.8 | **<0.001** |
|  |  | Field | 2 | 7.80 | **0.020** |
|  |  | Cropping design : Year | 6 | 36.5 | **<0.001** |
| Wageningen |  | Cultivar | 1 | 1.25 | 0.264 |
| Strip_cultivar |  | Year | 2 | 116.7 | **<0.001** |
|  |  | Field | 2 | 8.00 | **0.018** |
|  |  | Cultivar : Year | 2 | 40.1 | **<0.001** |
| Lelystad |  | Cultivar | 1 | 0.01 | 0.939 |
| Strip_cultivar |  | Year | 2 | 12.6 | **0.002** |
|  |  | Field | 1 | 1.77 | 0.184 |
|  |  | Cultivar : Year | 2 | 11.8 | **0.003** |

**Table S16.** Effect of cropping design and cabbage cultivar on *early sucker abundance*. Results from ANOVA using the final models when using data from both locations separately. Bold numbers indicate significant effects (α = 0.05).

| Dataset | Random effects | Predictor | Df | Chi sq | P |
| --- | --- | --- | --- | --- | --- |
| Wageningen |  | Cropping design | 4 | 4.55 | 0.336 |
|  |  | Placement | 1 | 4.58 | **0.032** |
|  |  | Year | 2 | 145.1 | **<0.001** |
|  |  | Field | 3 | 1.09 | 0.779 |
|  |  | Cropping design : Year | 8 | 36.4 | **<0.001** |
| Lelystad |  | Cropping design | 3 | 2.55 | 0.466 |
|  |  | Placement | 1 | 5.14 | **0.023** |
|  |  | Year | 2 | 201.6 | **<0.001** |
|  |  | Field | 2 | 12.0 | **0.002** |
|  |  | Cropping design : Year | 6 | 17.1 | **0.009** |
| Wageningen |  | Cultivar | 1 | 77.5 | **<0.001** |
| Strip_cultivar |  | Year | 2 | 49.8 | **<0.001** |
|  |  | Field | 2 | 0.14 | 0.931 |
|  |  | Cultivar : Year | 2 | 39.7 | **<0.001** |
| Lelystad |  | Cultivar | 1 | 1.68 | 0.195 |
| Strip_cultivar |  | Year | 2 | 87.0 | **<0.001** |
|  |  | Field | 1 | 5.38 | **0.020** |
|  |  | Cultivar : Year | 2 | 4.59 | 0.101 |

**Table S17.** Effect of cropping design and cabbage cultivar on *late sucker abundance*. Results from ANOVA using the final models when using data from both locations separately. Bold numbers indicate significant effects (α = 0.05).

| Dataset | Random effects | Predictor | Df | Chi sq | P |
| --- | --- | --- | --- | --- | --- |
| Wageningen |  | Cropping design | 4 | 12.5 | **0.014** |
|  |  | Placement | 1 | 7.72 | **0.005** |
|  |  | Year | 2 | 49.3 | **<0.001** |
|  |  | Field | 3 | 2.01 | 0.571 |
|  |  | Cropping design : Year | 8 | 18.3 | **0.019** |
| Lelystad |  | Cropping design | 3 | 1.23 | 0.745 |
|  |  | Placement | 1 | 0.88 | 0.348 |
|  |  | Year | 2 | 5.77 | 0.056 |
|  |  | Field | 2 | 7.53 | **0.023** |
|  |  | Cropping design : Year | 6 | 21.9 | **0.001** |
| Wageningen |  | Cultivar | 1 | 113.5 | **<0.001** |
| Strip_cultivar |  | Year | 2 | 15.9 | **<0.001** |
|  |  | Field | 2 | 0.96 | 0.618 |
|  |  | Cultivar : Year | 2 | 9.97 | **0.007** |
| Lelystad |  | Cultivar | 1 | 19.5 | **<0.001** |
| Strip_cultivar |  | Year | 2 | 2.13 | 0.345 |
|  |  | Field | 1 | 6.97 | **0.008** |
|  |  | Cultivar : Year | 2 | 1.82 | 0.402 |

**Table S18.** Effect of cropping design and cabbage cultivar on *early thrips abundance*. Results from ANOVA using the final models when using data from both locations separately. Bold numbers indicate significant effects (α = 0.05).

| Dataset | Random effects | Predictor | Df | Chi sq | P |
| --- | --- | --- | --- | --- | --- |
| Wageningen |  | Cropping design | 4 | 9.90 | **0.042** |
|  |  | Placement | 1 | 0.09 | 0.764 |
|  |  | Year | 2 | 62.2 | **<0.001** |
|  |  | Field | 3 | 1.76 | 0.623 |
|  |  | Cropping design : Year | 8 | 15.0 | 0.058 |
| Lelystad |  | Cropping design | 3 | 6.85 | 0.077 |
|  |  | Placement | 1 | 0.83 | 0.363 |
|  |  | Year | 2 | 32.6 | **<0.001** |
|  |  | Field | 2 | 6.68 | **0.035** |
|  |  | Cropping design : Year | 6 | 16.8 | **0.010** |
| Wageningen |  | Cultivar | 1 | 0.08 | 0.778 |
| Strip_cultivar |  | Year | 2 | 13.0 | **0.001** |
|  |  | Field | 2 | 1.97 | 0.373 |
|  |  | Cultivar : Year | 2 | 2.27 | 0.321 |
| Lelystad |  | Cultivar | 1 | 3.81 | 0.051 |
| Strip_cultivar |  | Year | 2 | 23.0 | **<0.001** |
|  |  | Field | 1 | 1.25 | 0.264 |
|  |  | Cultivar : Year | 2 | 0.18 | 0.916 |

**Table S19.** Effect of cropping design and cabbage cultivar on *late thrips abundance*. Results from ANOVA using the final models when using data from both locations separately. Bold numbers indicate significant effects (α = 0.05).

| Dataset | Random effects | Predictor | Df | Chi sq | P |
| --- | --- | --- | --- | --- | --- |
| Wageningen |  | Cropping design | 4 | 14.9 | **0.005** |
|  |  | Placement | 1 | 0.64 | 0.424 |
|  |  | Year | 2 | 5.91 | 0.052 |
|  |  | Field | 3 | 2.08 | 0.555 |
|  |  | Cropping design : Year | 8 | 18.0 | **0.021** |
| Lelystad |  | Cropping design | 3 | 33.2 | **<0.001** |
|  |  | Placement | 1 | 31.9 | **<0.001** |
|  |  | Year | 2 | 1186.4 | **<0.001** |
|  |  | Field | 2 | 15.5 | **<0.001** |
|  |  | Cropping design : Year | 6 | 39.1 | **<0.001** |
| Wageningen |  | Cultivar | 1 | 2.18 | 0.139 |
| Strip_cultivar |  | Year | 2 | 7.35 | **0.025** |
|  |  | Field | 2 | 0.84 | 0.657 |
|  |  | Cultivar : Year | 2 | 0.00 | 1.000 |
| Lelystad |  | Cultivar | 1 | 2.61 | 0.106 |
| Strip_cultivar |  | Year | 2 | 325.5 | **<0.001** |
|  |  | Field | 1 | 0.25 | 0.616 |
|  |  | Cultivar : Year | 2 | 0.08 | 0.960 |

**Table S20.** Effect of cropping design and cabbage cultivar on *early flea beetle abundance*. Results from ANOVA using the final models when using data from both locations separately. Bold numbers indicate significant effects (α = 0.05).

| Dataset | Random effects | Predictor | Df | Chi sq | P |
| --- | --- | --- | --- | --- | --- |
| Wageningen |  | Cropping design | 4 | 38.2 | **<0.001** |
|  |  | Placement | 1 | 15.9 | **<0.001** |
|  |  | Year | 2 | 78.0 | **<0.001** |
|  |  | Field | 3 | 185.2 | **<0.001** |
|  |  | Cropping design : Year | 8 | 24.8 | **0.002** |
| Lelystad |  | Cropping design | 3 | 47.8 | **<0.001** |
|  |  | Placement | 1 | 8.33 | **0.004** |
|  |  | Year | 2 | 166.6 | **<0.001** |
|  |  | Field | 2 | 55.7 | **<0.001** |
|  |  | Cropping design : Year | 6 | 24.6 | **<0.001** |
| Wageningen |  | Cultivar | 1 | 11.3 | **<0.001** |
| Strip_cultivar |  | Year | 2 | 48.9 | **<0.001** |
|  |  | Field | 2 | 82.6 | **<0.001** |
|  |  | Cultivar : Year | 2 | 7.63 | **0.022** |
| Lelystad |  | Cultivar | 1 | 0.07 | 0.793 |
| Strip_cultivar |  | Year | 2 | 39.8 | **<0.001** |
|  |  | Field | 1 | 2.32 | 0.128 |
|  |  | Cultivar : Year | 2 | 5.00 | 0.082 |

**Table S21.** Effect of cropping design and cabbage cultivar on *late flea beetle abundance*. Results from ANOVA using the final models when using data from both locations separately. Bold numbers indicate significant effects (α = 0.05).

| Dataset | Random effects | Predictor | Df | Chi sq | P |
| --- | --- | --- | --- | --- | --- |
| Wageningen |  | Cropping design | 4 | 50.7 | **<0.001** |
|  |  | Placement | 1 | 7.95 | **0.005** |
|  |  | Year | 2 | 129.0 | **<0.001** |
|  |  | Field | 3 | 97.8 | **<0.001** |
|  |  | Cropping design : Year | 8 | 35.8 | **<0.001** |
| Lelystad |  | Cropping design | 3 | 54.3 | **<0.001** |
|  |  | Placement | 1 | 1.39 | 0.238 |
|  |  | Year | 2 | 546.8 | **<0.001** |
|  |  | Field | 2 | 96.5 | **<0.001** |
|  |  | Cropping design : Year | 6 | 11.0 | 0.089 |
| Wageningen |  | Cultivar | 1 | 35.5 | **<0.001** |
| Strip_cultivar |  | Year | 2 | 81.9 | **<0.001** |
|  |  | Field | 2 | 36.2 | **<0.001** |
|  |  | Cultivar : Year | 2 | 11.9 | **0.003** |
| Lelystad |  | Cultivar | 1 | 17.2 | **<0.001** |
| Strip_cultivar |  | Year | 2 | 219.2 | **<0.001** |
|  |  | Field | 1 | 50.1 | **<0.001** |
|  |  | Cultivar : Year | 2 | 0.80 | 0.672 |

**Table S22.** Effect of early and late season plant size on *fresh weight of the cabbage head*, per cropping design. Both direct and indirect effects of early and late plant size variables are shown. Significant direct effects are indicated in bold (α = 0.05). For indirect or total effects it was not possible to assess significance.

|  | Early plant diameter | | |  | Early number of leaves | | |  | Fresh weight wrapper leaves (log) | | |
| --- | --- | --- | --- | --- | --- | --- | --- | --- | --- | --- | --- |
| Cropping system | Direct | Indirect | Total |  | Direct | Indirect | Total |  | Direct | Indirect | Total |
| All | **0.18** | 0.29 | 0.47 |  | 0.03 | -0.03 | 0.00 |  | **0.57** | 0.00 | 0.57 |
| Monoculture | 0.06 | 0.50 | 0.56 |  | **0.16** | -0.06 | 0.10 |  | **0.54** | 0.00 | 0.54 |
| Strip | **0.43** | 0.23 | 0.66 |  | 0.08 | 0.02 | 0.10 |  | **0.39** | 0.00 | 0.39 |
| Strip_cultivar | **0.17** | 0.14 | 0.31 |  | -0.09 | -0.05 | -0.14 |  | **0.60** | 0.00 | 0.60 |
| Strip_additive | **0.33** | 0.28 | 0.61 |  | 0.06 | 0.01 | 0.07 |  | **0.54** | 0.00 | 0.54 |
| Strip_diversity | 0.17 | 0.38 | 0.55 |  | 0.14 | 0.12 | 0.26 |  | **0.68** | 0.00 | 0.68 |

**Table S23.** Effects of early and late season plant size, herbivore abundance and richness, and cabbage head damage on *fresh weight of the cabbage head*, per cropping design. Both direct and indirect effects of early and late plant size variables are shown. Significant direct effects are indicated in bold (α = 0.05). For indirect or total effects it was not possible to assess significance.

|  | Early plant diameter | | |  | Early number of leaves | | |  | Fresh weight wrapper leaves (log) | | |  | Proportion damaged head weight | | |
| --- | --- | --- | --- | --- | --- | --- | --- | --- | --- | --- | --- | --- | --- | --- | --- |
| Cropping system | Direct | Indirect | Total |  | Direct | Indirect | Total |  | Direct | Indirect | Total |  | Direct | Indirect | Total |
| All | **0.21** | 0.27 | 0.48 |  | 0.05 | 0.02 | 0.07 |  | **0.56** | 0.01 | 0.57 |  | **0.07** | 0.00 | 0.07 |
| Monoculture | 0.06 | 0.50 | 0.56 |  | **0.16** | -0.05 | 0.11 |  | **0.54** | 0.00 | 0.54 |  | 0.00 | 0.00 | 0.00 |
| Strip | **0.46** | 0.20 | 0.66 |  | 0.09 | 0.01 | 0.10 |  | **0.37** | 0.02 | 0.39 |  | 0.14 | 0.00 | 0.14 |
| Strip_cultivar | **0.17** | 0.14 | 0.31 |  | -0.10 | -0.05 | -0.15 |  | **0.58** | 0.02 | 0.60 |  | 0.04 | 0.00 | 0.04 |
| Strip_additive | **0.43** | 0.19 | 0.62 |  | 0.06 | 0.01 | 0.07 |  | **0.49** | 0.04 | 0.53 |  | 0.11 | 0.00 | 0.11 |
| Strip_diversity | 0.19 | 0.36 | 0.55 |  | 0.14 | 0.12 | 0.26 |  | **0.64** | 0.04 | 0.68 |  | 0.04 | 0.00 | 0.04 |

|  | Early herbivore abundance (log) | | |  | Early herbivore richness | | |  | Late herbivore abundance (log) | | |  | Late herbivore richness | | |
| --- | --- | --- | --- | --- | --- | --- | --- | --- | --- | --- | --- | --- | --- | --- | --- |
| Cropping system | Direct | Indirect | Total |  | Direct | Indirect | Total |  | Direct | Indirect | Total |  | Direct | Indirect | Total |
| All | -0.06 | -0.01 | -0.07 |  | **-0.04** | 0.06 | 0.02 |  | -0.06 | 0.00 | -0.06 |  | **0.12** | 0.00 | 0.12 |
| Monoculture | 0.00 | 0.01 | 0.01 |  | -0.01 | 0.05 | 0.04 |  | -0.07 | 0.00 | -0.07 |  | 0.06 | 0.00 | 0.06 |
| Strip | 0.00 | -0.01 | -0.01 |  | -0.06 | 0.04 | -0.02 |  | 0.08 | -0.01 | 0.07 |  | 0.02 | 0.00 | 0.02 |
| Strip_cultivar | 0.00 | 0.03 | 0.03 |  | -0.04 | -0.05 | -0.09 |  | **-0.34** | 0.02 | -0.32 |  | **0.33** | 0.00 | 0.33 |
| Strip_additive | 0.00 | -0.09 | -0.09 |  | -0.02 | 0.12 | 0.10 |  | 0.17 | 0.00 | 0.17 |  | 0.03 | 0.00 | 0.03 |
| Strip_diversity | 0.00 | -0.01 | -0.01 |  | 0.00 | 0.00 | 0.00 |  | 0.01 | 0.00 | 0.01 |  | 0.14 | 0.00 | 0.14 |

**Table S24.** Effects of early and late season plant size and herbivore abundance and richness on *proportion damaged head weight*, per cropping design. Both direct and indirect effects of early and late plant size variables are shown. Significant direct effects are indicated in bold (α = 0.05). For indirect or total effects it was not possible to assess significance.

|  | Early plant diameter | | |  | Early number of leaves | | |  | Fresh weight wrapper leaves (log) | | |
| --- | --- | --- | --- | --- | --- | --- | --- | --- | --- | --- | --- |
| Cropping system | Direct | Indirect | Total |  | Direct | Indirect | Total |  | Direct | Indirect | Total |
| All | 0.00 | 0.03 | 0.03 |  | 0.00 | -0.01 | -0.01 |  | **0.09** | 0.01 | 0.10 |
| Monoculture | 0.00 | -0.09 | -0.09 |  | 0.00 | 0.02 | 0.02 |  | -0.09 | -0.01 | -0.10 |
| Strip | 0.00 | 0.05 | 0.05 |  | 0.00 | 0.01 | 0.01 |  | 0.08 | -0.01 | 0.07 |
| Strip_cultivar | 0.00 | -0.04 | -0.04 |  | 0.00 | -0.03 | -0.03 |  | 0.06 | 0.04 | 0.10 |
| Strip_additive | 0.00 | -0.04 | -0.04 |  | 0.00 | 0.00 | 0.00 |  | 0.00 | 0.00 | 0.00 |
| Strip_diversity | 0.00 | 0.05 | 0.05 |  | 0.00 | 0.05 | 0.05 |  | 0.22 | -0.02 | 0.20 |

|  | Early herbivore abundance (log) | | |  | Early herbivore richness | | |  | Late herbivore abundance (log) | | |  | Late herbivore richness | | |
| --- | --- | --- | --- | --- | --- | --- | --- | --- | --- | --- | --- | --- | --- | --- | --- |
| Cropping system | Direct | Indirect | Total |  | Direct | Indirect | Total |  | Direct | Indirect | Total |  | Direct | Indirect | Total |
| All | -0.06 | -0.01 | -0.07 |  | 0.00 | 0.02 | 0.02 |  | 0.07 | 0.00 | 0.07 |  | 0.00 | 0.00 | 0.00 |
| Monoculture | 0.01 | 0.00 | 0.01 |  | 0.00 | -0.02 | -0.02 |  | -0.02 | 0.00 | -0.02 |  | 0.00 | 0.00 | 0.00 |
| Strip | -0.01 | 0.00 | -0.01 |  | 0.00 | 0.00 | 0.00 |  | -0.06 | 0.00 | -0.06 |  | 0.00 | 0.00 | 0.00 |
| Strip_cultivar | -0.08 | -0.09 | -0.17 |  | 0.00 | 0.07 | 0.07 |  | **0.41** | 0.00 | 0.41 |  | 0.00 | 0.00 | 0.00 |
| Strip_additive | -0.08 | 0.00 | -0.08 |  | 0.00 | 0.00 | 0.00 |  | 0.00 | 0.00 | 0.00 |  | 0.00 | 0.00 | 0.00 |
| Strip_diversity | -0.18 | 0.00 | -0.18 |  | 0.00 | 0.01 | 0.01 |  | -0.09 | 0.00 | -0.09 |  | 0.00 | 0.00 | 0.00 |

**Table S25.** Effects of early and late season plant size, abundances of herbivore classes, and cabbage head damage on *fresh weight of the cabbage head*, per cropping design. Both direct and indirect effects of early and late plant size variables are shown. Significant direct effects are indicated in bold (α = 0.05). For indirect or total effects it was not possible to assess significance.

|  | Early plant diameter | | |  | Early number of leaves | | |  | Fresh weight wrapper leaves (log) | | |  | Proportion damaged head weight | | |
| --- | --- | --- | --- | --- | --- | --- | --- | --- | --- | --- | --- | --- | --- | --- | --- |
| Cropping system | Direct | Indirect | Total |  | Direct | Indirect | Total |  | Direct | Indirect | Total |  | Direct | Indirect | Total |
| All | **0.22** | 0.28 | 0.50 |  | 0.04 | -0.03 | 0.01 |  | **0.58** | -0.01 | 0.57 |  | **0.06** | 0.00 | 0.06 |
| Monoculture | 0.05 | 0.52 | 0.57 |  | **0.14** | -0.08 | 0.06 |  | **0.56** | 0.00 | 0.56 |  | -0.01 | 0.00 | -0.01 |
| Strip | **0.48** | 0.20 | 0.68 |  | 0.08 | -0.01 | 0.07 |  | **0.38** | 0.02 | 0.40 |  | **0.12** | 0.00 | 0.12 |
| Strip_cultivar | **0.22** | 0.07 | 0.29 |  | -0.11 | -0.07 | -0.18 |  | **0.57** | -0.02 | 0.55 |  | 0.01 | 0.00 | 0.01 |
| Strip_additive | **0.44** | 0.24 | 0.68 |  | 0.07 | -0.03 | 0.04 |  | **0.53** | -0.01 | 0.52 |  | **0.11** | 0.00 | 0.11 |
| Strip_diversity | 0.17 | 0.38 | 0.55 |  | 0.18 | 0.11 | 0.29 |  | **0.60** | 0.06 | 0.66 |  | 0.05 | 0.00 | 0.05 |

|  | Early chewer abundance | | |  | Early sucker abundance (log) | | |  | Early thrips abundance | | |  | Early flea beetle abundance | | |
| --- | --- | --- | --- | --- | --- | --- | --- | --- | --- | --- | --- | --- | --- | --- | --- |
| Cropping system | Direct | Indirect | Total |  | Direct | Indirect | Total |  | Direct | Indirect | Total |  | Direct | Indirect | Total |
| All | **-0.13** | -0.01 | -0.14 |  | 0.00 | 0.00 | 0.00 |  | 0.00 | 0.00 | 0.00 |  | 0.00 | 0.05 | 0.05 |
| Monoculture | 0.05 | 0.02 | 0.07 |  | 0.00 | -0.02 | -0.02 |  | 0.00 | 0.00 | 0.00 |  | 0.00 | 0.04 | 0.04 |
| Strip | **-0.19** | 0.02 | -0.17 |  | 0.00 | 0.01 | 0.01 |  | 0.00 | 0.00 | 0.00 |  | 0.00 | 0.04 | 0.04 |
| Strip_cultivar | **-0.20** | -0.06 | -0.26 |  | 0.00 | 0.00 | 0.00 |  | 0.00 | 0.00 | 0.00 |  | 0.00 | 0.08 | 0.08 |
| Strip_additive | -0.10 | 0.05 | -0.05 |  | 0.00 | -0.02 | -0.02 |  | 0.00 | 0.00 | 0.00 |  | 0.00 | 0.06 | 0.06 |
| Strip_diversity | 0.08 | 0.03 | 0.11 |  | 0.00 | 0.00 | 0.00 |  | 0.00 | 0.00 | 0.00 |  | 0.00 | 0.05 | 0.05 |

|  | Late chewer abundance | | |  | Late sucker abundance | | |  | Late thrips abundance | | |  | Late flea beetle abundance | | |
| --- | --- | --- | --- | --- | --- | --- | --- | --- | --- | --- | --- | --- | --- | --- | --- |
| Cropping system | Direct | Indirect | Total |  | Direct | Indirect | Total |  | Direct | Indirect | Total |  | Direct | Indirect | Total |
| All | 0.01 | 0.00 | 0.01 |  | 0.00 | 0.00 | 0.00 |  | **-0.10** | 0.00 | -0.10 |  | 0.03 | 0.00 | 0.03 |
| Monoculture | 0.01 | 0.00 | 0.01 |  | 0.00 | 0.00 | 0.00 |  | **-0.15** | 0.00 | -0.15 |  | **0.10** | 0.00 | 0.10 |
| Strip | 0.08 | 0.00 | 0.08 |  | 0.00 | 0.00 | 0.00 |  | -0.05 | 0.00 | -0.05 |  | 0.02 | 0.00 | 0.02 |
| Strip_cultivar | -0.02 | 0.00 | -0.02 |  | 0.00 | 0.00 | 0.00 |  | **-0.25** | 0.00 | -0.25 |  | 0.07 | 0.00 | 0.07 |
| Strip_additive | **-0.17** | 0.00 | -0.17 |  | 0.00 | 0.00 | 0.00 |  | -0.03 | 0.00 | -0.03 |  | **0.23** | 0.00 | 0.23 |
| Strip_diversity | 0.08 | 0.00 | 0.08 |  | 0.00 | 0.00 | 0.00 |  | 0.03 | 0.00 | 0.03 |  | 0.09 | 0.00 | 0.09 |

**Table S26.** Effects of early and late season plant size and abundances of herbivore classes on *proportion damaged head weight*, per cropping design. Both direct and indirect effects of early and late plant size variables are shown. Significant direct effects are indicated in bold (α = 0.05). For indirect or total effects it was not possible to assess significance.

|  | Early plant diameter | | |  | Early number of leaves | | |  | Fresh weight wrapper leaves (log) | | |
| --- | --- | --- | --- | --- | --- | --- | --- | --- | --- | --- | --- |
| Cropping system | Direct | Indirect | Total |  | Direct | Indirect | Total |  | Direct | Indirect | Total |
| All | 0.00 | -0.07 | -0.07 |  | 0.00 | -0.01 | -0.01 |  | **-0.10** | 0.00 | -0.10 |
| Monoculture | 0.00 | -0.08 | -0.08 |  | 0.00 | 0.01 | 0.01 |  | -0.11 | 0.00 | -0.11 |
| Strip | 0.00 | 0.04 | 0.04 |  | 0.00 | 0.00 | 0.00 |  | 0.08 | 0.00 | 0.08 |
| Strip_cultivar | 0.00 | -0.07 | -0.07 |  | 0.00 | -0.05 | -0.05 |  | 0.09 | 0.00 | 0.09 |
| Strip_additive | 0.00 | -0.02 | -0.02 |  | 0.00 | 0.00 | 0.00 |  | 0.00 | 0.00 | 0.00 |
| Strip_diversity | 0.00 | 0.01 | 0.01 |  | 0.00 | 0.03 | 0.03 |  | 0.17 | 0.00 | 0.17 |

|  | Early chewer abundance | | |  | Early sucker abundance (log) | | |  | Early thrips abundance | | |  | Early flea beetle abundance | | |
| --- | --- | --- | --- | --- | --- | --- | --- | --- | --- | --- | --- | --- | --- | --- | --- |
| Cropping system | Direct | Indirect | Total |  | Direct | Indirect | Total |  | Direct | Indirect | Total |  | Direct | Indirect | Total |
| All | -0.09 | 0.00 | -0.09 |  | 0.00 | 0.00 | 0.00 |  | 0.00 | 0.00 | 0.00 |  | 0.00 | -0.01 | -0.01 |
| Monoculture | 0.05 | -0.01 | 0.04 |  | 0.00 | 0.00 | 0.00 |  | 0.00 | 0.00 | 0.00 |  | 0.00 | -0.01 | -0.01 |
| Strip | -0.03 | 0.00 | -0.03 |  | 0.00 | 0.00 | 0.00 |  | 0.00 | 0.00 | 0.00 |  | 0.00 | 0.01 | 0.01 |
| Strip_cultivar | **-0.32** | -0.02 | -0.34 |  | 0.00 | 0.00 | 0.00 |  | 0.00 | 0.00 | 0.00 |  | 0.00 | 0.01 | 0.01 |
| Strip_additive | -0.06 | 0.00 | -0.06 |  | 0.00 | 0.00 | 0.00 |  | 0.00 | 0.00 | 0.00 |  | 0.00 | 0.00 | 0.00 |
| Strip_diversity | **-0.28** | 0.00 | -0.26 |  | 0.00 | 0.00 | 0.00 |  | 0.00 | 0.00 | 0.00 |  | 0.00 | 0.01 | 0.01 |

|  | Late chewer abundance | | |  | Late sucker abundance | | |  | Late thrips abundance | | |  | Late flea beetle abundance | | |
| --- | --- | --- | --- | --- | --- | --- | --- | --- | --- | --- | --- | --- | --- | --- | --- |
| Cropping system | Direct | Indirect | Total |  | Direct | Indirect | Total |  | Direct | Indirect | Total |  | Direct | Indirect | Total |
| All | 0.00 | 0.00 | 0.00 |  | 0.00 | 0.00 | 0.00 |  | 0.00 | 0.00 | 0.00 |  | 0.00 | 0.00 | 0.00 |
| Monoculture | 0.00 | 0.00 | 0.00 |  | 0.00 | 0.00 | 0.00 |  | 0.00 | 0.00 | 0.00 |  | 0.00 | 0.00 | 0.00 |
| Strip | 0.00 | 0.00 | 0.00 |  | 0.00 | 0.00 | 0.00 |  | 0.00 | 0.00 | 0.00 |  | 0.00 | 0.00 | 0.00 |
| Strip_cultivar | 0.00 | 0.00 | 0.00 |  | 0.00 | 0.00 | 0.00 |  | 0.00 | 0.00 | 0.00 |  | 0.00 | 0.00 | 0.00 |
| Strip_additive | 0.00 | 0.00 | 0.00 |  | 0.00 | 0.00 | 0.00 |  | 0.00 | 0.00 | 0.00 |  | 0.00 | 0.00 | 0.00 |
| Strip_diversity | 0.00 | 0.00 | 0.00 |  | 0.00 | 0.00 | 0.00 |  | 0.00 | 0.00 | 0.00 |  | 0.00 | 0.00 | 0.00 |
